# The distribution of fitness effects varies phylogenetically across animals

**DOI:** 10.1101/2025.05.13.653358

**Authors:** Meixi Lin, Sneha Chakraborty, Carlos Eduardo G. Amorim, Sergio F. Nigenda-Morales, Annabel C. Beichman, Paulina G. Nuñez-Valencia, Jonathan Mah, Jacqueline A. Robinson, Christopher C. Kyriazis, Christian D. Huber, Andrew E. Webb, Sarah D. Kocher, Frederick I. Archer, Andrés Moreno-Estrada, Robert K. Wayne, Kirk E. Lohmueller

**Author notes:** Centre for Genomic Regulation (CRG), The Barcelona Institute of Science and Technology, Barcelona, 08003, Spain; Universitat Pompeu Fabra (UPF), Barcelona, Spain. Deceased (Robert K. Wayne).

## Abstract

1

The distribution of fitness effects (DFE) describes the selection coefficients (*s*) of newly arising mutations and fundamentally influences population genetic processes. However, the extent and mechanisms of DFE variation have not been systematically investigated across species with divergent phylogenetic histories and ecological functions. Here, we inferred the DFE in natural populations of eleven animal (sub)species, including humans, mice, fin whales, vaquitas, wolves, collared flycatchers, pied flycatchers, halictid bees, *Drosophila*, and mosquitoes. We find that the DFE co-varies with phylogeny, where the expected mutation effects are more similar in closely related species (Pagel’s *λ* = 0.84, P = 0.01). Additionally, mammals have a higher proportion of strongly deleterious mutations (22% to 47% in mammals; 0.0% to 5.4% in insects and birds) and a lower proportion of weakly deleterious mutations than insects and birds. Population size is significantly negatively correlated with the expected impact of new deleterious mutations (PGLS_*λ*_, P = 0.03), and the proportion of new beneficial mutations (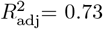, P *<* 0.001). These findings align with Fisher’s Geometric Model (FGM), which defines organismal complexity as the number of phenotypes under selection. Consistent with the FGM’s predictions, we observe that mutations are more deleterious in complex organisms, while beneficial mutations occur more frequently in smaller populations to compensate for the drift load. Our study demonstrates strong phylogenetic constraints in the evolution of a fundamental population genetics parameter, and proposes that, through mechanisms of global epistasis, long-term population size and organismal complexity drive variation in the DFE across animals.

**Significance Statement:** Understanding how mutations affect fitness is fundamental in evolution, but little is known about how and why the distribution of fitness effects (DFE) varies across species. In this study, we examine the DFE in diverse animal populations and show that closely related species exhibit similar patterns of mutation effects, with new mutations being more deleterious in mammals compared with birds and insects. Our findings corroborate Fisher’s Geometric Model, which explains the variation in the DFE across species as a function of organismal complexity and long-term population size. By connecting organismal complexity and population size with the DFE, we offer a phylogenetic view into the selective forces shaping species adaptation and evolution.

## 3 Introduction

Mutations are the raw material on which other evolutionary forces act. The distribution of fitness effects (DFE) represents the relative proportion of new mutations with different selection coefficients (*s*) affecting fitness of the individuals carrying them. In addition to its intrinsic value for understanding how mutations affect fitness, the DFE is a fundamental quantity in evolutionary genetics [1]. The DFE is essential for studying the maintenance of genetic variation, identifying causal variants underlying complex human disease, controlling for background selection in adaptive evolution, and understanding the long-term survival of small populations [1–6]. Consequently, considerable effort has been put into characterizing the biological factors underlying DFE evolution, and estimating the DFE for a given species of interest [1, 7]. However, the patterns of and mechanism behind the variation in the DFE have not been systematically surveyed across many species on a phylogenetic timescale [3].

Several theoretical models have proposed that organismal complexity and long-term population sizes (*N*_*a*_) are pivotal biological factors influencing how the DFE might differ across species. However, while these two factors are common to many models, predictions for how they affect the DFE vary across studies [8–10]. While intuitively comprehensible, precise and comprehensive definitions of complexity remain elusive. For example, complexity can refer to the number of unique cell types and protein-protein interactions [11, 12]. Theoretically, complexity can refer to the number of genetically uncorrelated phenotypic traits under selection, denoting the dimensionality of phenotypic space in Fisher’s Geometric Model (FGM; [13, 14]). We adopt this latter definition throughout this paper. Regardless, how complexity influences the DFE depends on the underlying model and associated assumptions. On the one hand, in complex organisms, highly connected networks confer robustness against the deleterious effects of mutations, therefore, more complex organisms are predicted to have fewer deleterious mutations [9, 15]. Alternatively, increased organismal complexity results in larger effect sizes for random mutations, predicting a higher proportion of deleterious mutations in complex organisms [7, 10, 16].

Similarly, the effect of long-term population size on the DFE is multifaceted. In species with larger effective population sizes, more efficient natural selection favors stable proteins that are then less susceptible to subsequent mutations that disrupt protein functions, thereby diminishing the average deleterious effects of new mutations [8, 17]. Conversely, the reduced selective strength in small populations leads to the accumulation of deleterious mutations, moving the population from the fitness optimum. In turn, an increased proportion of beneficial mutations is expected in species with smaller *N*_*a*_ [10], to compensate for the drift load. Taken together, the distribution of fitness effects at population scale (2*N*_*a*_*s*) could be conserved across species as the interplay between selective strength and protein stability may offset variation in long-term population sizes [8].

Despite the growing number of published DFE in animal species, a lack of standardized DFE comparisons across lineages has hindered testing hypotheses regarding the DFE evolution [3]. In natural populations, the DFE is usually estimated by contrasting allele frequencies, summarized by the site frequency spectrum (SFS), of neutrally evolving and selected variants in genomic data sets [18, 19]. While DFE estimates for model organisms like humans [19, 20], *Drosophila* [18, 21], mice [7, 22], and nematodes [23] have been available, the DFEs for non-model organisms, such as flycatchers [24], great apes [25], Hawaiian monk seals [26], and vaquitas [27], are emerging on a per-lineage basis. These DFE estimates agree that most amino-acid changing mutations are nearly neutral (*−*10 ^*−*5^ *< s* ≤ 0) or weakly deleterious (*−*0.001 *< s* ≤ *−*10^*−*5^), whereas beneficial (*s >* 0) and strongly deleterious (*s* ≤ *−*0.01) mutations are less common [1]. Additionally, the DFE appears to vary across divergent species [24–27]. For example, humans were found to carry a higher proportion of strongly deleterious mutations than *Drosophila* [7, 22]. Interestingly, closely related species demonstrated similar inferred DFE parameters [25, 28]. Unfortunately, the scarcity of high-quality genomic data and genetic parameter estimates (e.g. mutation rates) in non-model organisms [29], coupled with methodological variations and computational inefficiencies in DFE inference [20, 30, 31], have prevented a more comprehensive comparison.

The recent increase in available genomic resources for non-model organisms now offers unprecedented opportunities for comparative population genomics [29]. Leveraging such data, here we infer the DFE in eleven animal (sub)species with diverse phylogenetic relationships and life history traits using *varDFE*. We find strong signals of phylogenetic dependence in the evolution of DFE across species, with mammals bearing more deleterious mutations than insects and birds. To evaluate the biological processes behind the evolution of this fundamental population genetics parameter, we correlate proxies for organismal complexity and long-term population size with the expected mutation effects in different species. Mutations are more deleterious in more complex and smaller populations, suggesting patterns of macroscopic epistasis in DFE evolution.

## 4 Results

### 4.1 Population genetic inference for eleven animal (sub)species

We retrieved high-quality population-level polymorphism datasets for eleven animal (sub)species. These data include three insect species (mosquitoes, *Drosophila*, and halictid bees), two bird species (pied flycatchers and collared flycatchers), and six mammal (sub)species (arctic wolves, gray wolves, vaquitas, fin whales, mice, and humans; Table 1, Dataset S1). The eleven (sub)species we evaluated are distinctive in the potential biological features that affect the DFE, such as phylogenetic position (Fig. 1A), effective population size, and life history traits (Dataset S1). For example, the long-term population sizes (*N*_*a*_) varied from 6264 (vaquitas) to 2.8 *×* 10^6^ (*Drosophila*) individuals (Table 1). All datasets were filtered with the same stringency to retain only high-confidence genotype calls and at least eight diploid high-quality samples in coding regions (Text S1). We tallied the number of variants at different minor allele frequencies and generated folded site frequency spectra (SFS) in synonymous and nonsynonymous/missense regions (SYN-SFS and MIS-SFS, respectively) for each species (Fig. S1, Dataset S2). To implement a robust but flexible workflow to compare the DFE across species, we developed *varDFE* https://github.com/meixilin/varDFE, a python API based on the *∂a∂i* and Fit*∂a∂i* packages [20, 32] (Fig. S2).

**Table 1:** Summary of DFE inference in eleven (sub)species’ datasets, assuming a gamma-distributed DFE for new neutral or deleterious mutations (*s* ≤ 0). “Demography” is the demographic model of choice used for long-term population size inference. “SFS mask” reports if singletons are masked in the site-frequency spectrum during the inference. “*N*_*a*_” is the inferred long-term population size in each dataset. “*E*[|*s*|]” is the expected mutation effects (*s* ≤ 0). “*α*” and “*β*^′^” are the inferred shape and scale parameters respectively in the gamma DFE.

| Common name | Scientific name | Class | Demography | SFS mask | $N_a$ | $E[\vert s\vert ]$ | $\alpha$ | $\beta'$ |
| --- | --- | --- | --- | --- | --- | --- | --- | --- |
| mosquito | *Anopheles coluzzii* | Insecta | three-epoch | TRUE | 2286875 | 0.00056 | 0.30 | 0.0019 |
| fruit fly | *Drosophila melanogaster* | Insecta | two-epoch | FALSE | 2766461 | 0.00014 | 0.36 | 0.00038 |
| halictid bee | *Lasioglossum calceatum* | Insecta | two-epoch | TRUE | 280467 | 0.00051 | 0.30 | 0.0017 |
| pied flycatcher | *Ficedula hypoleuca* | Aves | two-epoch | FALSE | 246918 | 0.00052 | 0.29 | 0.0018 |
| collared flycatcher | *Ficedula albicollis* | Aves | three-epoch | FALSE | 138027 | 0.0021 | 0.22 | 0.0097 |
| gray wolf | *Canis lupus lupus* | Mammalia | three-epoch | FALSE | 45517 | 0.089 | 0.12 | 0.72 |
| arctic wolf | *Canis lupus arctos* | Mammalia | two-epoch | FALSE | 74139 | 0.67 | 0.11 | 6.30 |
| vaquita | *Phocoena sinus* | Mammalia | two-epoch | FALSE | 6264 | 0.013 | 0.13 | 0.096 |
| fin whale | *Balaenoptera physalus* | Mammalia | two-epoch | FALSE | 12411 | 0.027 | 0.14 | 0.19 |
| house mouse | *Mus musculus castaneus* | Mammalia | two-epoch | FALSE | 207682 | 0.010 | 0.21 | 0.048 |
| human | *Homo sapiens* | Mammalia | two-epoch | FALSE | 7041 | 0.014 | 0.19 | 0.073 |

**Figure 1:**
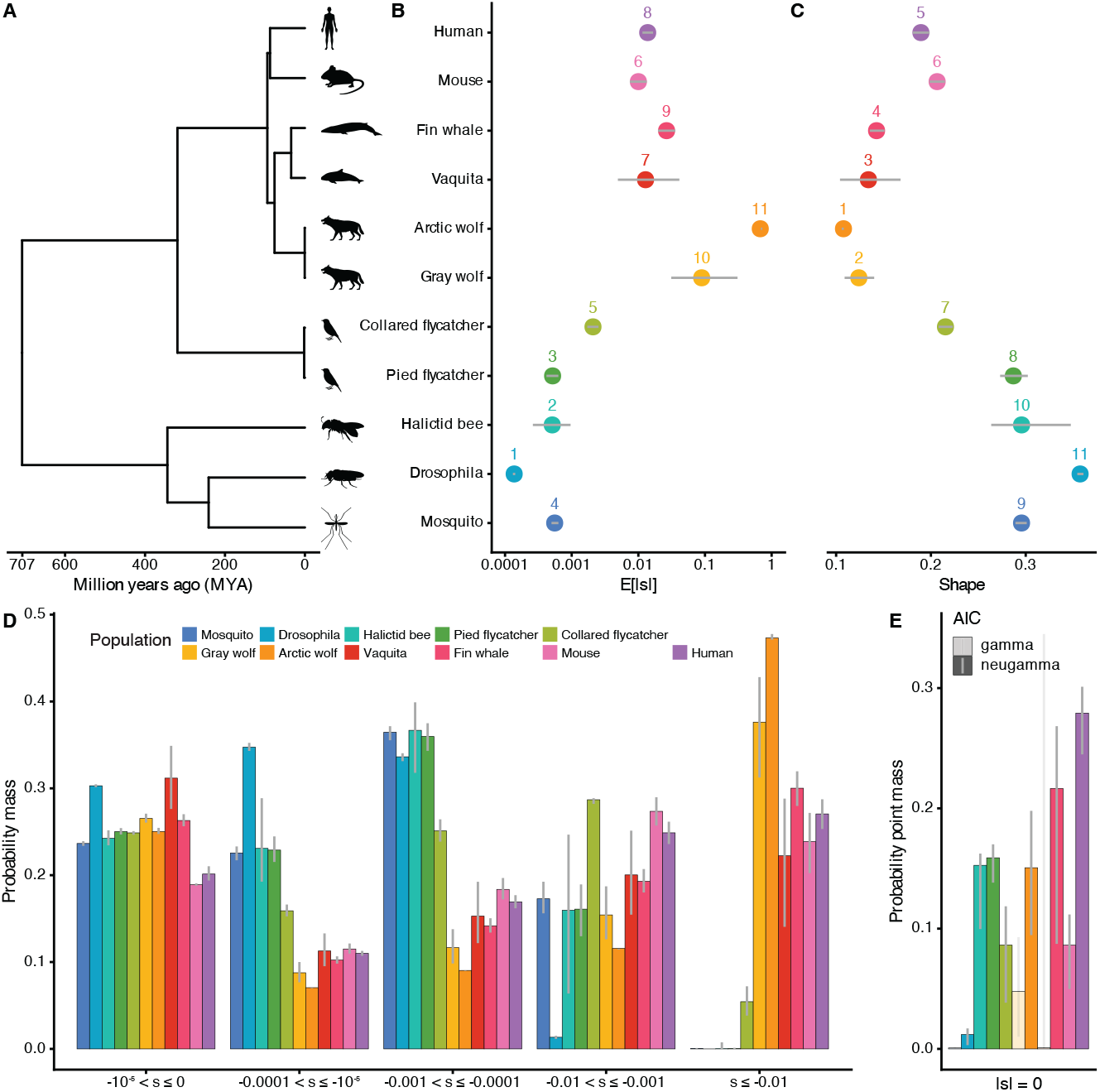
Mutations are more deleterious in mammals compared to insects and birds. (A) The phylogenetic relationship among the (sub)species included in this study (n = 11). (B) Assuming a gamma-distributed DFE of neutral and deleterious mutations, the expected deleterious mutation effects (*E*[|*s*|]) for eleven (sub)species. From top to bottom, species are increasingly divergent from humans: mice, fin whales, vaquitas, arctic wolves, gray wolves, collared flycatchers, pied flycatchers, halictid bees, *Drosophila* and mosquitoes. Numbers show the ranks of *E*[|*s*|]. (C) The shape parameter (*α*) in the gamma-distributed DFE for eleven (sub)species. Numbers show the ranks of *α*. (D) Proportions of mutations in various categories of |*s*|. From left to right, mutations range from (nearly) neutral (−10^−5^ *< s* ≤ 0) to very strongly deleterious (*s <* −0.01). (E) The inferred probability point mass at *s* = 0 (*p*_neu_) assuming a gamma-distributed DFE with a neutral point mass (neugamma DFE). In species where the Akaike Information Criterion (AIC) for the neugamma DFE is higher than that for the gamma DFE, which are mosquitoes, gray wolves and vaquitas, the bars are marked as translucent. (B-E) Gray lines represent 95% confidence intervals derived from 200 Poisson-resampled SFS.

To estimate long-term population size for each species, and control for population history and linked selection in the coding regions for downstream DFE inference [20, 31], we inferred demographic parameters from the putatively neutral SYN-SFS (Fig. S3, Dataset S3). A demographic model with a single change in population size (i.e. a two-epoch model) provided a good fit to SYN-SFS and reasonable parameter estimates for most species except for the mosquitoes, halictid bees, gray wolves, and collared flycatchers. For the gray wolves and collared flycatchers datasets, the three-epoch demographic model (i.e. two changes in population size) performed better with more realistic parameter estimates or drastically improved fit to the SYN-SFS. For the mosquitoes and halictid bees datasets, no demographic model based on the full SYN-SFS fit the data well, and using the singleton-masked SYN-SFS substantially improved model fit (Dataset S4, Text S1). All the inferred demographic models are qualitatively consistent with previous estimates (Dataset S5). We utilized the best-fit demographic parameters from the chosen models (Table 1) for downstream DFE inference.

### 4.2 Mutations are more deleterious in mammals compared with insects and birds

Conditional on the inferred demography, we estimated the DFE for new nonsynonymous mutations for each species, assuming that mutations are neutral or deleterious (*s* ≤ 0) and the DFE follows a gamma distribution [20]. The gamma distribution yielded a robust fit to the observed MIS-SFS across all datasets (Fig. S4, Dataset S6).

On average, mutations are 4.8 to 4923 times more deleterious in mammals (the expected selection coefficient, *E*[|*s*|] is 0.010 in mice to 0.67 in arctic wolves, n = 6), compared to insects (*E*[|*s*|] = 0.00014 in *Drosophila* to 0.00056 in mosquitoes, n = 3) and birds (*E*[|*s*|] = 0.00052 in pied flycatchers and *E*[|*s*|] = 0.0021 in collared flycatchers; Fig. 1B). Noticeably, the scale (*β*^*′*^) parameter for the arctic wolves reached the upper boundary of the parameter space during inference. To rule out potential data artifacts, we analyzed an independently generated whole-genome dataset from Russian Karelia gray wolves [33], which also yield a high *β*^*′*^ value, confirming *E*[|*s*|] is very deleterious in wolves. The shape (*α*) parameter of the gamma distribution differs between species groups as well. On average, mammals have lower estimates for the shape parameter (*α* = 0.11 in arctic wolves to *α* = 0.21 in mice, n = 6) compared to insects and birds (*α* = 0.22 in collared flycatchers to *α* = 0.36 in *Drosophila*, n = 5; Fig. 1C). The extent of variation in *α* that we detected is in line with previous findings [28].

We also observed that mammals have much higher proportions of strongly deleterious mutations than insects and birds (Fig. 1D). The maximum-likelihood gamma distribution for each species demonstrated that 22% (vaquitas) to 47% (arctic wolves) of mutations in mammals are very strongly deleterious (|*s*| ≥ 0.01), compared to 0.0% (*Drosophila*) to 5.4% (collared flycatchers) in insects and birds. On the other hand, insects and birds have a larger proportion of weakly deleterious mutations (10 ^*−*5^ ≤ |*s*| *<* 0.001, 41% in collared flycatchers to 68% in *Drosophila*, n = 5), compared to mammals (16% in arctic wolves to 30% in mice, n = 6). The proportions of neutral to nearly neutral mutations (0 ≤ |*s*| *<* 10^*−*5^) are similar in all eleven (sub)species. To directly estimate the proportion of (nearly) neutral mutations, we considered a different functional form, where DFE followed a mixture of gamma distribution with an additional point mass at neutrality (neugamma DFE [7], eq. 1, Dataset S7). This neugamma DFE improved model fit for most datasets (lower AIC scores) compared to the gamma DFE (Figs. S5, S6), except for mosquitoes, gray wolves, and vaquitas. Similar to the patterns obtained assuming a gamma DFE (Fig. 1D), the inferred proportion of neutral mutations (*p*_neu_) did not exhibit a phylogenetic pattern (Figs. 1E, S7). *p*_neu_ varied from 0.081% (mosquitoes) to 16% (pied flycatchers) in insects and birds (n = 5), and from 0.089% (vaquitas) to 28% (humans) in mammals (n = 6). We also attempted to directly estimate the proportion of lethal mutations (*s* = *−*0.5) [20, 34]. However, our approach likely lacks sufficient power to confidently infer the proportion of lethal mutations, due to poor model fit and unrealistic estimates of the proportion of lethal mutations (Figs. S6-S8, Dataset S8, Text S2).

To evaluate the robustness of our analysis, we tested if different probability distributions for the DFE, demographic models, or decisions to mask singletons in the SFS affected DFE inference (Fig. S9). We examined the gamma, neugamma, and lognormal distributions (Fig. S10, Dataset S9) for the DFE. When assuming the neugamma DFE, *E*[|*s*|] remained more deleterious in mammals compared with insects and birds. When assuming the lognormal DFE, the phylogenetic signal was less evident due to reduced *E*[|*s*|] estimates in mammals (Figs. S9A, S11). However, the AIC for the lognormal DFE is the lowest only in four datasets (mosquitoes, gray wolves, arctic wolves, and vaquitas; Fig. S12), suggesting that the *E*[|*s*|] derived from the lognormal DFE is less supported by the data. The gamma distribution was confirmed as an adequate functional form for DFE comparisons, given its parsimony, consistent second-lowest AIC across most datasets (Fig. S12), and good visual fit to the observed SFS (Fig. S4). Next, we tested the potential impacts of demographic model misspecification and singleton-masking treatments. The average mutation effects and proportion of each mutation category remain unchanged qualitatively (Figs. S9B-C, S13, S14; Datasets S10, S11).

To account for the combinative effects of DFE and demographic model specifications, we generated model-averaged *E*[|*s*|] estimates that weigh outputs from different DFE and demographic models by their AIC (Fig. 2A, Dataset S12). Using the full SFS, the model-averaged *E*[|*s*|] is more deleterious in mammals (*E*[|*s*|] = 0.0034 in humans to 0.28 in arctic wolves, n = 5) compared to insects (*E*[|*s*|] = 2.2 *×* 10^*−*5^ in halictid bees to 3.7 *×* 10^*−*5^ in mosquitoes, n = 3) and birds (*E*[|*s*|] = 0.00019 in pied flycatchers and *E*[|*s*|] = 0.00099 in collared flycatchers), with the exception that model-averaged *E*[|*s*|] estimates in mice is now 0.00078, due to the improved fit of a more complex demographic models with extreme demographic events (Text S3). The proportion of strongly deleterious mutations (|*s*| ≥ 0.01) remains higher in mammals (21% in fin whales to 56% in arctic wolves) except for the mice dataset (0.89%), compared to insects and birds (0% in mosquitoes to 0.95% in *Drosophila*, n = 5, Figs. 2B, S15). We also evaluated the model-averaged *E*[|*s*|] in all datasets using the singleton-masked SFS, because no model fits the full SFS well in the mosquito dataset (*min*(∆*LL*) = 2018). Using singleton-masked SFS, the model-averaged *E*[|*s*|] is more deleterious in mammals and birds (*E*[|*s*|] = 0.0011 in collared flycatchers to 0.06 in arctic wolves, n = 8), compared to insects (*E*[|*s*|] = 3.6 *×* 10^*−*5^ in *Drosophila* to 0.00034 in mosquitoes; Figs. 2A, S15).

**Figure 2:**
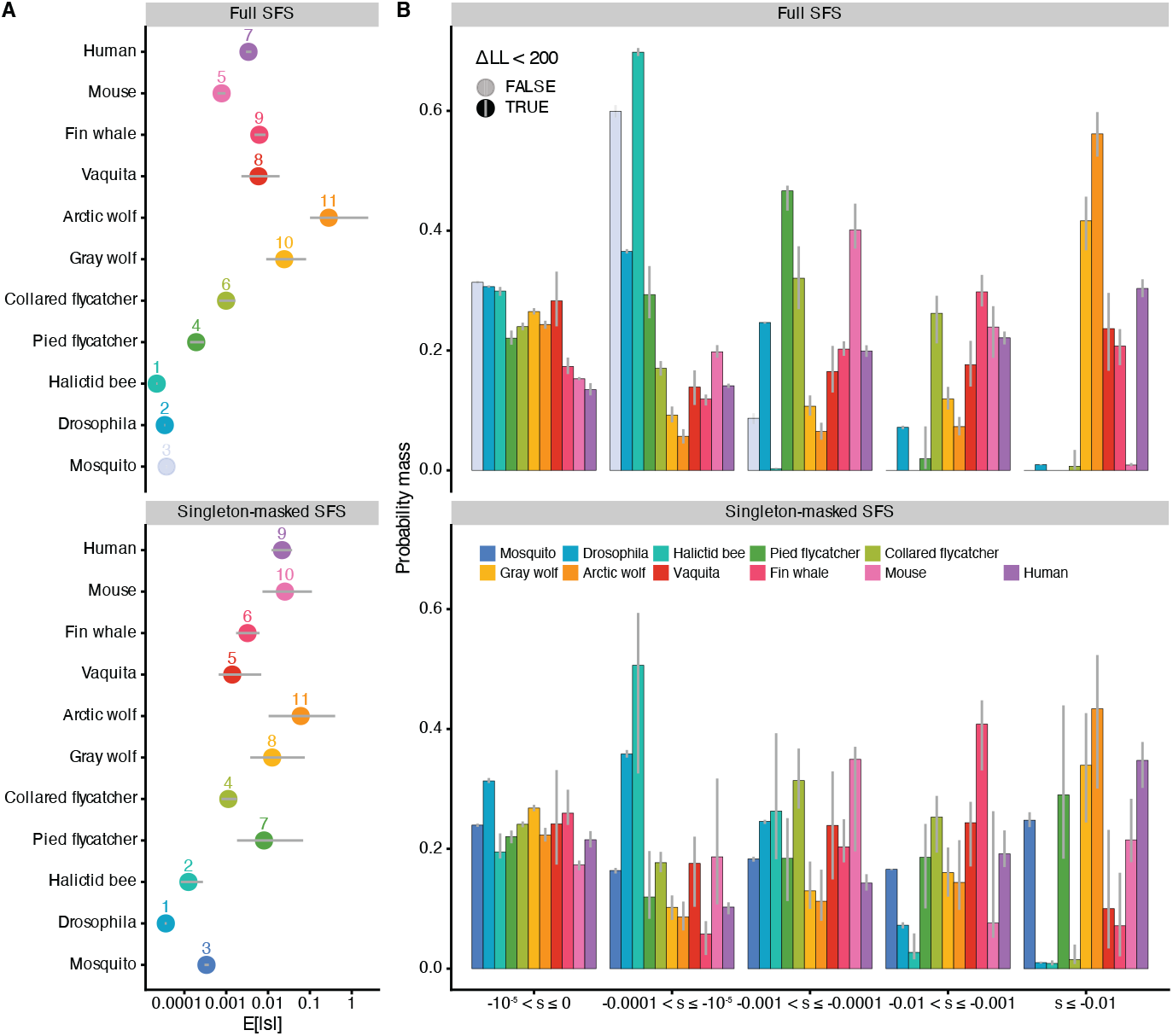
Model-averaged DFE estimates using full SFS or singleton-masked SFS. The estimates are weighted-averages of six DFE and demographic inference runs per species, combining outputs from gamma-, neugamma-, lognormal-distributed DFE given two-epoch or three-epoch demographic models by their AIC. Panels show whether full SFS or singleton-masked SFS is used in the inference. For results corresponding to singleton-masking treatments in Fig. 1, refer to Fig. S15. (A) The model-averaged expected deleterious mutation effects (*E*[|*s*|]) for eleven (sub)species. Numbers show the ranks of *E*[|*s*|]. (B) The model-averaged proportions of mutations in various categories of |*s*|. From left to right, mutations range from (nearly) neutral (−10^−5^ *< s* ≤ 0) to very strongly deleterious (*s <* −0.01). (A-B) Gray lines represent 95% confidence intervals.

To confirm our DFE inference can confidently detect strongly deleterious mutations even in very large populations, we performed a SLiM simulation using demographic parameters inferred from *Drosophila* and DFE parameters inferred from humans. The DFE inferred from the simulation is similar to humans (average *E*[|*s*|] = 0.017, n = 84), indicating our methods can confidently detect strongly deleterious mutations in large populations (Fig. S16). Therefore, the lack of strongly deleterious mutations inferred in species with large population sizes is likely not due to biases in our inference procedure, and instead reflect true differences in the DFE. Overall, our DFE inference proved mostly robust to analytical choices, with some exceptions noted in specific datasets (Text S3).

In summary, the distribution of fitness effects is more similar in closely related species, with mammals harboring more strongly deleterious and less weakly deleterious variations compared with insects and birds.

### 4.3 The DFE co-varies with phylogeny in animals

The apparent co-variation of the DFE with phylogeny motivated us to test for phylogenetic signal more formally. Pagel’s *λ* quantifies how trait evolution depends on the underlying phylogenetic tree across species, where *λ* = 0 suggests no phylogenetic signal and *λ* = 1 indicates strong phylogenetic dependence [35]. Assuming a gamma-distributed DFE, we calculated Pagel’s *λ* for log10 transformed expected selection coefficients [log_10_(*E*[|*s*|])], and the shape parameter (*α*, Fig. 1B-C). For log_10_ (*E*[|*s*|]), Pagel’s *λ* was 0.84 (P = 0.01, likelihood ratio test [LRT]). For the shape parameter, Pagel’s *λ* was 0.86 (P = 0.004, LRT). Since the DFE parameters for arctic wolves reached upper boundaries, Pagel’s *λ* was recalculated excluding the arctic wolves dataset. Strong phylogenetic signal was consistently observed for both metrics (*λ* = 0.86, P = 0.008 for log_10_ (*E*[|*s*|]); *λ* = 0.81, P = 0.01 for *α*). To acount for model uncertainties, we repeated Pagel’s *λ* analysis for the model-averaged *E*[|*s*|] estimates, and strong phylogenetic signal remains (*λ* = 0.80, P = 0.01 using full SFS; *λ* = 0.74, P = 0.02 using singleton-masked SFS). Both log_10_ (*E*[|*s*|]) and *α* suggested a significant tendency that the evolution of DFE is influenced by the shared evolutionary history determined by the phylogenetic structure (Dataset S13).

To formally test whether the DFE varies across species, we examined a null model where the shape and scale parameters of the gamma distribution were constrained to be the same across the eleven (sub)species datasets, while still accounting for their individual demographic histories. The maximum likelihood estimate (MLE) for this null model is found by summing the individual likelihood surfaces of each species’ DFE using a grid-search approach (Fig. 3A). The alternative model allows each species to have its own shape and scale parameters, as described in previous sections (Fig. 3B-C). This alternative model fits the data (MIS-SFS) significantly better than the null model (LRT, Λ = 18203, df = 20, P *<* 10^*−*16^ ; Figs. 3A-C, S17). Therefore, the variation of DFE across taxa is statistically supported by our analyses.

**Figure 3:**
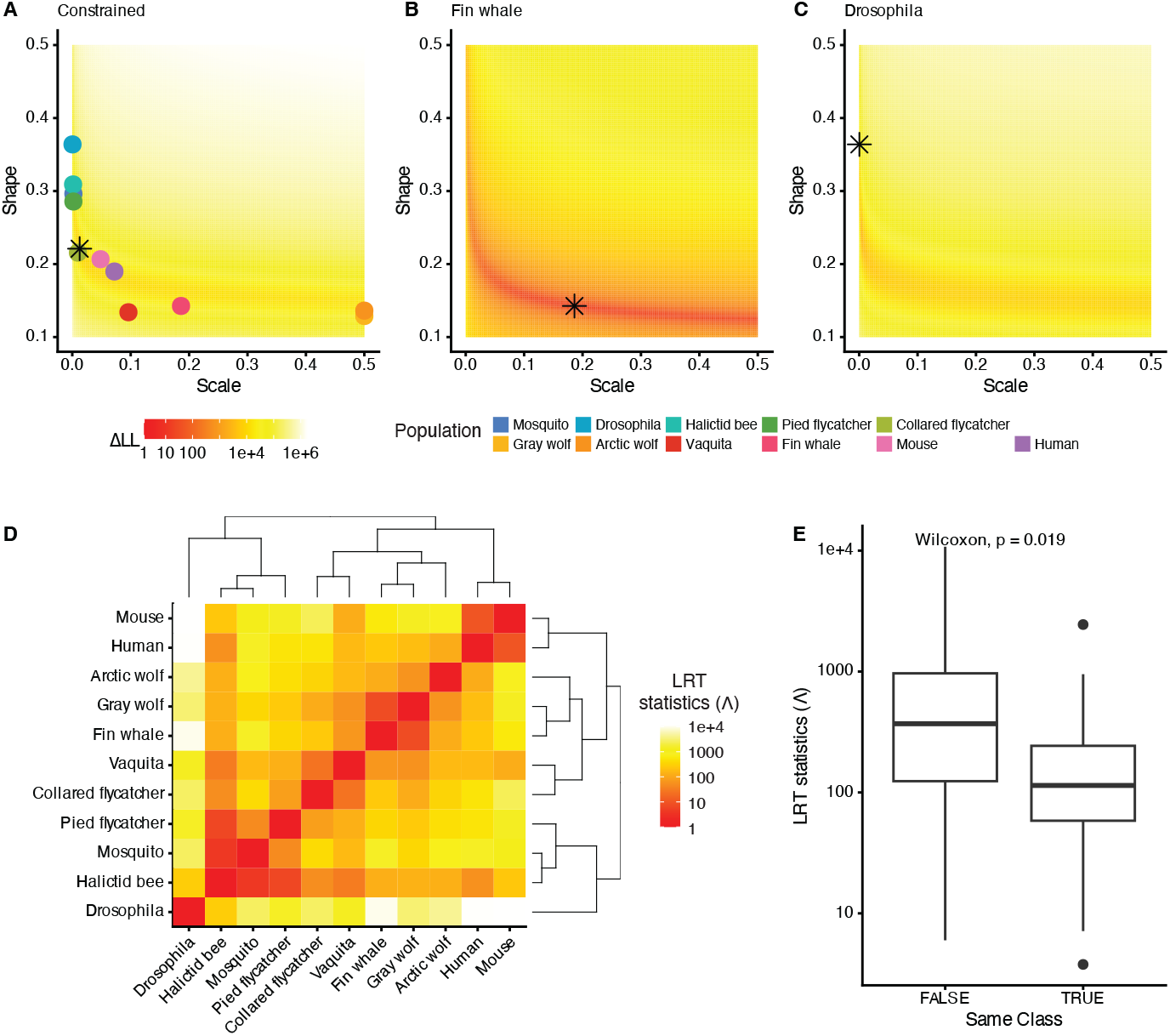
Grid-search-based likelihood-ratio tests confirm the phylogenetic signal in DFE evolution. (A-C) The log-likelihood surfaces for the shape (*α*) and scale (*β*^*′*^) parameters (A) under the null model where all species are constrained to the same parameters or under the alternative model allowing species’ parameters to vary in representative (B) fin whales and (C) *Drosophila* datasets (other species’ log-likelihood surfaces shown in Fig. S17). Background colors from yellow to red indicate the differences in log-likelihood for given parameters to data. On each log-likelihood surface, the maximum likelihood estimate (MLE) derived for (A) the null model or (B-C) the alternative model in respective species is overlaid as the asterisk. In (A), the colored points showing MLEs for each species derived from the alternative model are overlaid for comparison. The two wolves datasets’ MLEs derived from parameter optimization procedures (Fig. 1) exceed the grid-search derived MLE plotted here. For all other species, the MLEs from the grid search and parameter optimization are the same. (D) Pairwise LRT statistics (Λ) for each species pair is colored on a log10 scale and hierarchically clustered. Darker cells represent more similar DFE estimates in the species compared, such as the human-mouse pair, whereas lighter cells represent more distinct DFEs, such as the human-*Drosophila* pair. The dendrogram derived from hierarchical clustering is annotated. (E) The boxplot of pairwise LRT statistics in (D), categorized by whether species pair belong to the same Class. The y-axis is in log10 scale.

To further investigate the relationship between phylogenetic proximity and DFE similarity, we next used the likelihood ratio test framework to compare pairs of species. Here, the null model is relaxed to constrain DFE parameters to be identical only for the two species compared, rather than across all species. The resulting LRT statistic (Λ) serves as a measure of DFE similarity between species pairs, with lower values indicating higher similarity (eq. 2). Although the hierarchical clustering of log10-scaled pairwise LRT statistics [log_10_ (Λ)] did not fully recover the underlying phylogenetic relationships of the eleven (sub)species, insects and mammals clustered into two main lineages, the pied flycatchers fell into the insect lineage while the collared flycatchers clustered with vaquitas (Fig. 3D). In addition, log_10_ (Λ) is significantly lower (P = 0.02, Wilcoxon test) in pairwise comparisons within the same taxonomic class (average log_10_ (Λ) = 2.06 *±* 0.71, n = 19), such as mammal-mammal pairs, compared to inter-class pairwise comparisons (average log_10_ (Λ) = 2.59 *±* 0.76, n = 36), such as mammal-insect pairs (Fig. 3E). In summary, species that are more closely related phylogenetically, tend to have more similar DFEs compared to species that are more distantly related.

### 4.4 Population size and body mass are correlated with the evolution of the DFE

We next explored the relationship between the DFE and candidate life history traits related to long-term population size and organismal complexity, using the phylogenetic generalized least squares model (PGLS_*λ*_ [36]). PGLS_*λ*_ estimates the amount of phylogenetic signal (*λ*) in the regression residuals, and incorporates *λ* in regression parameter estimates, to control for interspecific autocorrelation due to phylogeny. As the long-term population size decreased (Fig. 4A), on average, mutations became more deleterious in a population (log_10_(*E*[|*s*|]) ∼ log_10_ (*N*_*a*_), PGLS_*λ*_, slope (b) = -0.83, *λ* = 0, P = 0.03). Although significant correlation exists between *N*_*a*_ and *E*[|*s*|] the log-likelihood surface of *λ* had another local optimum at *λ* = 0.78 (*LL*_*λ*=0.78_ = *−*13.24, *LL*_*λ*=0_ = *−*13.10), questioning the amount of phylogenetic signal in the correlation (Fig. S18). On the other hand, body mass was positively correlated with *E*[|*s*|] (Fig. 4D, PGLS_*λ*_, b = 0.26, *λ* = 0, P = 0.002). The two wolf (sub)species’ DFE deviated the most from the regression, likely due to their very deleterious estimated DFEs. Other parameters we tested, including the generation time, age at maturity, maximum longevity, and mutation rates, were not significantly correlated with *E*[|*s*|] after considering phylogenetic dependence (Fig. S19). To account for model uncertainty, we repeated PGLS_*λ*_ analysis for the model-averaged *E*[|*s*|] estimates. Significant correlation remains between *N*_*a*_ and *E*[|*s*|] (b = -0.98, *λ* = 0, P = 0.02 using full SFS; b = -0.88, *λ* = 0, P = 0.02 using singleton-masked SFS; Fig. 4B-C), body mass and *E*[|*s*|] (b = 0.31, *λ* = 0, P = 9 *×* 10^*−*4^ using full SFS; b = 0.20, *λ* = 0, P = 0.02 using singleton-masked SFS; Figs. 4E-F, S19, S20). We repeated the PGLS_*λ*_ analyses excluding the arctic wolves outlier and for the shape (*α*) parameter (Text S4). Long-term population size and body mass remains correlated with both *E*[|*s*|] and *α* regardless of outlier treatments (Fig. S21, Dataset S14).

**Figure 4:**
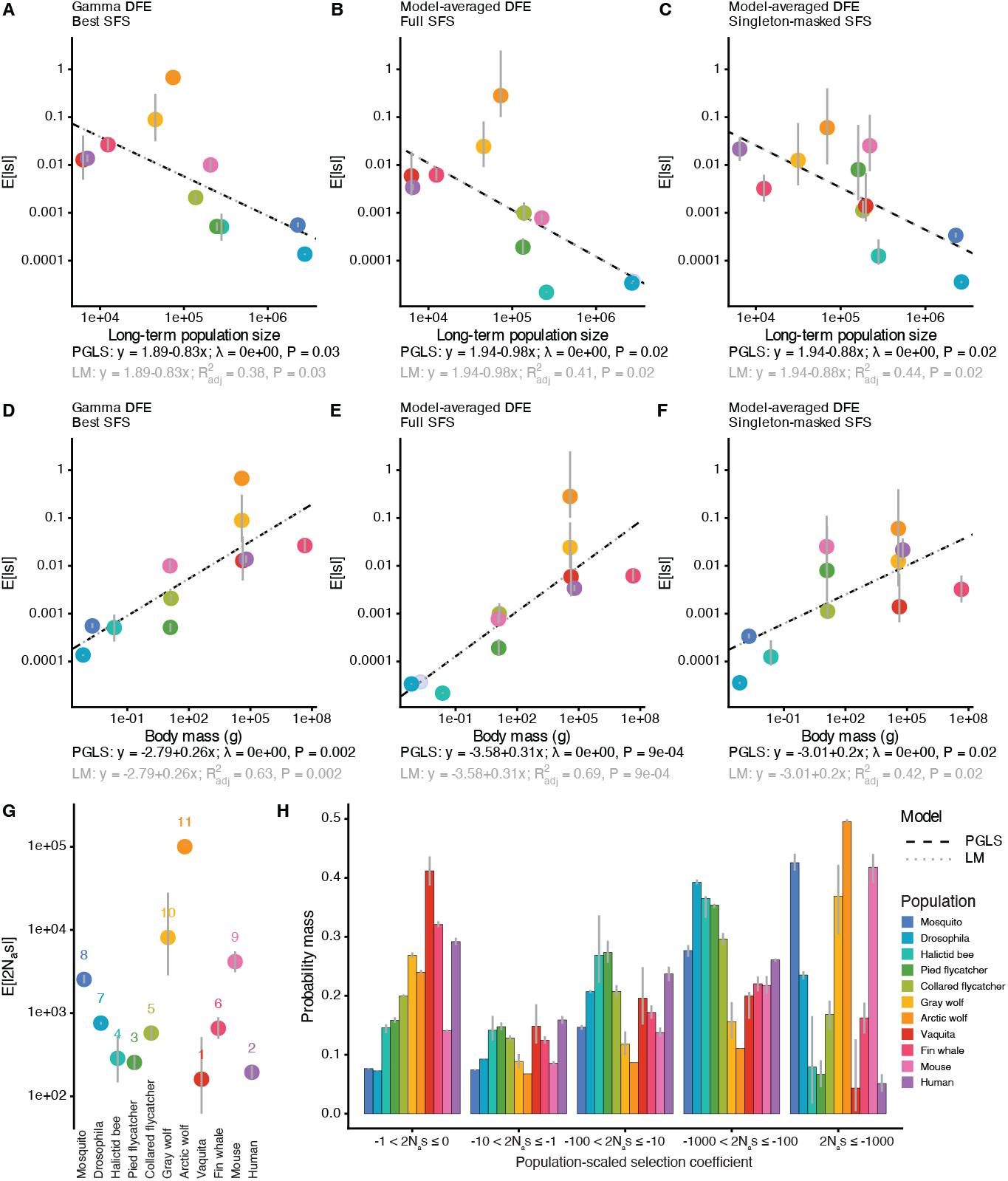
Long-term population size (*N*_*a*_) and body mass are correlated with the expected mutation effects (*E*[|*s*|]), but *N*_*a*_ does not explain all the variations in the DFE across species. The phylogenetic generalized least square model with simultaneously inferred phylogenetic signal (PGLS_*λ*_) was fitted for *E*[|*s*|] with (A C) long-term population size, (D-F) body mass, and other life history traits (Figs. S19, S20). *E*[|*s*|] were computed from (A,D) gamma DFE, (B,E) model-averaged DFE using full SFS or (C,F) model-average DFE using singleton-masked SFS. The PGLS_*λ*_ results are overlaid as a dashed black line, with equations, inferred phylogenetic signal (*λ*), and likelihood-ratio derived p-value annotated at the bottom. The linear regression results are overlaid as dotted gray line, with equations and p-values annotated at the bottom. (G) Assuming a gamma-distributed DFE, the expected population-scaled deleterious mutation effects (*E*[|2*N*_*a*_*s*|]) are not conserved across eleven species. Numbers show the ranks of *E*[|2*N*_*a*_*s*|]. (A-G) Each point represents one species, with gray vertical lines representing confidence intervals. All axes are in log10 scale. (H) Proportions of mutations with various ranges of |2*N*_*a*_*s*|. From left to right, mutations ranged from (nearly) neutral (−1 *<* 2*N*_*a*_*s* ≤ 0) to very strongly deleterious (2*N*_*a*_*s* ≤ −1000). Gray lines represent 95% confidence intervals.

The negative correlation of population size and expected mutation effects at the individual level suggests potential constancy for population-scaled mutation effects (*γ* = 2*N*_*a*_*s*), an important prediction on the DFE when considering protein thermostability [8]. Therefore, we next sought to test if population size by itself can explain the variation in the DFE observed across species by examining the population-scaled DFE. Assuming the population-scaled DFE follows a gamma distribution, the expected mutation effects (*E*[|2*N*_*a*_*s*|]) still vary across species (Fig. 4G-H). Compared to the co-variation of *E*[|*s*|] with phylogeny, we no longer observe similar *E*[|2*N*_*a*_*s*|] within lineages. The expected population-scaled mutation effects are the highest in two wolves populations (*E*[|2*N*_*a*_*s*|]= 1.0 *×* 10^5^ in arctic wolves and 8120 in gray wolves), and the lowest in the vaquitas (161) and humans (195; Fig. 4G). The proportion of each type of mutation fluctuated for each species as well (Fig. 4H). Adopting the grid-search analysis for population-scaled DFE comparisons, we found that the alternative model where each species has its own population-scaled shape and scale parameters fit the data significantly better than the null model assuming a constant *γ* in different species (LRT, Λ = 99508, df = 20, P *<* 10^*−*16^ ; Fig. S22). When comparing the population-scaled DFE between species pairs, log_10_ (Λ) for *γ* remained significantly lower (P = 0.01, Wilcoxon test) in pairwise comparisons within the same taxonomic class (log_10_ (Λ) = 2.98 *±* 0.62, n = 19) compared to inter-class comparisons (3.47 *±* 0.75, n = 36; Fig. S23), as observed in the log_10_ (Λ) statistics for *s* (Fig. 3E). Therefore, although population size is strongly correlated with the expected selection coefficient (Fig. 4A), it is not the sole predictor for the variation in the DFE across species (Figs. 4G-H, S22, S23).

### 4.5 Fisher’s geometric model enables concurrent examination of population size and organismal complexity’s influence on the DFE

Previous work has suggested that Fisher’s geometric model (FGM) can explain patterns of DFE variation across divergent species. FGM models the fitness landscape with a minimal set of parameters to study the emergent properties of mutational effects [14]. In FGM, the fitness of an organism is characterized by a multi-dimensional Gaussian decay function with a local optimum in *n*-dimension phenotype space. Mutations create new phenotypes, modeled as random vectors with effect scale *σ* that act within a subset of *m* phenotype dimensions (*m* ≤ *n*) [10]. The phenotype dimensionality *n* is termed “organismal complexity”, and both *m* and *σ* could be a direct measurement of organismal complexity from different perspectives [7, 16]. When the population is at the fitness optimum and mutations are universally pleiotropic (*m* = *n*), the derived DFE is a gamma distribution with only neutral or deleterious mutations (*s* ≤ 0) [16]. When the population is under mutation-selection-drift equilibrium, the increase in fitness from beneficial mutations should counteract the drift load, i.e. the decrease in fitness caused by fixed deleterious mutations, and the population has an equilibrium phenotypic distance (*z*_eq_; *z*_eq_ *>* 0) to the fitness optimum. The formula for this FGM-derived DFE, which models the full DFE spectrum (*−*0.5 ≤ *s* ≤ 0.5), is a function of three key parameters in FGM (eq. 3, derived in [10]): long-term population size (*N*_*a*_), mutation pleiotropy (*m*), and scale of mutation effects (*σ*).

We inferred the parameters for this FGM-derived DFE for eleven animal (sub)species (eq. 3). The FGM-derived DFE provided an equally good or better fit to the MIS-SFS compared to the gamma DFE (Fig. S24, Dataset S15). In addition, the inferred long-term population sizes from the FGM-derived DFE largely agree with the estimates of *N*_*a*_ derived from demographic inference using genetic variation data (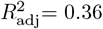, P = 0.03 linear regression; Fig. 5A). Both suggest that most populations are likely under mutation-selection-drift equilibrium and the FGM-based DFE describes their mutation effects well. The vaquita population’s *N*_*a*_ estimate deviated the most between the two methods, likely reflective of its long-term small population size and recent bottleneck [27], thus departing from mutation-selection-drift equilibrium.

**Figure 5:**
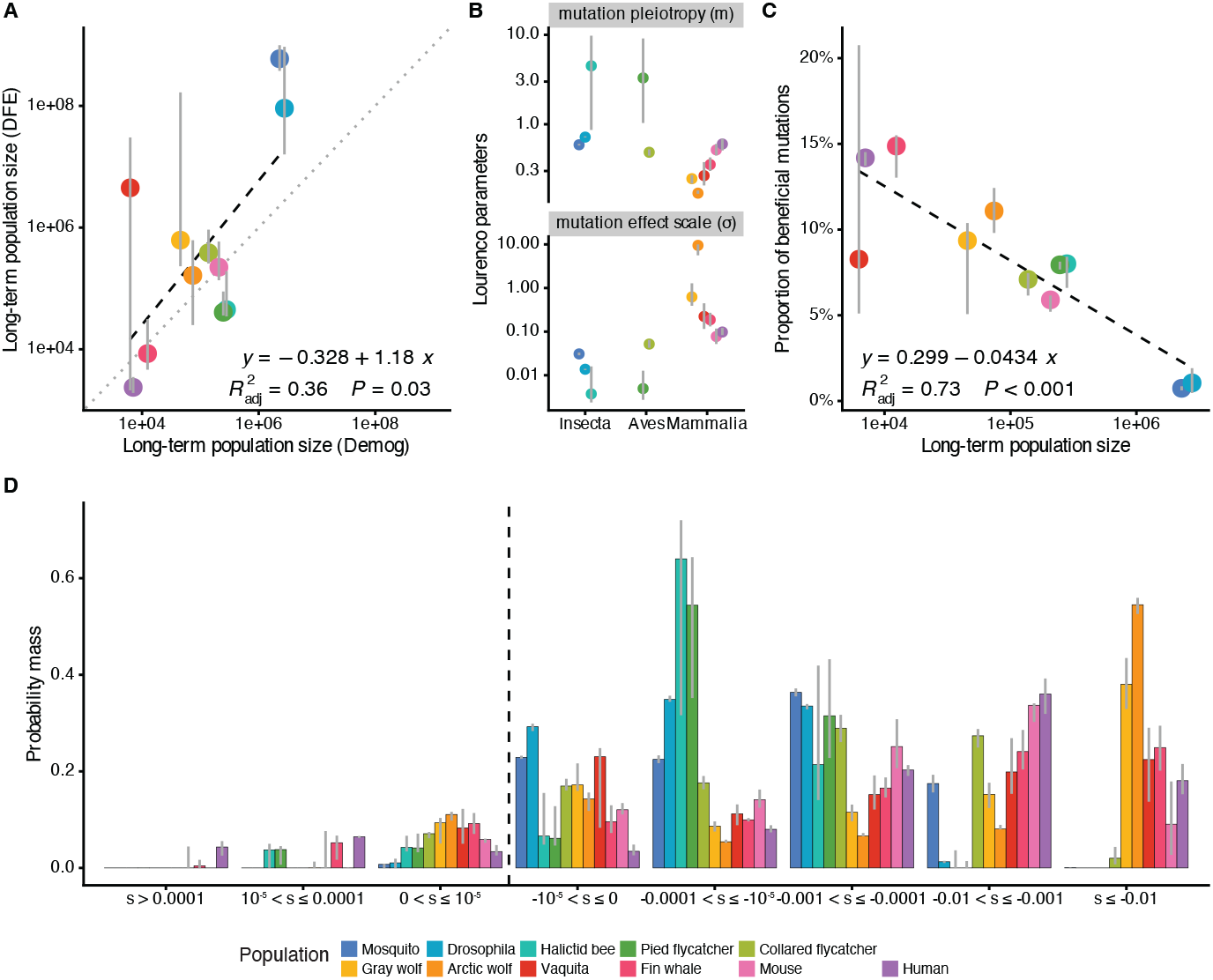
Fitting a Fisher’s Geometric Model (FGM) derived DFE. (A) The inferred long-term effective population size (*N*_*a*_) from the FGM-derived DFE (y-axis) is correlated with the effective size inferred from genetic variation data using *∂a∂i* (x-axis). Axes are on a log10 scale. (B) The estimated scale of mutation effects (*σ*) and mutation pleiotropy (*m*) parameters for eleven (sub)species, grouped by the Class of each species. Note that *σ* is higher in mammals compared to insects. The y-axis is on a log10 scale. (C) The estimated proportion of beneficial mutations (*s >* 0) is negatively correlated with long-term population size inferred from *∂a∂i*. The x-axis is on a log10 scale. (D) Proportions of mutations in various categories of *s* estimated from the FGM-derived DFE for each species. From left to right, mutations range from beneficial (*s >* 0), (nearly) neutral (−10^−5^ *< s* ≤ 0) to very strongly deleterious (*s* ≤ −0.01). Dashed line marks *s* = 0. Gray lines represent 95% confidence intervals.

We found some support that the mutation effect scale (*σ*) is reflective of a species’ phylogenetic position. Mammals have larger *σ* (0.077 to 9.5, n = 6) compared with insects and birds (0.0037 to 0.052, n = 5; Fig. 5B). The mutation pleiotropy (*m*) is negatively correlated with the inferred *σ* (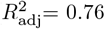, P *<* 0.001, linear regression; Fig. S25), suggesting potential auto-correlations during inference. Although less conserved within lineages, mammals tend to have smaller *m* (0.17 to 0.60, n = 6) compared with insects and birds (0.49 to 4.5, n = 5). However, Pagel’s *λ* did not suggest phylogenetic signal for either *σ* (*λ* = 0, P = 1) or *m* (*λ* = 0.51, P = 0.67; Dataset S13).

Next we examined the proportion of mutations with various ranges of *s* predicted from the FGM-derived DFE (Fig. 5D). Similar to the findings assuming the gamma DFE, strongly deleterious mutations are scarce in insects and birds populations (0.0% to 2.0% in collared flycatchers), but much more abundant in mammals (9.1% in mice to 55% in arctic wolves). Additionally, FGM predicts that as the population size decreases, the proportion of beneficial mutations increases to counteract the increased drift load [10]. Indeed, assuming FGM, smaller populations were found to harbor more beneficial mutations (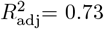, P *<* 0.001, linear regression; Fig. 5C).

## 5 Discussion

Here, we evaluated the long-standing question of the evolutionary stability of the DFE across species, taking advantage of the advances in whole genome sequencing of non-model organisms. To do this, we curated high-quality whole-genome based polymorphism data with a large sample size per species. Our results suggest that the DFE evolution is contingent upon the underlying phylogenetic relationships across species, with closely-related species sharing a more similar DFE (Fig. 3). Overall, mutations are more deleterious in mammals, compared to insects and birds (Figs. 1, 2). We also observed strong correlations between the DFE with long-term population size and organismal complexity (Figs. 4, 5).

The DFE estimates we obtained align with those of previous studies (Dataset S16). In humans, the inferred gamma distribution shape parameter in our study (0.19 *±* 0.0048), falls within the range of previous studies: from 0.12 ∼ 0.16 [28], 0.16 [25], 0.18 ∼ 0.21 [19], 0.17 ∼ 0.21 [20], to 0.2 [18]. In *Drosophila*, our inferred shape parameter of 0.36 *±* 0.0017 aligns with previous estimates: from 0.35 [18], to 0.32 ∼ 0.41 [28]. Even in non-model organisms, our DFE inferences are concordant with published studies [22, 24, 27, 28]. The consistency of our DFE estimates with the literature further confirmed that the DFE variation observed across species is likely of biological relevance. Notably, we found an unusually high proportion of deleterious mutations in the DFE in two wolves species (Fig. 1B). As we currently lack a satisfying explanation for the distinct DFE in canids, it is worthy of further investigation [37]. Similar to prior research, the DFEs inferred in this study are restricted to nonsynonymous mutations in coding regions. We also assume all mutations are additive and future studies are needed to jointly infer the DFE with dominance [38–42], and to explore the DFE variations for non-coding and structural variants [43–45].

We found significant phylogenetic correlations during the evolution of the DFE in animals, (Pagel’s *λ* = 0.84, Fig. 1B), and the pairwise likelihood ratio test showed that the DFE is more similar in species-pairs within the same taxonomic class (Fig. 3E). Several studies have suggested that the DFE is similar in closely related species, but this phylogenetic signal has not been systematically evaluated until now. For example, high levels of DFE correlation were reported between populations within species, in humans, *Drosophila* and wild tomato [46], and between species within lineages, such as in great apes [25] and *Populus* trees [47]. In addition, stark DFE differences have been consistently observed between species with early divergences, such as humans and *Drosophila* [7, 18]. Previous work comparing the DFE across diverse groups of animals had reported among-taxa variation [28, 48], similar to our findings. Chen et al. [28] acknowledged the differences in DFE between invertebrates and vertebrates, whereas the uncertainties for *E*[|*s*|] in their study prevented further investigations. Although Galtier and Rousselle [48] considered the strong group effect on the shape parameter to be a methodological artifact and deemed the DFE to be the same across species, it has later been shown that their model’s inclusion of lethal mutations flattened the variations in entries of different SFSs, and was unsuitable for fitting a single gamma DFE to multi-species data [49]. Recently, between-species, but not within-species, variation in the DFE was reported for six forest tree species, corroborating the phylogenetic constraints we found in the animal DFEs [50]. Emergent studies in comparative population genomics suggest that phylogenetic correlations exist not only in mutation’s fitness effects, as demonstrated here, but also in other fundamental population genetics parameters, such as mutation rates and mutational spectra in vertebrates [51, 52]. Although our study did not find correlations of mutation rate with *E*[|*s*|] after correcting for phylogenetic signal (Fig. S19), simulations have suggested a complex joint distribution of selection coefficients and mutation rates [53]. Future studies could leverage the broad phylogenetic context provided by macrogenetics to distinguish the co-evolution of mutation rates and their effects, and to address core evolutionary hypotheses, such as adaptive evolution and Lewontin’s paradox [22, 29, 54].

Fisher’s geometric model offers a simplistic yet versatile genotype-phenotype-fitness landscape to explain the mechanism underlying phylogenetic signals in the DFE variation. In FGM, an organism’s phenotype is abstracted into a point in an *n*-dimensional space. Mutations move the phenotype with its associated fitness, which decays with the distance to the phenotypic optimum [14]. Solutions for an organism’s DFE at mutation-selection-drift equilibrium can be derived under this model (eq. 3) and fit the empirical data well (Fig. S24), suggesting that FGM’s fitness landscape is appropriate for describing the DFE across species. Additionally, our data aligned with two key predictions from the FGM-derived equilibrium DFE regarding long-term population size (*N*_*a*_) and organismal complexity (*n*). First, as *N*_*a*_ decreases, a species’ phenotype at mutation-selection-drift equilibrium is farther away from its fitness optimum. The variance of the DFE increases, leading to the increased proportions of both beneficial and strongly deleterious mutations in smaller populations (Fig. S26). Indeed, we observed that the proportion of beneficial mutations increased in smaller populations (Fig. 5C). Moreover, mammals with lower *N*_*a*_ exhibited a markedly higher proportion of highly deleterious mutations compared to insects and birds. This trend is consistent across both the full DFE derived from the FGM (Fig. 5D), and the deleterious component of the DFE modeled by the gamma distribution (Fig. 1D). Note that *N*_*a*_ by itself does not explain all the variation in the DFE across species, because we did not observe a constant *E*[|2*N*_*a*_*s*|] across populations (Fig. 4G-H). Second, the expected effect of mutations (*E*[*s*]) becomes more deleterious as organismal complexity increases, because mutations are more likely to disrupt something important when more dimensions are available in the phenotype space [7]. While biological complexity lacks a straightforward definition, and the FGM’s complexity parameter *n* (the dimensionality in the phenotype space) is not directly measurable, evidence suggests that mammals are more complex than insects. This is indicated by their greater number of unique cell types [55], larger and more complex genomes [56], and more extensive protein-protein interactions and transcriptional regulatory networks [57]. Although we cannot directly estimate *n*, the mutation effect scale (*σ*), which is correlated with *n*, can be inferred from the FGM-derived DFE [10, 16]. *σ* is also larger in mammals compared to insects and birds (Fig. 5B). Indeed, we discovered that mutations are on average more deleterious in the more complex mammal species, compared with insects (Fig. 1).

Beyond Fisher’s geometric model, our findings that the DFE co-varies with ancestral population size and complexity can be interpreted under the broader concept of macroscopic or global epistasis. In other words, a mutation’s fitness effect depends on the population’s background fitness [58, 59]. Theoretically, a stable mutation-selection-drift equilibrium requires that the selective coefficients depend on the background fitness. To maintain stable fitness at equilibrium, the distribution of *s* needs to broaden as fitness declines [60]. With a log-concave fitness function, *w*(*z*) = *exp*(*−z*^2^), where *z* is the distance to phenotype optimum, the FGM has a built-in mechanism of epistasis [7, 14]. This leads to the prediction that the DFE variance increases as the equilibrium distance *z* increases in species with smaller ancestral population size and higher complexity. However, macroscopic epistasis is not exclusive to FGM, and can be observed from a neutral genotype network model [61] as well as first principles of population genetics. Natural selection is more efficient in larger populations than in smaller populations. As such, in larger populations, positive selection will be more effective at fixing mutations that increase protein stability, function, and robustness (Fig. 6A). When new deleterious mutations occur in large populations, these deleterious mutations occur in the optimal background and may have only weak effects. These are the mutations whose fitness effects have been measured in our present analysis. In smaller populations, however, selection will be unable to fine-tune the proteome (Fig. 6B), resulting in unstable proteins. When subsequent deleterious mutations occur in smaller populations, they occur in a less optimal background, and as such, they are more likely to be strongly deleterious. Evidence for global epistasis comes not only from DFEs derived from genetic variation data, as this study shows, but also from experimental findings. For example, the observation that larger populations have a lower proportion of beneficial mutations (Fig. 5C), is consistent with the widely observed diminishing-return epistasis in experimental microbial evolution, where the strength of beneficial mutations quickly declines as populations gain fitness in laboratory settings [62, 63]. Overall, the correlation in *E*[|*s*|] with long-term population size is analogous to the drift-barrier hypothesis on mutation rate variation across large phylogenetic scales [64]. Our current results suggest that a similar drift-barrier exists for *E*[|*s*|] (Fig. 6).

**Figure 6:**
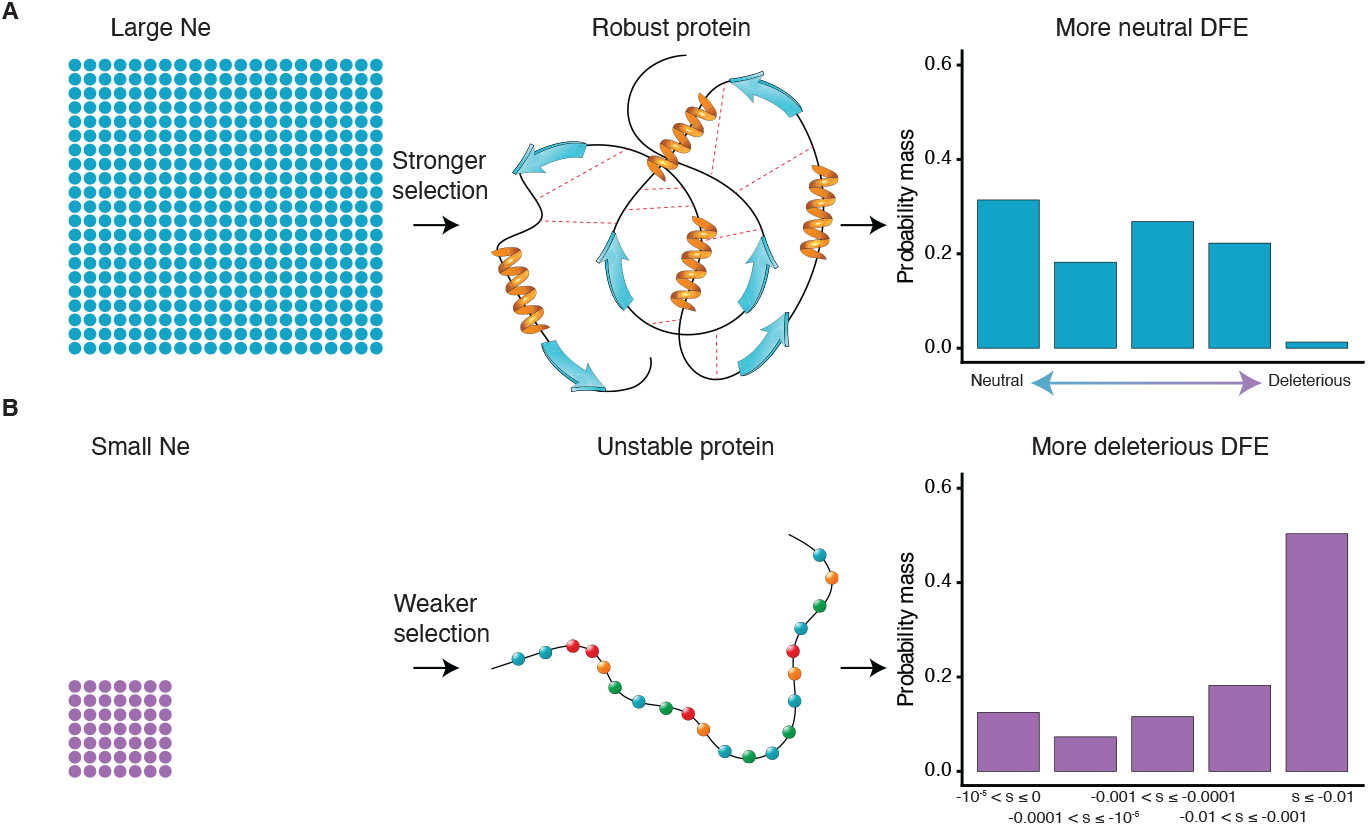
The drift-barrier of DFE evolution. (A) In large populations, efficient natural selection results in robust proteins. Because proteins are robust, subsequent deleterious mutations, whose effects we measured in our study, are only weakly deleterious. The inferred DFE becomes more neutral. (B) In smaller populations, natural selection is less effective at optimizing proteins. Thus, subsequent deleterious mutations, whose effects we measured in our study, are more strongly deleterious. The inferred DFE becomes more deleterious. Proportions of mutations in various categories of |*s*| is generated assuming a gamma-distributed DFE, with the same population-scaled shape and scale parameters of *α* = 0.2, *β* = 5000, but different *N*_*a*_ of 10^6^ in (A) and 10000 in (B).

Our findings on the phylogenetic dependence of DFE variation have several practical implications. In conservation biology, genomics-informed simulations are increasingly popular to estimate the genetic health and quantify the deleterious load of small and fragmented populations [27, 65–67]. The relative conservation of the DFE parameters within closely-related lineages suggests that it is reasonable to assume DFE parameters from closely-related species when the study species’ DFE parameter is not available. However, we caution against using constant DFE parameters across divergent lineages (e.g. mammals vs insects), as significant variation in the DFE exists. Our study also provides additional insights for adaptive evolution, by offering a phylogenetic constraint in the constant evolutionary processes of genetic drift and purifying selection, allowing for a more appropriate evolutionary null model for identifying adaptive variants [4].

Overall, we speculate that while *N*_*a*_ and its changes are similar in closely-related species due to shared evolutionary histories, they are not inherently phylogenetically constrained. Long-term population size and demographic events significantly influence a population’s DFE by modifying the balance between selection and drift, therefore altering the fitness background. Conversely, organismal complexity reflects the physiological constraints imposed by phylogenetic structure, and probably governs the DFE at the mutation-phenotype mapping level through molecular mechanisms, such as protein stability and functional redundancy. Notably, we observed that mutations tend to be more deleterious in more complex mammal species (Fig. 1), contradicting Kimura’s classical proposition that more mutations are neutral in more complex species [15]. In the future, we propose extending Fisher’s geometric model from a single-species to a multi-species model where organismal complexity across species is regulated by a hyperparameter that adjusts the degree of phylogenetic dependence. In conclusion, our study demonstrated strong phylogenetic correlations in the DFE variation across animals. Through mechanisms of global epistasis, long-term population size and organismal complexity are likely underlying the phylogenetic dependence of the DFE.

## 6 Methods

### 6.1 Data sets and site frequency spectra

Polymorphism datasets in eleven animal (sub)species were compiled from various whole genome resequencing projects using a standardized curation workflow designed to retain only high-quality variants and samples for DFE inference (Text S1). Most datasets had pre-annotated variant effects. If not, we annotated the variant call files (VCF) using default options in snpEff (v.5.1 [68]). To retain only high-quality variants, we selected genomic sites that: 1. passed the filters set in the original dataset, 2. were in canonical coding region (CDS) but not in soft-masked, unknown or CpG islands of the genome, 3. were monomorphic or biallelic single nucleotide polymorphisms (SNP) that are annotated as “synonymous” or “nonsynonymous/missense”. To retain only high-quality samples, we selected at least eight diploid samples for each species that: 1. had at least 8x average sequencing depth, 2. belonged to one natural population with no population substructure or admixture identified by principal component analysis (PCA), 3. were not related to other samples identified by kinship analysis.

To summarize polymorphism data, the obtained VCFs were further filtered to exclude genomic sites with more than 20% missing or 75% heterozygous genotypes. We projected down the sample size and computed folded synonymous and nonsynonymous/missense site frequency spectra (SYN-SFS and MIS-SFS, respectively) for each species. Computing a folded SFS mitigates uncertainties in ancestral state classifications. The projected sample size was calculated using the hypergeometric probability distribution to maximize the number of SNPs available and account for the sporadic missing genotypes (Dataset S1). The total synonymous and nonsynonymous sequence lengths (*L*_SYN_ and *L*_MIS_) were obtained using previous estimates of nonsynonymous to synonymous mutation rate ratio 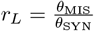.

To further understand the mechanisms of DFE evolution, we also compiled several life history traits for each species (sources and citations in Dataset S1).

### 6.2 Demographic inference

To control for how population size changes influence the SFS, we first inferred demographic parameters from the SYN-SFS using the `Demog1D_sizechangeFIM` module in *varDFE* (v.0.1.0, Fig. S2, based on *∂a∂i* v.2.1.1 [20, 32] and python v.3.10.2). For each species, we fitted four models, including the standard neutral, two-epoch, three-epoch, and four-epoch models. To account for the uncertainties in genotype calls, we repeated the demographic inference by masking the singleton entries in the SYN-SFS. In total, eight demographic inference runs (four demographic models in SYN-SFS with or without masking singletons) for each species were conducted. All settings were the same across datasets except for the starting parameter positions, which were set based on knowledge from prior demographic inference (Datasets S1, S3). For each run, the best-fit parameters with the maximum multinomial log-likelihood in 100 replicates were chosen, and their uncertainties were estimated through Fisher’s Information Matrix (FIM).

To choose the best-fit demographic model and singleton-masking treatment out of the eight runs per (sub)species for downstream DFE inference, we performed a step-wise likelihood-based model selection procedure (Fig. S27, Dataset S4, Text S1). At the start of model selection, the two-epoch demographic model based on full SYN-SFS run was chosen for all datasets. However, if it resulted in poor model fit, we chose a more complex demographic model or masked the singletons in the SYN-SFS. We calculated the population-scaled synonymous mutation rate *θ*_SYN_, and ancestral population size using 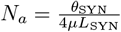 in the chosen demographic run for downstream analyses (Table 1).

### 6.3 DFE inference

Conditional on the inferred demographic scenarios, we estimated the DFE for new nonsynonymous mutations using methods from Fit*∂a∂i* [20]. Briefly, Fit*∂a∂i* takes advantage of the pattern that deleterious mutations are less likely to segregate in the sample of individuals and those that are segregating are more likely to be at lower frequency compared to neutral mutations. Assuming the MIS-SFS is under selection while the SYN-SFS is putatively neutral, the best DFE parameters should fit the differences between MIS-SFS and SYN-SFS in each minor allele frequency bin. We first computed and stored the expected MIS-SFS for a range of population-scaled selection coefficients *γ* = 2*N*_*a*_*s* (−10000 ≤ *γ* ≤ -10^*−*5^ and 10 ^*−*5^ ≤ *γ* ≤ 100) in each species, given their best fit demographic scenarios, using the `DFE1D_refspectra` module in *varDFE*.

To infer the deleterious-only DFE, we assumed that the DFE follows a gamma distribution, given its strong theoretical and empirical support [7, 16, 19]. We parameterized the DFE for each species as *γ* ∼ *Gamma*(*α, β*), where *α* (*α >* 0; shape) and *β* (*β >* 0; scale) are parameters to be inferred. For each species, 100 replicates were run from a permuted starting parameter of *α* = 0.2 and *β* = 4000 using the DFE1D_inferenceFIM module in *varDFE* [7]. The best-fit parameters with the maximum Poisson log-likelihood were chosen, and the corresponding Akaike information criterion (AIC) was calculated. To quantify uncertainty the parameter estimates, we Poisson resampled the MIS-SFS 200 times, and repeated the DFE inference to obtain the 95% confidence interval [19, 20]. Recall that we optimized the gamma distribution parameters for the population-scaled selection coefficient *γ*. To obtain the distribution of *s*, we unscaled the gamma distribution by 1*/*(2*N*_*a*_), therefore, *s* ∼ *Gamma*(*α, β*^′^), where *β* ^′^ = *β/*2*N*_*a*_ and *N*_*a*_ had been inferred from demography estimation. The expected selection coefficient *E*[|*s*|] for each species was calculated given the gamma distribution’s property that *E*[|*s*|] = *αβ*^′^. Similarly, *E*[|2*N*_*a*_*s*|] = *αβ*. To compute the proportion of mutations with different values of *s* or 2*N*_*a*_*s*, we found the cumulative probability in the given range using the `pgamma` function in R (v.3.6.2 [69]).

To estimate the proportion of effectively neutral mutations, we added a point mass at neutrality (*p*_neu_ ≥ 0) to the original gamma distribution. The DFE function can be written as *γ* ∼ *Neugamma*(*α, β, p*_neu_).

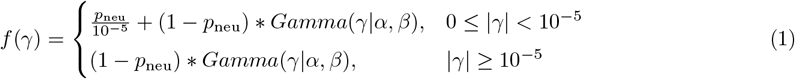

The distribution of *s* is *s* ∼ *Neugamma*(*α, β*^′^, *p*_neu_), where *β* ^′^ = *β/*(2*N*_*a*_). The expected selection coefficient *E*[|*s*|] for each species is 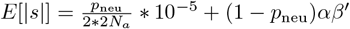.

To examine the robustness of DFE inference, we conducted additional analyses examining alternative functional forms of the DFE, evaluating demographic model impacts, and assessing the effects of singleton masking. For the lognormal distribution, we parameterized the DFE for each species as *γ* ∼ *Lognormal*(*µ*_*s*_, *σ*^2^). The distribution of *s* was unscaled as 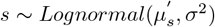, where 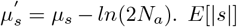 for each species was calculated using 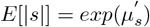. To account for uncertainty in the functional form of the DFE, we performed DFE inference assuming three different functional forms (gamma, neugamma, and lognormal) under both two-epoch and three-epoch demographic models, with and without singleton masking. As masking singleton changes the data log-likelihood, for each species, we generated separate model-averaged DFE estimates based on either the singleton-masked SFS or the full SFS. The model-averaged DFE estimates were obtained by computing weighted-average in six combinations of DFE and demographic models with weights determined by respective AIC values, as described in [25].

### 6.4 Phylogenetic signal detection

To evaluate whether the DFE co-varies with phylogeny, we calculated Pagel’s *λ* for the log10 transformed expected selection coefficient (log_10_ (*E*[|*s*|])) and the shape parameter (*α*) in the gamma-distributed DFE (Dataset S13) using the `phytools` package (v.1.2.0 [70]). The underlying phylogenetic structure and divergence times for eleven (sub)species were compiled from the TimeTree database (http://timetree.org/). To test if the candidate life history traits (*X*) are correlated with the observed DFE parameters (*Y* is *E*[|*s*|] or *α*), and correct for phylogenetic dependency when appropriate, we utilized the PGLS_*λ*_ method in `phytools`. The regression was specified as log_10_(*E*[|*s*|]) ∼ log_10_(*X*) or *α* ∼ log_10_ (*X*) to normalize the data. The *λ* estimated in PGLS_*λ*_ does not equate to the Pagel’s *λ* estimated individually for *X* or *Y*, but represents whether the correlation of *X* and *Y* can be accounted for by phylogenetic dependence in residual error. Candidate explanatory variables (*X*) include long-term population size (*N*_*a*_), body mass in grams, generation time in years per generation, age at maturity in days, maximum longevity in years and mutation rates in mutations per bp per generation. To control for outliers and model uncertainties, we repeated the PGLS_*λ*_ analysis excluding the arctic wolves data point, and on the model-averaged *E*[|*s*|] estimates (Dataset S14).

To formally test for variation in the DFE across species, we used a grid-search approach (`DFE1D_gridsearch` in *varDFE*) to compare each species’ DFE estimates to a null model where the DFE was constrained to be the same across species by likelihood-ratio tests (LRT). 2000 grid points were evenly spaced in biologically meaningful ranges for shape (*α* = 0.1 to 0.5) and scale (*β* ^*′*^ = 0.0001 to 0.5) parameters. Using `integrate` in *∂a∂i* we obtained the expected SFS for each *α−β*^′^ pair and calculated the Poisson log-likelihood relative to empirical MIS-SFS in each species. For the null model, we summed log-likelihoods across species to find the MLE. To test whether the shape (*α*) and scale (*β*^′^) are different in any *x* number of species, the LRT was constructed as Λ = *−*2 ∗ *ln*(*L*_0_*/L*_1_) = *−*2 ∗ (*LL*_0_ *− LL*_1_), where *LL*_0_ is the log-likelihood for the null model, with two DFE parameters inferred 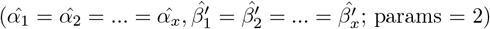 given the inferred demographic parameters for each species (Θ_*D*,1_, Θ_*D*,2_, …, Θ_*D,x*_), and *LL*_1_ is the log-likelihood for the alternate model, where each species is allowed to have its own DFE parameters 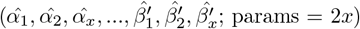. Across the eleven (sub)species tested, asymptotically, Λ should follow a *χ*^2^ distribution with *df* = 2*x −* 2 = 20.

We also calculated the pairwise LRT statistics in all possible pairs of species (*x* = 2). For example, the LRT statistics comparing humans (population 1) and *Drosophila* (population 2) DFE can be written as follows:

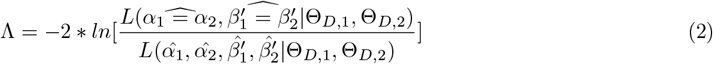

Hierarchical clustering for log_10_ (Λ) was performed by the hclust function in R. To test the variations of *γ* = 2*N*_*a*_*s* across species, we repeated the LRT by evenly spacing 2000 grid points in shape (*α* = 0.1 to 0.5) and population-scaled scale (*β* = 100 to 2.5 *×* 10^4^) parameters.

### 6.5 Fitting the FGM-derived DFE

To investigate the genetic mechanisms underlying DFE variation and to incorporate beneficial mutations, we implemented a functional form of the DFE directly derived from the Fisher’s geometric model [10]. The FGM proposes that fitness (*w*) can be described as a Gaussian function of the distance to the optimum (*z*), *w*(*z*) = *exp*(*−z*^2^), in an *n*-dimension phenotypic space. Random mutations likely affect a subset of *m* phenotypes (*m* ≤ *n*), with fitness effect size *r* following a zero mean Gaussian distribution with scale *σ*. The DFE for a well-adapted population can be described using the mutation pleiotropy (*m*), scale of mutation effects (*σ*) and long-term population size (*N*_*a*_), *γ* ∼ *Lourenco*(*m, σ, N*_*a*_). Here Γ(.) is the gamma function and *K*(.) is the modified Bessel function of the second kind [10].

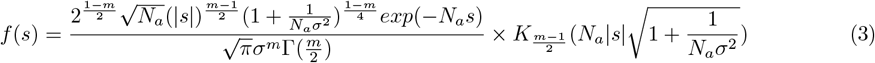

While complex in its expression, this FGM-derived distribution is functionally equivalent to the previously described gamma and lognormal distributions for generating mutational effects. Therefore, we estimated its parameters (*m, σ*, and *N*_*a*_) following methods described in the DFE inference section, through maximum-likelihood optimization by fitting the observed differences between MIS-SFS and SYN-SFS across minor allele frequency bins. To compute the proportion of mutations with different values of *s*, we found the cumulative probability for *s* in given ranges using the `scipy.integrate.quad` function (v.1.8.0) in python. The proportion of beneficial mutations was obtained by calculating the cumulative probability for *s >* 0. We performed two linear regressions using the `lm` function in R: first, log_10_ (*N*_*a*_) inferred in this FGM-derived DFE and the log_10_ (*N*_*a*_) obtained in demographic inference; second, the proportion of beneficial mutations and log_10_ (*N*_*a*_). To test for phylogenetic signal in the inferred *m* and *σ* parameters, we calculated Pagel’s *λ* (Dataset S13).

### 6.6 Simulating human DFE with drosophila demography

To investigate whether Fit*∂a∂i* can accurately detect strongly deleterious mutations in species with large population sizes, we performed a simulation study using Drosophila demography and human DFE. We used the stdpopsim library to obtain coding sequence (CDS) annotations for *Drosophila melanogaster* chromosomes 2L, 2R, 3L, and 3R from the FlyBase BDGP6.32.51 CDS [71]. CDS regions were labeled as “exon”, and non-annotated regions as “bkgd”. To simulate realistic genomic regions, we randomly sampled 84 (each 1 Mb) segments from these chromosomes, with probabilities proportional to chromosome lengths. Annotations for each segment were subset and converted to SLiM’s coordinate format. Next, we implemented a two-epoch Drosophila demographic model in which the ancestral population size was set to 2,766,461, and the population expanded to 7,482,090 over 507,689 generations. We used a uniform recombination rate of 2.03806 *×* 10^*−*8^ which is an average of recombination rates for Drosophila chromosomes from the stdpopsim catalog. Mutation rate was set to 1.5 *×* 10^*−*9^ per bp per generation. Deleterious mutations followed a human-like gamma DFE (mean = –0.0276, shape = 0.189) with a nonsynonymous-to-synonymous ratio of 2.85. Background (noncoding) regions were assigned neutral mutations only. From each simulation, we output site frequency counts using a sample size of 100 and performed inference using methods described above. To reduce computational cost, we applied a scaling factor Q = 100, adjusting population size, mutation rate, recombination rate, and selection coefficients accordingly. We then inferred the DFE on each simulated dataset as done for the empirical data.

## 7 Acknowledgments

For the mosquito dataset, we would like to thank the MalariaGen team for providing open access data from Ag1000G Phase 1 release. The authors thank Dr. Jeremy Field for providing the *L. calceatum* samples used in this study. For the vaquita dataset, we thank Dr. Morin for providing early access to the SFS data. We thank the Southwest Fisheries Science Center’s Marine Mammal and Sea Turtle Research Collection for use of archival vaquita tissue samples. We thank F. Gulland, The Marine Mammal Center, and NOAA Fisheries for funding genome resequencing. For the fin whale dataset, we would like to thank Cei Abreu-Goodger for his support and laboratory space during the initial analysis of the data. Samples were collected by the Southwest Fisheries Science Center (California, USA) in accordance with national guidelines and regulations. We thank Y. Zhen, B. Kim, M. I. A. Cavassim, J. Garcia, E. Wade, X. Qiao, and the MOILAB for their helpful discussions. We especially thank Dr. Exposito-Alonso for his support in completing this work.

## 9 Author contributions

ML and KEL conceived the study. ML, CEGA, SN-M, ACB, PN-V, JAR, AEW, SDK, AME, and RKW generated the data. ML, CEGA, ACB, SC and JM performed the analysis of the data. SC and ML performed simulations. JM, CCK and CDH provided scripts for some analyses. FIA collected and contributed the samples and sample information. KEL performed funding acquisition. ML wrote the first draft of the manuscript. ML and KEL revised the manuscript with input from all the authors.

## 10 Data availability

The *varDFE* python package is available in this github repository: https://github.com/meixilin/varDFE. Additional scripts used to perform the data processing and analysis are available in this github repository: https://github.com/meixilin/varDFE_analyses. The raw sequence data for Arctic wolves and halictid bees will be available from NCBI’s Sequence Read Archive database website upon publication. All other raw sequence data used in this analysis is already publicly available. Additional supportive information will be available from Zenodo upon publication.

