## Supplementary Information for "The distribution of fitness effects varies phylogenetically across animals"

[1]Department of Ecology and Evolutionary Biology, University of California, Los Angeles, Los Angeles, California, 90095, USA
[2]Department of Integrative Biology, University of California, Berkeley, Berkeley, California, 94720, USA
[3]Biology Department, California State University, Northridge, California, 91330, USA
[4]Advanced Genomics Unit, National Laboratory of Genomics for Biodiversity (Langebio), Center for Research and Advanced Studies (Cinvestav), Irapuato, Guanajuato, 36824, Mexico
[5]Department of Biological Sciences, California State University San Marcos, San Marcos, California, 92096, USA
[6]Department of Genome Sciences, University of Washington, Seattle, Washington, 98195, USA
[7]Centro de Ciencias Genómicas, Universidad Nacional Autónoma de México (UNAM), Cuernavaca, Morelos, 62210, Mexico
[8]Bioinformatics Interdepartmental Program, University of California, Los Angeles, Los Angeles, California, 90095, USA
[9]Institute for Human Genetics, University of California, San Francisco (UCSF), San Francisco, California, 94143, USA
[10]Conservation Genetics, San Diego Zoo Wildlife Alliance, Escondido, California, 92027, USA
[11]Department of Biology, Pennsylvania State University, University Park, Pennsylvania, 16802, USA
[12]Dept. of Ecology and Evolutionary Biology, Lewis-Sigler Institute for Integrative Genomics, Princeton University, Princeton, New Jersey, 08544, USA
[13]Howard Hughes Medical Institute, Chevy Chase, Maryland, 20815, USA
[14]Marine Mammal and Turtle Division, Southwest Fisheries Science Center, La Jolla, California, 92037, USA
[b]Current address: Centre for Genomic Regulation (CRG), The Barcelona Institute of Science and Technology, Barcelona, 08003, Spain; Universitat Pompeu Fabra (UPF), Barcelona, Spain.
[c]Deceased

May 11, 2025

This PDF file includes:
Text S1 to S4
Figures S1 to S27
Legends for Datasets S1 to S16
SI References
Other supporting materials for this manuscript include the following:
Datasets S1 to S16

### 1 Text S1: Materials and Methods

#### 1.1 Data sets and initial processing

Polymorphism datasets in eleven animal (sub)species were compiled from various whole genome resequencing projects using a standardized curation workflow designed to retain only high-quality variants and samples for DFE inference (Fig. S2). To ensure variant quality and consistency, we began by selecting datasets with genotypes called using the GATK Best Practices Pipeline [1]. Most datasets had pre-annotated variant effects. For datasets without pre-built variant effect databases, which were halictid bees, collared flycatchers, pied flycatchers, and gray wolves, we annotated and predicted the effects of variants using default options in snpEff (v.5.1 [2]). To retain only high-quality variants, we selected genomic sites that: 1. passed the filters set in the original dataset, 2. were in canonical coding region (CDS) but not in soft-masked, unknown, or CpG islands of the genome, 3. were monomorphic or biallelic single nucleotide polymorphisms (SNP) that are annotated as "synonymous" or "nonsynonymous/missense". To retain only high-quality samples, we selected at least eight diploid samples for each species that: 1. had at least 8x average sequencing depth, 2. belonged to one natural population with no population substructure or admixture identified by principal component analysis (PCA), 3. were not related to other samples identified by kinship analysis.

To further understand the mechanisms of DFE evolution, we also compiled several life history traits for each species (sources and citations in Dataset S1).

The datasets for *Drosophila* [3], vaquitas [4], mice [5] and humans [3], had been quality controlled previously using similar methods described below and not described here.

##### 1.1.1 Mosquitoes

For mosquitoes (*Anopheles gambiae coluzzii*), we queried the Ag1000G phase 3 (Ag3) release using the python package `malariagen-data` (v.0.15.0 [6, 7]) on 2021-12-22 with custom python scripts. We limited our search to a single-species *An. coluzzii* sample set (AG1000G-AO, n = 81) which were collected from breeding sites in Luanda, Angola in 2009 [8].

We first obtained coordinates of canonical CDS in autosomes (2R, 2L, 3R, 3L regions) by identifying the longest transcript of each gene. SNP calls were obtained and filtered using the `ag3.snp_call` method with the following criteria: 1. The sites should pass `varian_filter_pass_gamb_colu` filters provided by the Ag3 release; 2. The sites should fall into previously identified canonical CDS regions; 3. The sites should not be in soft-masked or unknown regions of the genome (`a/t/c/g/N` bases in the genome sequence); 4. The sites should be invariant or biallelic. Variants were annotated using the canonical CDS' annotations provided in `ag3.snp_effects`. The most deleterious annotation is retained for each variant. Only `SYNONYMOUS_CODING` (SYN) and `NON_SYNONYMOUS_CODING` (MIS) variants are downloaded as variant call format (VCF) files.

We performed PCA (function `snpgdsPCA`) and kinship analyses (function `snpgdsIBDKING`) for the LD-pruned SYN-VCF using the R package SNPRelate (v.1.16.0 [9]) to evaluate population structure. In total, 69 individuals SNP-only SYN-VCF and MIS-VCF were retained for downstream site frequency spectrum projection, and 12 individuals were excluded due to high kinship (n = 9; kinship > 0.15) or outliers in PCA analyses [n = 3; outliers defined as $\text{abs}(x - \text{median}(x)) > (6 * \text{sd}(x))$ in PCA].

##### 1.1.2 Halictid bees

For halictid bees (*Lasioglossum calceatum*), whole genome sequencing data were generated for 40 diploid females collected from two populations in the UK (north and south, n = 17 and 23 respectively [10]). Reads were mapped to the chromosome-length Lasioglossum calceatum reference genome (LCAL2.1.1 [11]) and variants were called following the GATK Best Practices Guide [1].

A custom snpEff (v.5.1) database for LCAL2.1.1 was built from the gff3 file using default settings. We annotated and predicted the effects of variants using default options in snpEff. The most deleterious annotation was retained for each variant. Custom variant filtration was performed to retain only high-quality samples and genomic sites, with the following criteria: 1. The sites should be within assembled autosomes; 2. The sites should pass GATK-recommended hard filters ( `QD < 2.0 || FS > 60.0 || MQ < 40.0 || MQRankSum < -12.5 || ReadPosRankSum < -8.0 || SOR > 3.0`); 3. The sites should be outside of repeat regions identified by RepeatMasker [12]; 4. The sites should have a sequencing depth of no more than 1200 or no less than 180; 5. The sites should be invariant or biallelic. For the 153,244,615 sites that passed the above filters out of 282,627,975 total sites, we performed genotype-level filtration. Specifically, for each individual, genotypes with a minimum depth of four reads, a maximum depth of 2.5x mean depth or 30 reads (whichever is higher), and a minimum Phred score of six were kept. Genotypes that failed filtration were converted to missing. Individuals (n = 20 out of 40, $n_{\text{south}} = 9$, $n_{\text{north}} = 11$) with less than 20% of missing genotypes after genotype-level filtration were kept for downstream population structure validations.

We investigated population structure by performing PCA and kinship analyses as described above for mosquitoes using all LD-pruned SNPs. The samples collected from north and south populations separated in the PCA and had low kinship overall. We chose the north population for DFE inference. In total, SNP-only SYN-VCF and MIS-VCF for eleven individuals from the north population of halictid bees in the UK were retained for downstream SFS projection.

##### 1.1.3 Pied flycatchers and collared flycatchers

For pied flycatchers (*Ficedula hypoleuca*) and collared flycatchers (*Ficedula albicollis*), we downloaded filtered SNP-only VCF files for four flycatcher species from dryad on 2022-03-24 [13]. This dataset was derived from a study assessing the genomic differentiation landscapes in Ficedula species pairs [14]. In this dataset, hard filter thresholds, repeat region masks, and genotype filters (`minDP = 5, maxDP = 200, minGQ=30`) had been applied [13].

We lifted the scaffold names and positions (`CHR` and `POS` fields) from the provided VCF file to the NCBI RefSeq chromosome coordinates using custom python scripts (FicAlb1.5, `https://ftp.ncbi.nlm.nih.gov/genomes/all/GCF/000/247/815/GCF_000247815.1_FicAlb1.5/GCF_000247815.1_FicAlb1.5_assembly_structure`, accessed 2022-03-26). The lifted `REF` and `ALT` bases were verified using `BioPython` (v.1.79) and discordant sites were removed (0.01%).

We investigated population structure by performing PCA and kinship analyses described above using all LD-pruned SNPs. A custom snpEff (v.5.1) database for FicAlb1.5 was built from the gtf file using default settings. We annotated and predicted the effects of variants using default options in snpEff. The most deleterious annotation was retained for each variant. Only `synonymous_variant` (SYN) and `missense_variant` (MIS) variants in assembled autosomes (chr1 to chr28) were retained. We additionally masked sites (13.2%) that fell into repeat regions identified by WindowMasker (soft-masked bases in

RefSeq genome; [15]) and CpG islands identified by UCSC genome browser [16].

In total, SNP-only SYN-VCF and MIS-VCF for nine individuals from the pied flycatcher population in mainland Sweden [17], were retained for downstream SFS projection, and no individual was excluded. For the collared flycatcher population (n = 95) in Baltic Sea island Öland, Sweden [17], 88 individuals SNP-only SYN-VCF and MIS-VCF were retained for downstream site frequency spectrum (SFS) projection, and seven individuals were excluded due to high kinship (n = 4; kinship > 0.15) or being outliers in the PCA [n = 3; outliers defined as $abs(x - median(x)) > (6 * sd(x))$ in PCA].

###### 1.1.4 Arctic wolves

For artic wolves (*Canis lupus arctos*), whole genome sequencing data were generated for 15 individuals collected from the arctic wolf population in Canada [18, 19] and this study. Variant calling was performed as described in previous studies [18, 19]. Only sites that passed variant quality filtration and did not fall in repeat regions or CpG islands were retained for this study. Variants were annotated using snpEff v.4.3.1, based on the dog reference genome annotation build CanFam3.1.75 available with snpEff installation. We considered sites where all three potential SNPs were annotated as either `synonymous_variant` or `missense_variant` exclusively, and discarded sites with additional types of annotations (e.g., splicing sites and protein-truncating variants, such as stop-gained variants).

We estimated kinship and excluded relatives up to 4th degree (e.g. first cousins) using the KING-robust estimator [20] implemented in PLINK [21]. In total, SNP-only SYN-VCF and MIS-VCF for 14 individuals (n = 14) were retained for downstream SFS projection, and one individual (n = 1) was excluded due to high kinship.

###### 1.1.5 Gray wolves

For gray wolves (*Canis lupus lupus*), we downloaded filtered biallelic SNP-only VCF files for worldwide wolf and dog populations from Dryad on 2022-09-20 [22]. This dataset was derived from a study assessing dog introgression in the Fennoscandian wolf populations [23]. In this dataset, hard variant filters as recommended by the GATK Best practices Alternative protocol 2, in addition to variant depth, missingness and minor allele frequency filters, and genotype level filters (`minGQ = 30`) had been applied.

We validated the population structure reported in [23] by performing PCA and kinship analyses described above using all LD-pruned SNPs. We annotated and predicted the effects of variants using default options in snpEff (v.5.1) based on the pre-built CanFam3.1.99 database (downloaded on 2022-10-10). The most deleterious annotation was retained for each variant. Only `synonymous_variant` and `missense_variant` variants in assembled autosomes (chr1 to chr38) were kept. We also masked sites that fell into repeat and low complexity regions (soft-masked bases in Ensemble genome release 99) and CpG islands identified by UCSC genome browser [16].

The Russian Karelian gray wolf population was selected for downstream SFS projection because of its high sequencing coverage (mean 30x), relatively large sample size (n = 15), clear clustering on the PCA, and historically large population size. In total, SNP-only SYN-VCF and MIS-VCF for thirteen individuals in this population were generated and two individuals were excluded due to high kinship ($kinship > 0.15$).

##### 1.1.6 Fin whales

For fin whales (*Balaenoptera physalus*), we previously obtained VCF files from a historically stable population (n = 30) in the Eastern North Pacifics [24]. Only sites that passed variant quality filtration and did not fall in repeat regions or CpG islands were retained for this study.

Variants were annotated using SIFT4G v.6.0 [25] and snpEff v.4.3.1. The concordance of variant annotations was confirmed across the two software, and the most deleterious annotation derived from SIFT4G was used per SNP.

We performed PCA and kinship analyses using all LD-pruned SNPs to evaluate population structure [24]. In total, biallelic SNP-only SYN-VCF and MIS-VCF for 27 individuals (n = 27) were retained for downstream SFS projection, and three individuals were excluded due to admixture with another population (n = 2) or low genotyping rate (n = 1).

#### 1.2 Calculation of site frequency spectra and sequence lengths

We summarized polymorphism data using the synonymous and nonsynonymous/missense site frequency spectra (SYN-SFS and MIS-SFS, respectively). SFS generation for *Drosophila* [3], vaquitas [4], mice [5] and humans [3], utilized similar methods as described below and in previous publications.

Here we describe the SFS generation procedures for mosquitoes, halictid bees, flycatchers, wolves and fin whales. Each SNP-only SYN-VCF and MIS-VCF file that passed the previously described quality control steps, was further filtered. Genomic sites with more than 20% missing or 75% heterozygous genotypes were excluded using the GATK (v.3.8) `SelectVariants` [26] or bcftools (v.1.9) `filter` functions [27]. We projected down the sample size and computed folded SYN-SFS and MIS-SFS from SYN-VCF and MIS-VCF for each species using a modified `easySFS` module (`https://github.com/isaacovercast/easySFS`) and `make_data_dict_vcf`, `from_data_dict` methods in $\partial a\partial i$ (v.2.1.1) package [28]. Computing a folded SFS mitigates uncertainties in ancestral state classifications. The projected sample size was calculated using the hypergeometric probability distribution to maximize the number of SNPs available and account for the sporadic missing genotypes (Dataset S1).

To calculate the total synonymous and nonsynonymous sequence lengths ($L_{\mathrm{SYN}}$ and $L_{\mathrm{MIS}}$), we first obtained the length of coding regions ($L_{\mathrm{CDS}}$) for each species. For mosquitoes, halictid bees, fin whales, and arctic wolves, variants had been called for all sites, including monomorphic sites, in the genome. We intersected the coordinates of coding regions with the sites that passed filters and had allele counts (`INFO/AC`) no less than the projected SFS sample size in each species' all-sites VCF (e.g. `INFO/AC` $\geq$ `136` for mosquitoes). For flycatchers, because the all-sites VCF is not available, we calculated an approximated $L_{\mathrm{CDS}}$ (11.8Mb) by multiplying the total CDS length (24.6Mb) from gtf file, with the proportion of callable sites (76.8%) reported for the autosomes (Table S1 in [13]), and the proportion of SNPs (62.7%) passed the additional filters applied during dataset QC. For gray wolves, neither all-sites VCF nor the proportion of callable sites were available as provided [23, 29]. Therefore, we utilized the proportion of callable sites reported in the Arctic wolves dataset as an approximate proxy.

For all species, we then calculated the synonymous and nonsynonymous sequence lengths ($L_{\mathrm{SYN}}$ and $L_{\mathrm{MIS}}$; $L_{\mathrm{SYN}} + L_{\mathrm{MIS}} = L_{\mathrm{CDS}}$) from previous estimates of nonsynonymous to synonymous mutation rate ratio $r_L = \frac{\theta_{\mathrm{MIS}}}{\theta_{\mathrm{SYN}}}$. We used an $r_L$ of 2.31 for vertebrates ($L_{\mathrm{MIS}} = 2.31 * L_{\mathrm{SYN}}$; $L_{\mathrm{MIS}} = \frac{2.31}{2.31+1} * L_{\mathrm{CDS}}$), and an $r_L$ of 2.85 for invertebrates [3].

#### 1.3 *varDFE* package description

We developed a robust DFE comparison software, *varDFE*, as an extension for the $\partial a \partial i$ and Fit$\partial a \partial i$ packages. The *varDFE* package can be installed through `pip` in any python environment (version $\geq$ 3.10). Our workflow consists of four main modules (Fig. S2):

1. `Demog1D_sizechangeFIM`: Demographic inference on putatively neutral SFS data.
2. `DFE1D_refspectra`: Precomputation of reference SFS database under selection, given all possible selection coefficients $s$ and the inferred demographic scenarios in step one.
3. `DFE1D_inferenceFIM`: DFE inference on selected SFS data, given the assumed function form of the DFE and the pre-computed reference SFS database in step two.
4. `DFE1D_gridsearch`: DFE parameter space exploration through a grid search, given the assumed function form of the DFE and the pre-computed reference SFS database in step two.

Similar to `dadi-cli`(`https://dadi-cli.readthedocs.io/en/latest/`), each module in *varDFE* offers a streamlined and easy-to-use command line tool that simplifies the application of $\partial a \partial i$ in research settings. In addition, our package extends the functionality of $\partial a \partial i$ with the ability to explore continuous positive selection coefficients, infer parameters for a greater range of functional forms of the DFE, and automate a grid search for optimal parameter values when considering multiple datasets. It also provides automatic quality control features, including plotting, convergence testing, and quantifying uncertainty using Fisher's Information Matrix.

In the methods sections below, we illustrate the usage of *varDFE* workflow and specific settings used in this study.

#### 1.4 Demographic inference

We first inferred demographic parameters from the putatively neutral synonymous SFS (SYN-SFS) using the `Demog1D_sizechangeFIM` module in *varDFE* package. For each species, we fitted four models forward in time:

1. snm or one-epoch: standard neutral model with no population size change.
2. two-epoch: single population model with one size change event. The ratio of the current population size, i.e. the first size change, to ancestral population size (`nua`) and duration of size change (`Ta`, in units of $2 * N_a$ generations) are inferred.
3. three-epoch: single population model with two size change events. The ratio of the first size change population size, to ancestral population size (`nua`), the ratio of the current population size, i.e. the second size change, to ancestral population size (`nub`), and duration of two size change events (`Ta` and `Tb`) are inferred.
4. four-epoch: single population model with three size change events. The ratio of the first size change population size, to ancestral population size (`nua`), the ratio of the second size change population size, to ancestral population size (`nub`), the ratio of the current population size, i.e. the third size change, to ancestral population size (`nuc`), and duration of three size change events (`Ta`, `Tb` and `Tc`) are inferred.

To account for increased uncertainty in calling genotypes of rare variants, we also repeated the demographic inference by masking the singleton entries in the SYN-SFS. In total, eight demographic inference runs (four demographic models in SYN-SFS with or without singleton masks) for each species were conducted. All settings were the same across datasets except for the starting parameter positions, which were set from prior demographic inference (Datasets S1,S3). For datasets without prior demographic inference information (halictid bees, collared flycatchers, arctic wolves and gray wolves), we set the parameters to `nua, nub, nuc, Ta, Tb, Tc = 1` for initial runs and used the inferred demographic parameters as the starting positions in the final runs.

For each run, 100 replicates from a permuted starting parameter (`fold=1`) were run. Within each of the hundred replicates, we set extrapolation grid points as sample sizes plus 5, 15, and 25 [28], maximum iteration as 100, and performed parameter optimization using the `optimize_log` function based on the multinomial log-likelihood calculated from the expected SFS in each iteration. The best-fit parameters with the maximum multinomial log-likelihood in the 100 replicates were chosen for each run. We examined the convergence of the inference across replicates with different starting values by calculating the difference in log-likelihood in the 20 replicates with the highest log-likelihoods. The best-fit parameters' uncertainties were estimated through Fisher's Information Matrix (FIM, `Godambe.FIM_uncert` function in $\partial a \partial i$). To evaluate the best-fit parameters in the eight runs per species, we plotted each best-fit expected SFS with the observed SYN-SFS using ggplot2 (v.3.3.2 [30]) in R (v.3.6.2 [31]). We calculated the population-scaled synonymous mutation rate $\theta_{\text{SYN}}$, and estimated ancestral population size using $N_a = \frac{\theta_{\text{SYN}}}{4\mu L_{\text{SYN}}}$, where $\mu$ is the exon mutation rate for each species and $L_{\text{SYN}}$ is the previously estimated synonymous region sequence length. It is important to note that the exon mutation rate is hard to estimate in non-model organisms [32]. Existing estimates of $\mu$ can vary greatly. For example, in baleen whales, published mutation rates ranged from $5.7 \times 10^{-9}$/bp/gen [33] to $3.3 \times 10^{-8}$ [34]. To minimize the uncertainty in $\mu$, we selected the most widely used, phylogenetic-based and whole-genome-based mutation rate estimates in each species (Dataset S1).

To choose the best-fit demographic model and singleton-masking treatment out of the eight runs per (sub)species for downstream DFE inference, we performed a step-wise model selection procedure (Fig. S27). To maximize the data available and use the most parsimonious model whenever possible, at the start of model evaluation, the two-epoch demographic model based on full (unmasked) SYN-SFS run was chosen for all datasets. This "two-epoch, full SFS" run was chosen for a dataset, if it met all the criteria below:

1. The log-likelihood differences between the data (SYN-SFS) and model maximum likelihood $\Delta LL = LL_{\text{data}} - LL_{\text{demographic model}}$ for demographic inference is less than 200.
2. Based on the inferred demographic parameters from the evaluated run ("two-epoch, full SFS" in this case), the log-likelihood differences between the data (MIS-SFS) and model maximum likelihood $\Delta LL = LL_{\text{data}} - LL_{\text{DFE model|demographic model}}$ for the gamma DFE inference is less than 200.
3. The parameter estimates in the current demographic model are realistic. (unrealistic parameters defined as $\texttt{nu} < 10^{-5}$ or $\texttt{nu} > 20$ or $\texttt{T} < 10^{-5}$ or $\texttt{T} > 9$).

Otherwise, we proceeded to evaluate the more complex three- or four-epoch models with the same criteria. If the full/unmasked SFS resulted in poor model fit across all tested demographic models, we proceeded to mask the singletons, and continued the evaluation until one run met all criteria. This approach allows us to balance the risks of model overfitting and data loss, while ensuring a good model fit for each individual species.

Using this approach, the most parsimonious "two-epoch, full SFS" run was suitable for most species, except for the collared flycatchers, gray wolves, mosquitoes and halictid bees datasets. For the collared flycatchers, the "three-epoch, full SFS" run was chosen based on criteria 1, as the fit of demographic models greatly improved over the "two-epoch, full SFS" run ($\Delta LL_{\text{two-epoch}} = 1517$ and $\Delta LL_{\text{three-epoch}} = 1.35$). For the gray wolves, the "three-epoch, full SFS" run was chosen based on criteria

3, as the parameter estimates were extreme for the "two-epoch, full SFS" run (`nua` = 2.27, `nub` = 0.30, `Ta` = 1.59, `Tb` = 0.039 for the three-epoch model and `nua` = $8.5 \times 10^{-5}$, `Ta` = $4.5 \times 10^{-6}$ for the two-epoch model). For mosquitoes and halictid bees, all models based on the unmasked SFS failed to provide a good fit due to various reasons. The "three-epoch, masked SFS" run passed all criteria for mosquitoes and the "two-epoch, masked SFS" run was selected for halictid bees (details can be found at Dataset S4).

#### 1.5 DFE inference

Conditional on the inferred demographic scenarios, we estimated the DFE for new nonsynonymous mutations based on the MIS-SFS for each species using methods from Fit$\partial a \partial i$ [35]. Briefly, Fit$\partial a \partial i$ takes advantage of the pattern that more deleterious mutations are less likely to segregate in the sample of individuals and those that are segregating are more likely to be at lower frequency compared to neutral mutations. Assuming the MIS-SFS is under selection while the SYN-SFS is putatively neutral, the best DFE parameters should fit the differences between MIS-SFS and SYN-SFS in each minor allele frequency bin. Two modules in *varDFE*, `DFE1D_refspectra` and `DFE1D_inferenceFIM`, provide a flexible workflow to estimate both the full and deleterious-only DFE with any given demographic and DFE models. To ensure consistency across species, during DFE inference, we only allowed species-specific settings including demographic parameters, mutation rates ($\mu$), length of nonsynonymous region ($L_{\mathrm{MIS}}$), population-scaled mutation rates ($\theta$) to vary across species.

We first computed and stored the expected SFS for a range of population-scaled selection coefficients $\gamma = 2N_a s$, given the best fit demographic scenarios using the `DFE1D_refspectra` module in *varDFE*. Here we calculated the reference spectra given $\gamma$ but not $s$, because the SFS reflects the effect of selection at population level, which depends on $\gamma = 2N_a s$ but not $s$ alone. We implemented a slightly modified `Cache1D_mod` module in $\partial a \partial i$, named `Cache1D_mod2`, with a method to integrate over continuous positive gamma space. The range of positive gammas was set from $+10^{-5}$ to +100, with 701 points evenly distributed on the log10 scale, and negative gammas from -10000 to $-10^{-5}$, with 901 points evenly distributed on the log10 scale. The extrapolation points (`pts_l`) were set as `[1000,1200,1400]`. The reference spectra, from which the expected SFS given any $\gamma$ can be computed, were cached to save time for recalculations and improve consistency.

##### 1.5.1 DFE inference assuming the gamma DFE

To infer the deleterious-only DFE ($s \leq 0$), we assumed that the DFE follows a gamma distribution and performed the inference using the `DFE1D_inferenceFIM` module. We used the gamma distribution because it had strong support from theoretical models, such as the FGM [36], and was shown to fit the data well in previous studies [3, 37]. We parameterized the DFE for each species as $\gamma \sim Gamma(\alpha, \beta)$, where $\alpha$ ($\alpha > 0$; shape) and $\beta$ ($\beta > 0$; scale) are parameters to be inferred.

To avoid finding local maxima, for each species, 100 replicates were run from a permuted starting parameter (fold=1) of $\alpha = 0.2$ and $\beta = 4000$, informed from prior DFE estimations [3]. We set parameter upper bounds at $\max(\alpha) = 2.0$ and $\max(\beta) = 10^6$, lower bounds at $\min(\alpha) = 0.001$ and $\min(\beta) = 0.01$, maximum iteration as 100. The population-scaled nonsynonymous mutation rate $\theta_{\mathrm{MIS}}$ was pre-calculated from the $\theta_{\mathrm{SYN}}$ in the best-fit demographic model, using a scaling factor $r_L$ of 2.31 for vertebrates and $r_L$ of 2.85 for invertebrates [3]. We performed parameter optimization using the `optimize_log` function based on Poisson log-likelihood calculated from the expected SFS, generated using the `integrate` method from the precomputed reference spectra, in each iteration.

The best-fit parameters with the maximum Poisson log-likelihood in the 100 replicates were chosen for each run. We evaluated replicate convergence across starting values by calculating the difference in log-likelihood in the 20 replicates with the highest log-likelihoods. The AIC for each run was calculated using the equation: $AIC = 2k - 2ln(\hat{L})$, where $ln(\hat{L})$ is the highest log-likelihood associated with the best-fit parameters in DFE inference, and $k$ is the total number of estimated parameters for the DFE and underlying demographic model. For example, $k = 5$ in a DFE inference run assuming a gamma-distributed DFE and a two-epoch demographic model, because two parameters (shape and scale) are inferred during DFE inference, and three parameters (`nua`, `Ta`, and $\theta_{\mathrm{SYN}}$) are inferred during demographic inference. Recall that we optimized the gamma distribution parameters for the population-scaled selection coefficient $\gamma$. To obtain the distribution of $s$, we unscaled the gamma distribution by $1/(2N_a)$, therefore, $s \sim Gamma(\alpha, \beta')$, where $\beta' = \beta/2N_a$ and $N_a$ had been inferred from demography estimation.

The expected selection coefficient $E[|s|]$ for each species was calculated from the best-fit parameters given the gamma distribution's property that $E[|s|] = \alpha\beta'$. To compute the proportion of mutations with different values of $s$ in the maximum-likelihood gamma distribution, we found the cumulative probability for $|s|$ ranged in [0, $10^{-5}$), [$10^{-5}$, 0.0001), [0.0001, 0.001), [0.001, 0.01), and [0.01, 0.5] using the `pgamma` function in R. The expected population-scaled mutation effects ($E[|2N_a s|]$) were calculated as $E[|2N_a s|] = \alpha\beta$. The cumulative probability for $|2N_a s|$ ranged in [0,1), [1, 10), [10, 100), [100, 1000) and [1000, Inf] was calculated using the `pgamma` function in R as well. For uncertainty estimates, we followed the methods described in [35, 37]. Briefly, for each species, we Poisson resampled the MIS-SFS 200 times, and refit the DFE on the resampled data. We calculated the expected selection coefficients, and the proportion of mutations with different values of $s$ using these 200 estimates. The 95% confidence intervals were determined using the 2.5% and 97.5% quantiles of the resampled estimates.

##### 1.5.2 Inferring the proportion of neutral mutations

To estimate the proportion of effectively neutral mutations, we added a point mass at neutrality to the original gamma DFE distribution. The DFE function can be written as $\gamma \sim Neugamma(\alpha, \beta, p_{\mathrm{neu}})$. $\alpha$ ($\alpha > 0$; shape) and $\beta$ ($\beta > 0$; scale) are parameters from a gamma distribution, and $p_{\mathrm{neu}}$ ($p_{\mathrm{neu}} \geq 0$) is the proportion of neutral mutations ($0 \leq |\gamma| < 10^{-5}$).

$$f(\gamma) = \begin{cases} \frac{p_{\mathrm{neu}}}{10^{-5}} + (1 - p_{\mathrm{neu}}) * Gamma(\gamma|\alpha, \beta), & 0 \leq |\gamma| < 10^{-5} \\ (1 - p_{\mathrm{neu}}) * Gamma(\gamma|\alpha, \beta), & |\gamma| \geq 10^{-5} \end{cases} \tag{1}$$

During inference, we set the parameter start positions, upper bounds and lower bounds the same as the gamma distribution for the $\alpha$ and $\beta$ parameters. The starting position for $pneu$ is 0.3, the upper bound is $\max(p_{\mathrm{neu}}) = 1$, and the lower bound is $\min(p_{\mathrm{neu}}) = 10^{-5}$.

To obtain the distribution of $s$, we unscaled the neugamma distribution by $1/(2N_a)$, therefore, $s \sim Neugamma(\alpha, \beta', p_{\mathrm{neu}})$, where $\beta' = \beta/(2N_a)$ and $N_a$ had been inferred from the demographic model. The expected selection coefficient $E[|s|]$ for each species was calculated from the best-fit parameters using $E[|s|] = \frac{p_{\mathrm{neu}}}{2*2N_a} * 10^{-5} + (1 - p_{\mathrm{neu}})\alpha\beta'$.

##### 1.5.3 Inferring the proportion of lethal mutations

To estimate the proportion of effectively lethal mutations, we added a point mass at lethality to the original gamma DFE distribution. The DFE function can be written as $\gamma \sim Gammalet(\alpha, \beta, p_{\text{let}})$. $\alpha$ ($\alpha > 0$; shape) and $\beta$ ($\beta > 0$; scale) are parameters from a gamma distribution, and $p_{\text{let}}$ ($0 \leq p_{\text{let}} < 0.5$) is the proportion of lethal mutations ($|s| = 0.5$).

$$f(\gamma) = \begin{cases} (1 - p_{\text{let}}) * Gamma(\gamma|\alpha, \beta), & 0 \leq |\gamma| < N_a \\ p_{\text{let}}, & |\gamma| = N_a \end{cases} \tag{2}$$

During inference, we set the parameter start positions, upper bounds and lower bounds the same as the gamma distribution for the $\alpha$ and $\beta$ parameter. The starting position for $p_{\text{let}}$ is 0.001, the upper bound is $\max(p_{\text{let}}) = 0.5$, the lower bound is $\min(p_{\text{let}}) = 10^{-5}$.

To obtain the distribution of $s$, we unscaled the gammalet distribution by $1/(2N_a)$, therefore, $s \sim Gammalet(\alpha, \beta', p_{\text{let}})$, where $\beta' = \beta/(2N_a)$ and $N_a$ had been inferred from demography estimation. The expected selection coefficient $E[|s|]$ for each species was not calculated for this distribution, given the presence of lethal mutations.

#### 1.6 Assessing the robustness of DFE inference

##### 1.6.1 Individual effects of competing models

To examine the robustness of our DFE inference, we implemented three functional forms of the DFE in addition to the gamma distribution: neugamma, gammalet, and lognormal. For this analysis, we kept the same demographic models and singleton masking treatments as used in the original analysis shown in Table 1.

The functional forms for the neugamma and gammalet distributions have been described above. For the lognormal distribution, we parameterized the DFE for each species as $\gamma \sim Lognormal(\mu_s, \sigma^2)$. During inference, we set the starting positions before perturbation as $\mu_s = 1, \sigma = 0.1$, upper bounds as $\max(\mu_s) =$ 100, $\max(\sigma) = 100$, lower bounds as $\min(\mu_s) =$ -100, $\min(\sigma) = 10^{-5}$. Since $\mu_s$ could take a negative value, we used the `optimize` instead of `optimize_log` function for parameter optimization. To obtain the distribution of $s$, we unscaled the lognormal distribution, therefore, $s \sim Lognormal(\mu_s^{'}, \sigma^2)$, where $\mu_s^{'} = \mu_s - ln(2N_a)$ and $N_a$ had been inferred from demography estimation. Median mutation effects $E[|s|]$ for each species was calculated using $E[|s|] = exp(\mu_s^{'})$ [3].

Considering that different demographic models were used across species during the DFE inference, we investigated whether the specific choices of these models influenced our results. The reference spectra for all species assuming two-epoch, three-epoch or four-epoch demographic models were calculated. Assuming a gamma-distributed DFE, we repeated the DFE inference process under different demographic assumptions with the best-fit singleton-masking treatments.

Assuming a gamma-distributed DFE and the best-fit demographic models, we repeated the DFE inference process for each species using both unmasked and singleton-masked SYN-SFS and MIS-SFS.

##### 1.6.2 Combinative effects of competing models

To study the combinative effects of competing models, we utilized a model averaging approach as described in [38]. For each species, we performed DFE inference assuming three DFE functional forms (gamma, neugamma, and lognormal) under both two-epoch and three-epoch demographic models, with and without singleton masking. As masking singleton changes the data log-likelihood, for each species, we generated separate model-averaged DFE estimates based on either the singleton-masked SFS or the full SFS. For each species and singleton-masking treatment, six combinations of DFE and demographic models were evaluated: 1. gamma-distributed DFE under two-epoch demography, 2. gamma-distributed DFE under three-epoch demography, 3. neugamma-distributed DFE under two-epoch demography, 4. neugamma-distributed DFE under three-epoch demography, 5. lognormal-distributed DFE under two-epoch demography, 6. lognormal-distributed DFE under three-epoch demography. We computed the model weights of six combinations according to equations described in [38], and generated weighted-averages for the expected mutation effects ($E[|s|]$) in each species and singleton-masking treatment. To facilitate comparisons with the main text results, we also generated model-averaged $E[|s|]$ estimates using the best singleton-masking treatment, where only the mosquitos and halictid bees data are masked whereas the full SFS are used in the other nine species.

#### 1.7 Testing for phylogenetic signal

To further evaluate whether the DFE co-varies with phylogeny, we calculated Pagel's $\lambda$ using the `phylosig` function from the phytools package (v.1.2.0 [39]) in R. The log10 transformed expected selection coefficient ($\log_{10}(E[|s|])$) and the shape parameter ($\alpha$), derived from the gamma-distributed DFE, were considered candidate traits to test for phylogenetic dependence. In addition, we calculated the Pagel's $\lambda$ for life history traits, including the generation time, mutation rate, body mass, age at maturity, maximum longevity and the long-term population size (Dataset S13). The underlying phylogenetic structure and divergence times for eleven (sub)species were compiled from the TimeTree database (http://timetree.org/) based on their scientific names.

To formally test whether the DFE differs across different species, we compared each species' DFE estimates to a null model where the DFE was constrained to be the same across species by likelihood-ratio tests (LRT). To do this, we employed a grid-search approach using the `DFE1D_gridsearch` module in the *varDFE* package.

To test for differences in $s$ across species, we evenly spaced the same 2000 grid points in biologically meaningful ranges of shape ($\alpha$ = 0.1 to 0.5) and scale ($\beta'$ = 0.0001 to 0.5) parameters. We obtained the expected SFS for each $\alpha - \beta'$ pair for each species using the `integrate` method from reference spectra, and calculated the Poisson log-likelihood relative to empirical MIS-SFS. From four million $\alpha - \beta'$ pairs, we explored the full Poisson log-likelihood surface for each species simultaneously. To obtain the null model where the DFE is constrained to be the same across species, we assumed that each species' SFS is independent of each other given their distant phylogenetic divergence. Therefore, the log-likelihood for the null model can be calculated by summing the log-likelihood at each $\alpha - \beta'$ pair for each species and the MLE can be found from within the four million $\alpha - \beta'$ parameter grids. To formally test whether the shape ($\alpha$) and scale ($\beta'$) are different in any $x$ number of species, we used a likelihood ratio test [3]. The LRT was constructed as $\Lambda = -2 * ln(L_0/L_1) = -2 * (LL_0 - LL_1)$, where $LL_0$ is the log-likelihood for the null model, with two DFE parameters inferred ($\hat{\alpha_1} = \hat{\alpha_2} = ... = \hat{\alpha_x}, \hat{\beta'_1} = \hat{\beta'_2} = ... = \hat{\beta'_x}$; params = 2) given the inferred demographic parameters for each species ($\Theta_{D,1}, \Theta_{D,2}, ..., \Theta_{D,x}$), and $LL_1$ is the log-likelihood for the alternate model, where each species is allowed to have its own DFE parameters ($\hat{\alpha_1}, \hat{\alpha_2}, \hat{\alpha_x}, ..., \hat{\beta'_1}, \hat{\beta'_2}, \hat{\beta'_x}$; params = $2x$). Across the eleven (sub)species tested, asymptotically, $\Lambda$ should follow a $\chi^2$ distribution with $df = 2x - 2 = 20$.

$$\Lambda = -2 * ln[\frac{L(\alpha_1 = \alpha_2 \widehat{= ...} = \alpha_x, \beta'_1 = \beta'_2 \widehat{= ...} = \beta'_x | \Theta_{D,1}, \Theta_{D,2}, ..., \Theta_{D,x})}{L(\hat{\alpha_1}, \hat{\alpha_2}, ..., \hat{\alpha_x}, \hat{\beta'_1}, \hat{\beta'_2}, ..., \hat{\beta'_x} | \Theta_{D,1}, \Theta_{D,2}, ..., \Theta_{D,x})}] \quad (3)$$

To demonstrate the phylogenetic correlations in the DFE evolution, we calculated the pairwise likelihood ratio test statistics in all possible pairs of species ($x = 2$). For example, the LRT statistics comparing humans (population 1) and *Drosophila* (population 2) DFE can be written as follows:

$$\Lambda = -2 * ln[\frac{L(\alpha_1 \widehat{=} \alpha_2, \beta'_1 \widehat{=} \beta'_2 | \Theta_{D,1}, \Theta_{D,2})}{L(\hat{\alpha_1}, \hat{\alpha_2}, \hat{\beta'_1}, \hat{\beta'_2} | \Theta_{D,1}, \Theta_{D,2})}] \quad (4)$$

Asymptotically, $\Lambda$ should follow a $\chi^2$ distribution with $df = 2x - 2 = 2$. We hierarchically clustered the log10 scaled LRT statistics [$\log_{10}(\Lambda)$] using the `hclust` function in R.

To test the variations of population-scaled DFE, $\gamma = 2N_a s$, for each species, we conducted the grid search by evenly spacing the same 2000 grid points in biologically meaningful ranges of shape ($\alpha = 0.1$ to 0.5) and population-scaled scale ($\beta = 100$ to $2.5 \times 10^4$) parameters for population-scaled DFE, and repeated the overall and pairwise LRT analyses as outlined above.

For the overall LRT analyses in population-scaled DFE, The LRT statistics for $x$ populations can be written as follows:

$$\Lambda = -2 * ln[\frac{L(\alpha_1 = \alpha_2 \widehat{= ...} = \alpha_x, \beta_1 = \beta_2 \widehat{= ...} = \beta_x | \Theta_{D,1}, \Theta_{D,2}, ..., \Theta_{D,x})}{L(\hat{\alpha_1}, \hat{\alpha_2}, ..., \hat{\alpha_x}, \hat{\beta_1}, \hat{\beta_2}, ..., \hat{\beta_x} | \Theta_{D,1}, \Theta_{D,2}, ..., \Theta_{D,x})}] \quad (5)$$

For the pairwise LRT analyses in population-scaled DFE, The LRT statistics for two populations (population 1 and 2) can be written as follows:

$$\Lambda = -2 * ln[\frac{L(\alpha_1 \widehat{=} \alpha_2, \beta_1 \widehat{=} \beta_2 | \Theta_{D,1}, \Theta_{D,2})}{L(\hat{\alpha_1}, \hat{\alpha_2}, \hat{\beta_1}, \hat{\beta_2} | \Theta_{D,1}, \Theta_{D,2})}] \quad (6)$$

#### 1.8 Correlating life history traits with the DFE variation across species (PGLS$_\lambda$)

To test if the candidate life history traits ($X$) are correlated with the observed $E[|s|]$ ($Y$), and correct for phylogenetic dependency when appropriate, we utilized the PGLS$_\lambda$ method (phylogenetic generalized least squares; [40]), implemented by the ape (v.5.4.1 [41]), nlme (v.3.1.149 [42]) and phytools packages in R. The PGLS$_\lambda$ method optimizes the phylogenetic signal of the residuals of $E[|s|]$ ($\lambda$) simultaneously with the regression parameters (function `phyl.resid` in phytools). Importantly, the $\lambda$ estimated in PGLS$_\lambda$ does not equate to the Pagel's $\lambda$ estimated individually for $X$ or $Y$ (described in the previous section), but represents whether the correlation of $X$ and $Y$ can be accounted for by phylogenetic dependence in residual error. Compared to the default PGLS model with a Brownian motion correlation structure ($\lambda = 1$), it allows for a more flexible variance-covariance matrix, therefore, improving model performances by applying an optimized degree of phylogenetic correction.

In our analysis, the candidate explanatory variables ($X$) included are:

1. Long-term population size ($N_a$);
2. Body mass in grams;
3. Generation time in years per generation;
4. Age at maturity in days;

5. Maximum longevity in years;
6. Mutation rates in mutations per bp per generation.

The regression was specified as $\log_{10}(E[|s|]) \sim \log_{10}(X)$ to normalize the data. Given that the shape ($\alpha$) parameter is often estimated more confidently, we repeated PGLS$_\lambda$ analysis for $\alpha$, with the regression specified as $\alpha \sim \log_{10}(X)$. Since parameters for the arctic wolf dataset reached upper boundaries during inference (Fig. 1B, Dataset S6), we tested the impact on regression outputs when this data point was excluded (Dataset S14).

#### 1.9 Fitting the FGM-derived DFE

To investigate the genetic mechanism underlying DFE variations and incorporate beneficial mutations in our inference, we implemented a functional form of the DFE which is directly derived from the Fisher's geometric model [43]. Briefly, the FGM proposes that organisms can be seen as a collection of phenotypes under stabilizing selection around a local maximum. A genotype generates a single phenotype, which is characterized by a point in an $n$-dimension phenotypic space. The phenotypic space's dimensionality, in other words, the total number of phenotypes under selection, is defined as complexity ($n$). Fitness ($w$) can be described as a Gaussian function of the distance to the optimum ($z$), $w(z) = exp(-z^2)$. Random mutations do not impact all phenotypes equally, but will likely affect a subset of $m$ phenotypes with fitness effect size $r$. Here, $m$ refers to mutation pleiotropy [43]. Effect size $r$ follows a zero mean Gaussian distribution with scale $\sigma$. When the population is under mutation-selection-drift balance, the increase in fitness from beneficial mutations should counteract the drift load, the decrease in fitness caused by deleterious mutations. The population will have an equilibrium phenotypic distance ($z_{\text{eq}}$) to the fitness optimum. Therefore, the DFE for the well-adapted population (equation 15 in [43]) can be described using the mutation pleiotropy ($m$), scale of mutation effects ($\sigma$) and long-term population size ($N_a$), $\gamma \sim Lourenco(m, \sigma, N_a)$. Here $\Gamma(.)$ is the gamma function and $K(.)$ is the modified Bessel function of the second kind [43]. We note that there is a missing exponential term in equation 15 in [43], which is corrected in the equation below.

$$f(s) = \frac{2^{\frac{1-m}{2}}\sqrt{N_a}(|s|)^{\frac{m-1}{2}}(1+\frac{1}{N_a\sigma^2})^{\frac{1-m}{4}}exp(-N_a s)}{\sqrt{\pi}\sigma^m\Gamma(\frac{m}{2})} \times K_{\frac{m-1}{2}}(N_a|s|\sqrt{1+\frac{1}{N_a\sigma^2}}) \quad (7)$$

While complex in its expression, this FGM-derived distribution is functionally equivalent to the previously described gamma and lognormal distributions for generating mutational effects. Therefore, we estimated its parameters ($m$, $\sigma$, and $N_a$) following methods described in the DFE inference section, through maximum-likelihood optimization by fitting the observed differences between MIS-SFS and SYN-SFS across minor allele frequency bins. We inferred the maximum likelihood estimates of the $m$, $\sigma$, and $N_a$ parameters using the `DFE1D_inferenceFIM` module as described in the DFE inference section. Since this DFE considers beneficial mutations, we calculated the expected SFS using the `integrate_continuous_pos` method implemented in our *varDFE* package. To compute the proportion of mutations with different values of $s$ in the inferred FGM-derived DFE, we found the cumulative probability for $s$ ranged in [-0.5, -0.01), [-0.01, -0.001), [-0.001, -0.0001), [-0.0001, -$10^{-5}$), [-$10^{-5}$, 0), [0, $10^{-5}$), [$10^{-5}$, 0.0001), and [0.0001, 0.5] using the `scipy.integrate.quad` function (v.1.8.0) in python. The proportion of beneficial mutations was obtained by calculating the cumulative probability for $s > 0$.

To compare the correlations of $N_a$ inferred in this FGM-derived DFE with the $N_a$ obtained in demographic inference, we performed linear regression $\log_{10}(N_a^{\text{DFE}}) \sim \log_{10}(N_a^{\text{Demog}})$ using the `lm` function in R. A linear regression between the proportion of beneficial mutations and $N_a$ was also performed: $P(s > 0) \sim \log_{10}(N_a^{\mathrm{Demog}})$ using the same method. To test for phylogenetic signal in the inferred DFE parameters, we calculated Pagel's $\lambda$ for $m$ and $\sigma$ using methods described above (Dataset S13).

### 2 Text S2: Estimating proportion of lethal mutations using the gammalet DFE

We attempted to directly estimate the proportion of effectively lethal mutations ($s = -0.5$), by assuming that the DFE follows a mixture of a gamma distribution with additional lethal point mass (gammalet DFE [44], Dataset S8).

Specifically, the gammalet DFE is constructed by adding a point mass at lethality ($0 \leq p_{\text{let}} < 0.5$) to the original gamma DFE distribution. The DFE function can be written as $\gamma \sim Gammalet(\alpha, \beta, p_{\text{let}})$. Here, $\gamma = 2N_a s$, is the population-scaled selection coefficients.

$$f(\gamma) = \begin{cases} (1 - p_{\text{let}}) * Gamma(\gamma|\alpha, \beta), & 0 \leq |\gamma| < N_a \\ p_{\text{let}}, & |\gamma| = N_a \end{cases} \tag{8}$$

The distribution of $s$ is $s \sim Gammalet(\alpha, \beta', p_{\text{let}})$, where $\beta' = \beta/(2N_a)$. $E[|s|]$ was not calculated for this distribution, given the presence of lethal mutations.

Only three datasets (mosquitoes, arctic wolves, vaquitas) obtained a lowered AIC score for the gammalet DFE compared with the gamma DFE (Figs. S6, S8). In these datasets, the proportion of lethal mutations reached unrealistic high values (31% in vaquitas to 50% in mosquitoes, Fig. S7).

Given that lethal mutations are more likely to be recessive, it challenges Fit$\partial a \partial i$'s underlying assumption of additive fitness effects for all mutations. Therefore, we suspected that our current approach lacks the power to confidently infer the proportion of lethal mutations [35, 44].

### 3 Text S3: DFE inference is mostly robust to the assumed functional form of the DFE, demographic models, and inclusion of singletons

To evaluate the robustness of our analysis, we tested if different probability distributions for the DFE, demographic models, or decisions to mask singletons in the SFS affected DFE inference (Fig. S9).

In addition to the three DFE distributions (gamma, neugamma, gammalet) based on gamma DFE, we also examined the lognormal distribution (Fig. S10, Dataset S9). Given that the gammalet DFE performed poorly regarding model fit and parameter estimations compared to the gamma DFE, we focus our comparisons on three functional forms: gamma, neugamma, and lognormal (Fig. S9A). The neugamma DFE had the lowest AIC, i.e. most supported by the data, in seven datasets (*Drosophila*, halictid bees, pied flycatchers, collared flycatchers, fin whales, mice, and humans), medium AIC in arctic wolves and highest AIC in mosquitoes, gray wolves and vaquitas datasets. The lognormal DFE had the highest AIC in the seven datasets where the neugamma DFE performed the best and had the lowest AIC in the other four datasets (mosquitoes, gray wolves, arctic wolves, and vaquitas; Fig. S12). Given that the gamma DFE is more parsimonious than the neugamma DFE, consistently had the second-lowest AIC of all functional forms in all datasets except for the arctic wolves (Fig. S12), and provided a good visual fit to the observed SFS (Fig. S4), we confirmed that the gamma distribution is a good candidate function form for DFE comparisons.

Next, we evaluated how estimates of the expected selection coefficient ($E[|s|]$) and proportion of each mutation category varied across different functional forms of the DFE (Figs. S9A, S11). When assuming the neugamma DFE, $E[|s|]$ remains more deleterious in mammals compared with insects and birds (Fig. S9A). When assuming the lognormal DFE, the phylogenetic signal is less evident compared with the gamma DFE, due to the overall reduced $E[|s|]$ estimates in mammals (average $E[|s|]_{\text{gamma}} = 0.14$, $E[|s|]_{\text{lognormal}} = 0.044$, n = 6; Fig. S9A). However, the log-likelihood for lognormal DFE is worse than that of the gamma DFE in fin whales, mice, and humans, suggesting that the $E[|s|]$ obtained here is less supported by the data.

We also tested the potential impacts of demographic model misspecification, given that different demographic models were selected in each dataset. Assuming the DFE follows a gamma distribution, we repeated the DFE inference for each dataset using the demographic parameters inferred from two-, three-, and four-epoch models (Dataset S10). Notably, in some datasets, unreasonable demographic model assumptions could lead to failed DFE inference, with the difference between the log-likelihoods of data and model ($\Delta LL$) exceeding 200 units, such as two-epoch models in the mosquitoes and collared flycatchers datasets (translucent points in Fig. S9B). Overall, the average mutation effects and proportion of each mutation category remain unchanged qualitatively (Figs. S9B, S13). However, when assuming three-epoch or four-epoch model in the mice dataset, $E[|s|]$ is reduced from 0.010 (two-epoch) to 0.00089 (three-epoch) or 0.0014 (four-epoch). However, the two-epoch based $E[|s|]$ estimate is more appropriate for the mice dataset, because there is minimal improvement in log-likelihood and visual fit for the three- and four-epoch models (Fig. S3), and the estimated parameters in these models reflected extreme demographic events (Dataset S4).

Finally, we tested whether masking singletons in the SFS during demography and DFE inference altered the $E[|s|]$ estimation (Dataset S11). Again, the average mutation effects and proportion of each mutation category are largely unchanged qualitatively (Figs. S9C, S14), except that $E[|s|]$ is reduced from 0.013 (not masking singletons) to 0.0023 (masking) in the vaquitas dataset.

In summary, our examination of different functional forms of the DFE, demographic models, and singleton masking strategies, suggest that the DFE inference is mostly robust to choices made during the analysis.

### 4 Text S4: PGLS$_\lambda$ analyses excluding outliers and for the shape parameter

Given that the DFE parameters for arctic wolves reached upper boundaries with uncertain biological significance, we repeated the PGLS$_\lambda$ analysis excluding the arctic wolves data point. Correlations of $N_a$ and body mass with $E[|s|]$ remained significant (P = 0.005 and 0.001 respectively). Notably, mutation rates became significantly correlated with $E[|s|]$ after the exclusion (P = 0.008; Dataset S14).

Since the shape ($\alpha$) parameter is often estimated more confidently than the scale parameter in DFE inference [35, 45], we performed the PGLS$_\lambda$ analysis between $\alpha$ and life history traits (Fig. S21). Long-term population size was positively correlated with $\alpha$ (P = 0.002), whereas body mass was negatively correlated with $\alpha$ (P $<$ 0.001). Although not significantly correlated with $E[|s|]$, age at maturity is negatively correlated with $\alpha$ (P = 0.004). For full results, see Dataset S14.

568 # 5 Supplementary Figures

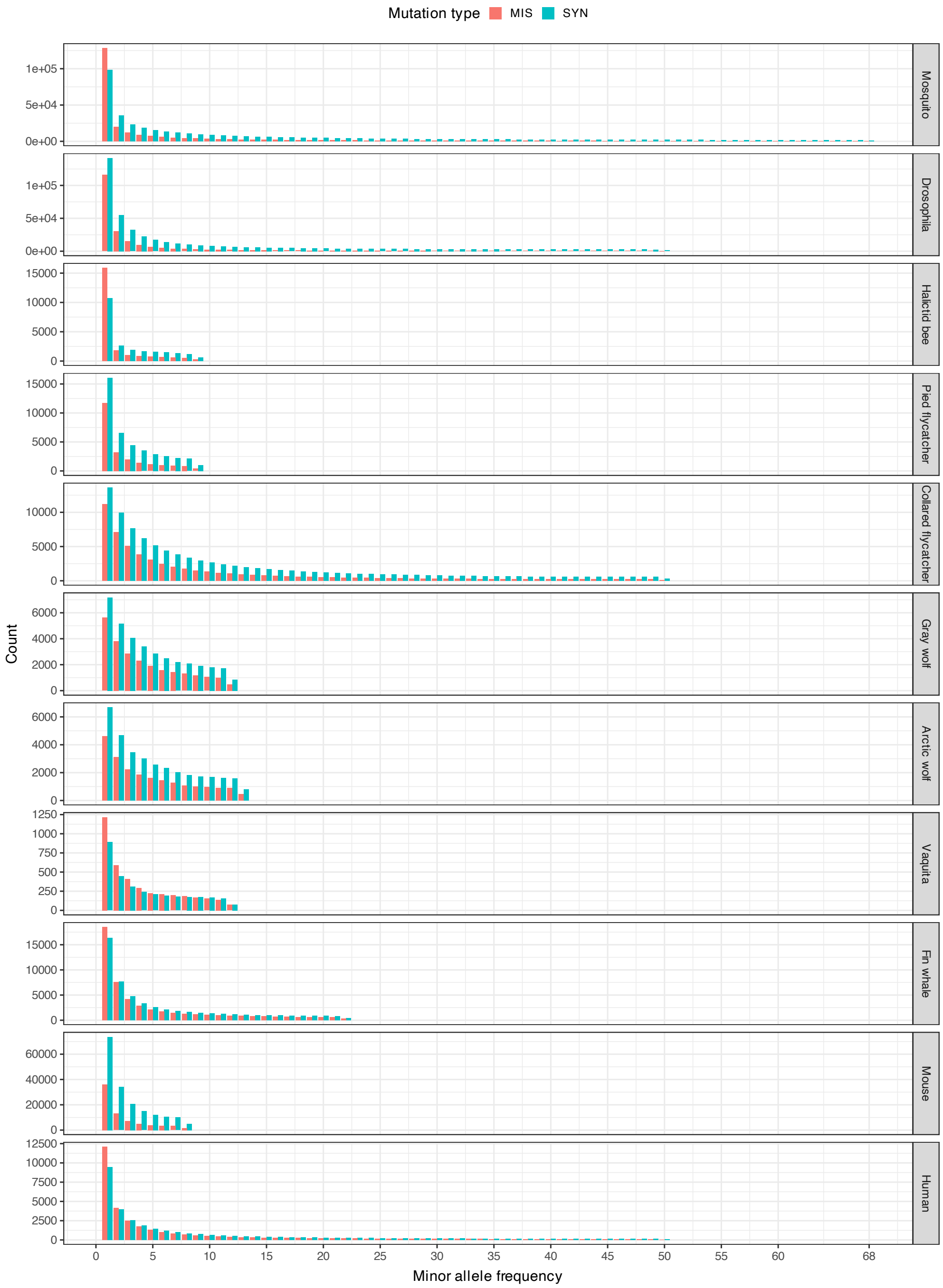


Figure S1: The projected folded SFS for the eleven (sub)species datasets. The red bars represent nonsynonymous/missense SFS while the cyan bars show the synonymous SFS. Dataset names are marked at the right.

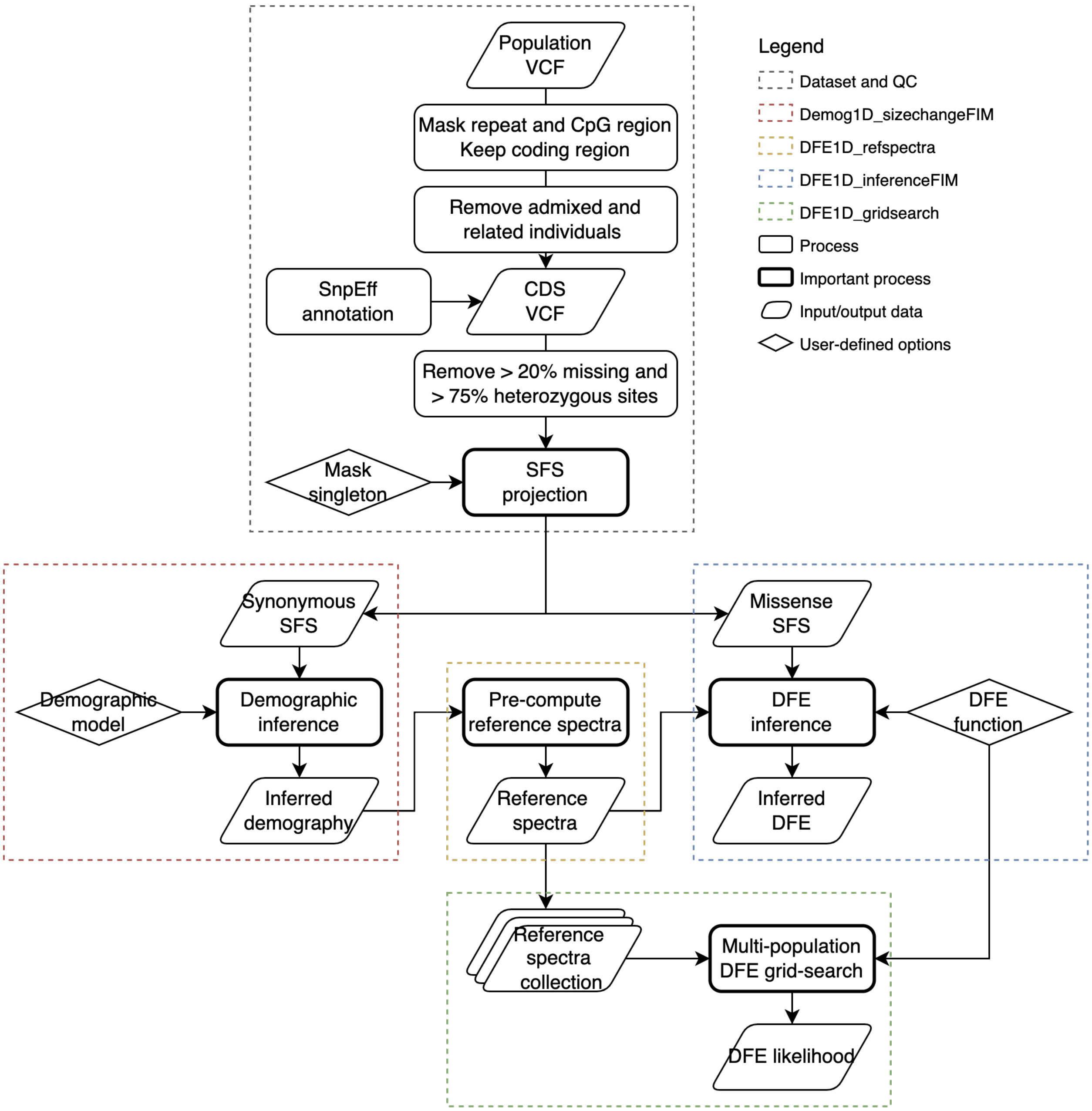


Figure S2: The *varDFE* API workflow. Each colored dashed rectangle represents a module in *varDFE*. The gray dashed rectangle shows the quality control and preprocessing steps for the eleven (sub)species datasets that are not part of *varDFE* API.

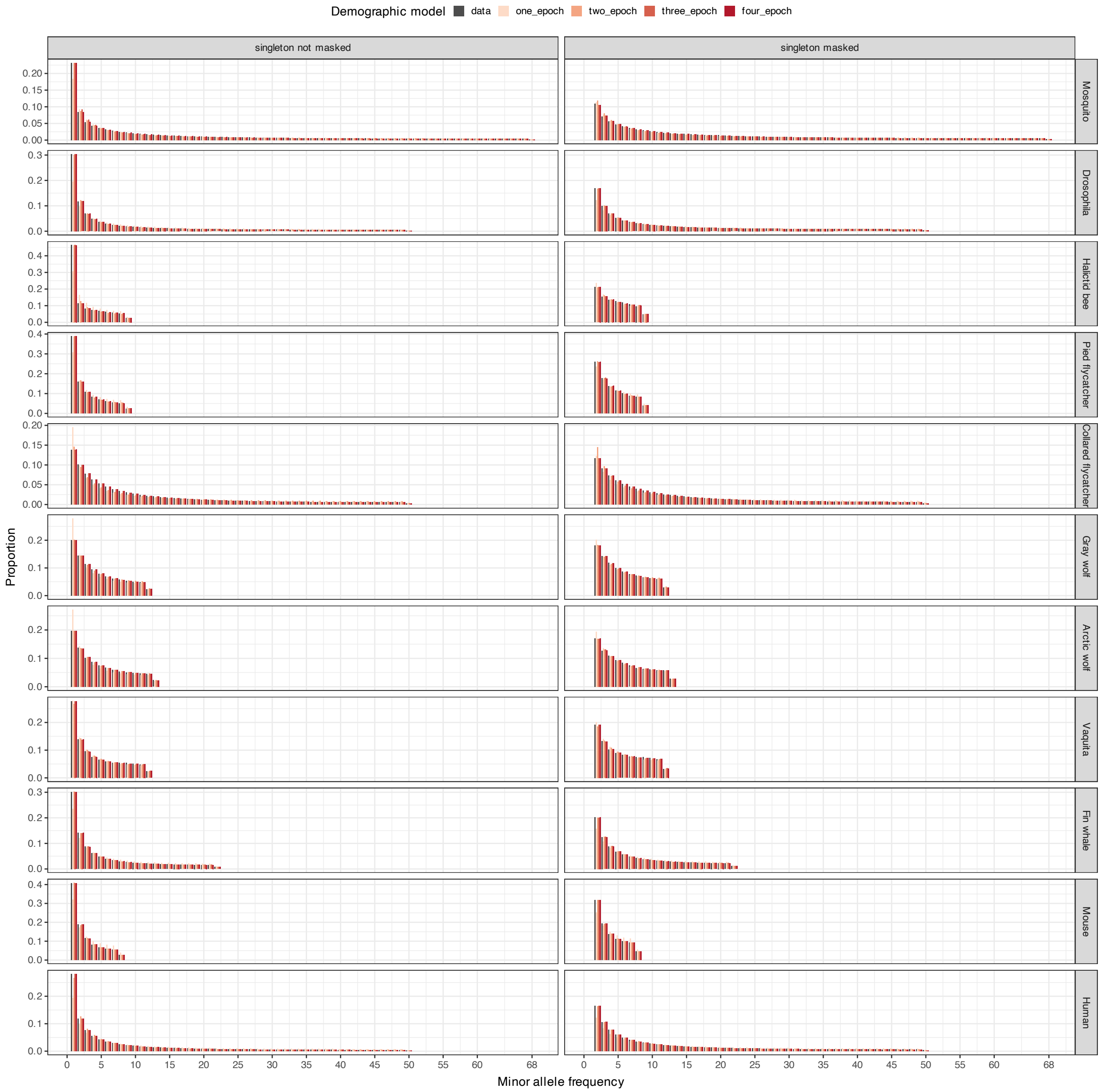


Figure S3: Comparison of the observed synonymous SFS (SYN-SFS) to the predicted SFS under different demographic models. For each species (each row), the demographic inference results using the full SFS (left column) or singleton-masked SFS (right column) are shown. Within each panel, the proportional SYN-SFS data is plotted as the left-most bar, and the maximum-likelihood estimate's expected SFS for the standard neutral model (one-epoch), two-epoch, three-epoch and four-epoch models are shown to the right of the data. Note the overall fit of the two-epoch model using the full SFS and the improved fit of respective chosen models for the mosquitoes, halictid bees, gray wolves, and collared flycatchers datasets.

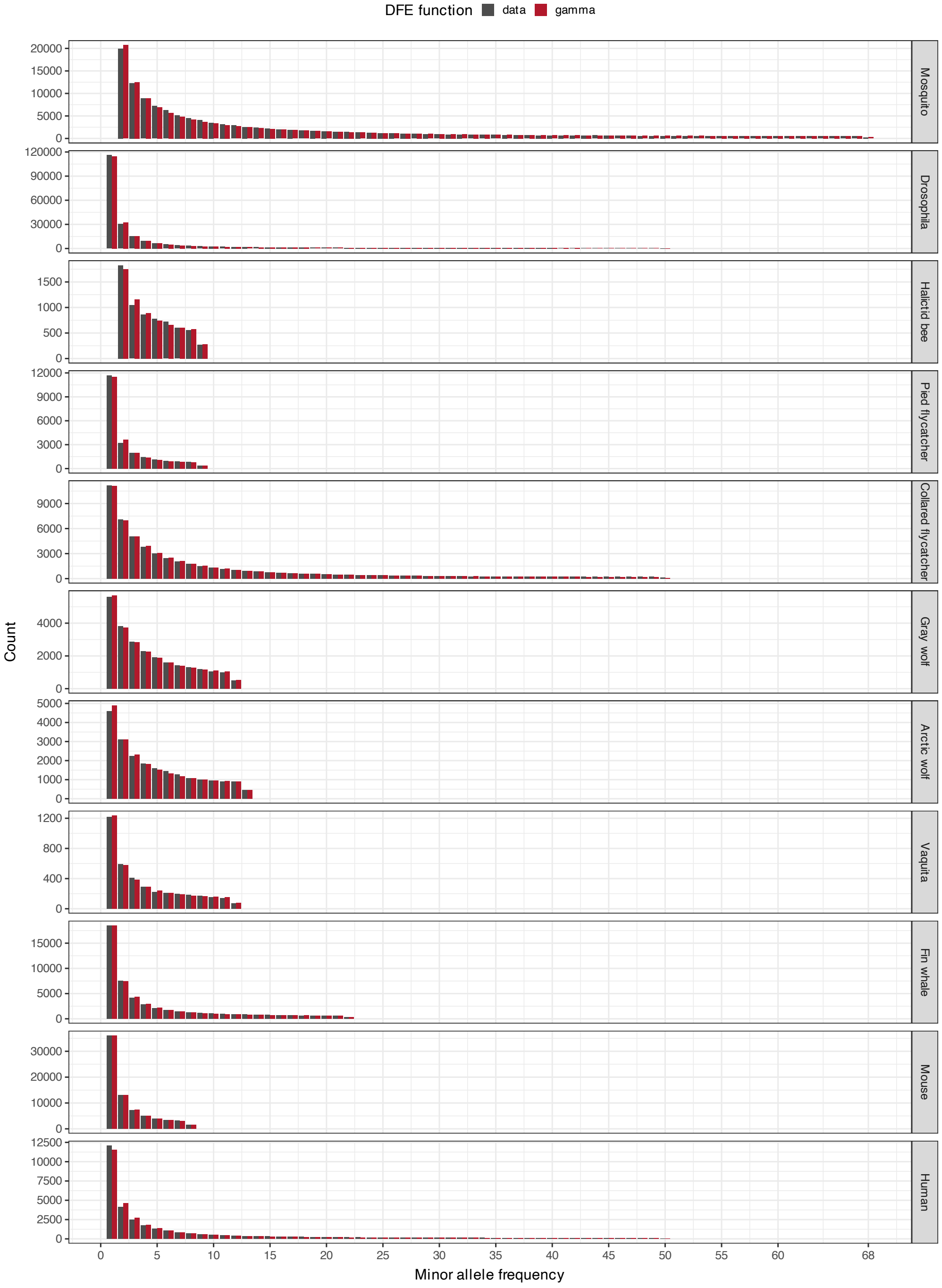


Figure S4: Comparison of the observed nonsynonymous SFS (MIS-SFS) to the predicted SFS assuming a gamma DFE. The MIS-SFS data (gray) and expected SFS at the maximum-likelihood parameter estimates (red) are shown for each dataset.

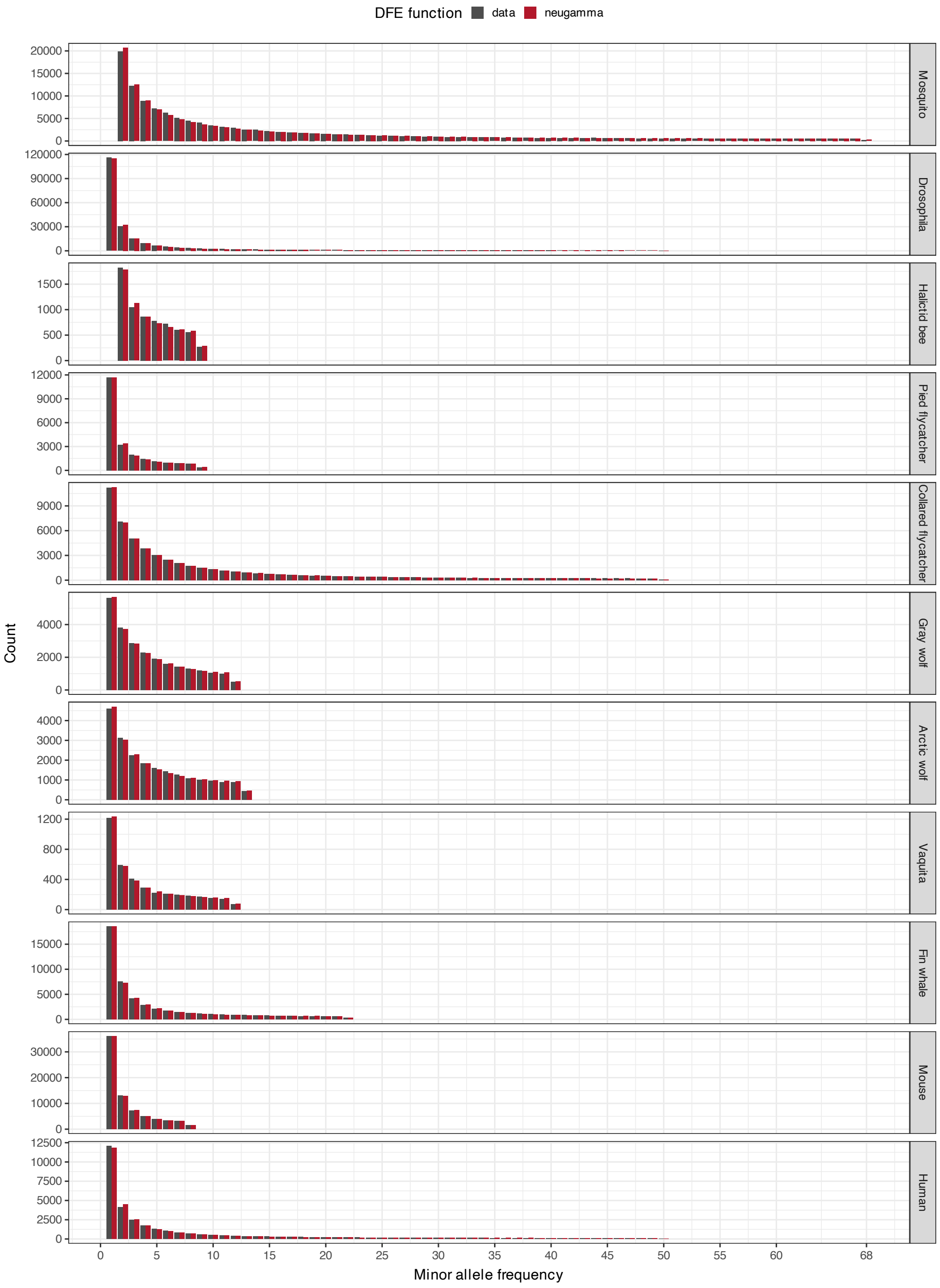


Figure S5: Comparison of the observed nonsynonymous SFS (MIS-SFS) to the predicted SFS assuming the DFE is a mixture of a gamma distribution and a neutral point mass (nuegamma DFE). The MIS-SFS data (gray) and expected SFS at the maximum-likelihood parameter estimates (red) are shown for each dataset.

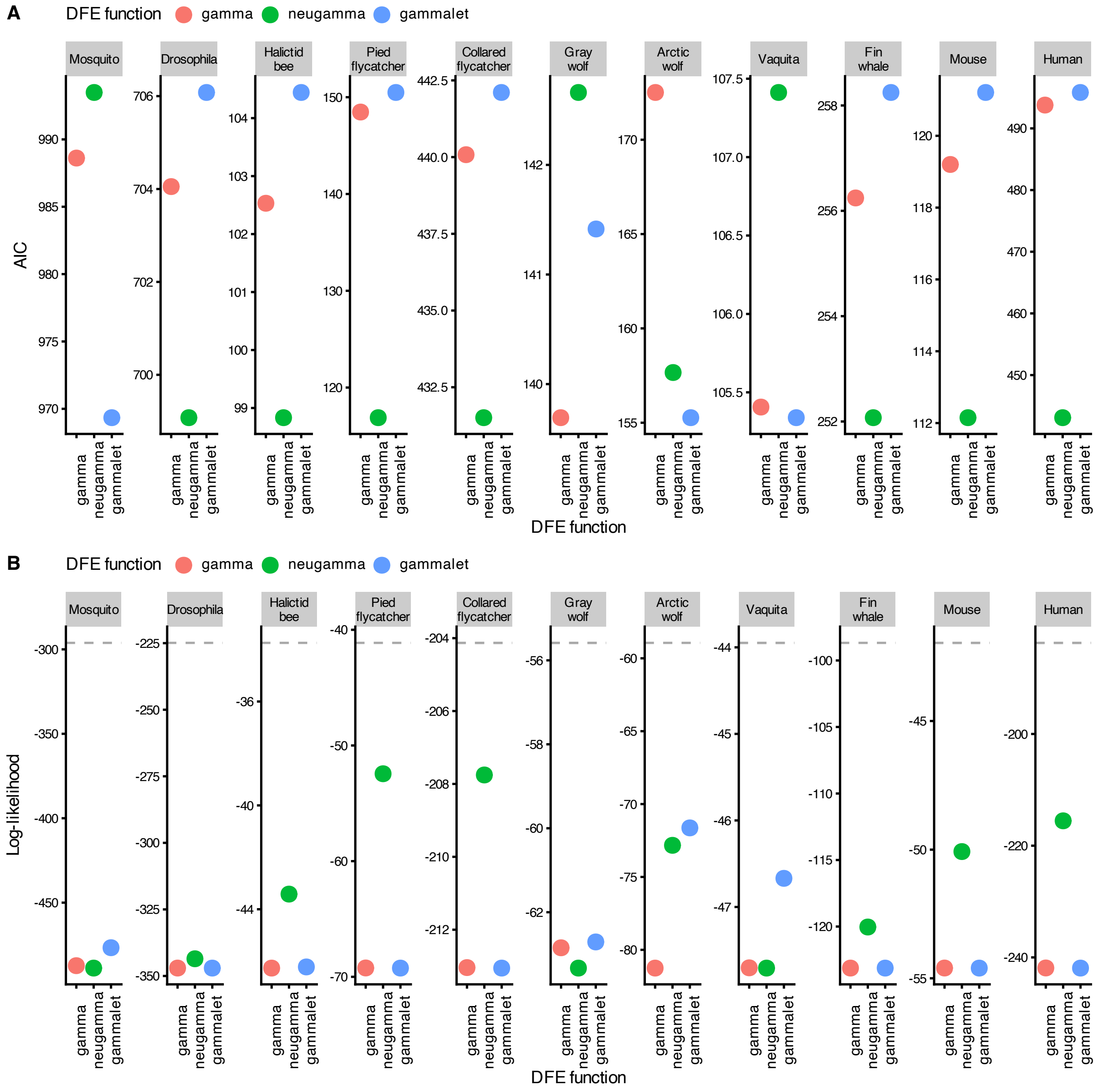


Figure S6: Comparisons on the (A) AIC, and (B) log-likelihoods of the DFE inference assuming that the DFE follows a gamma, neugamma, or gammalet distribution. The gray dashed line in (B) represents the maximum log-likelihood derived from the true data.

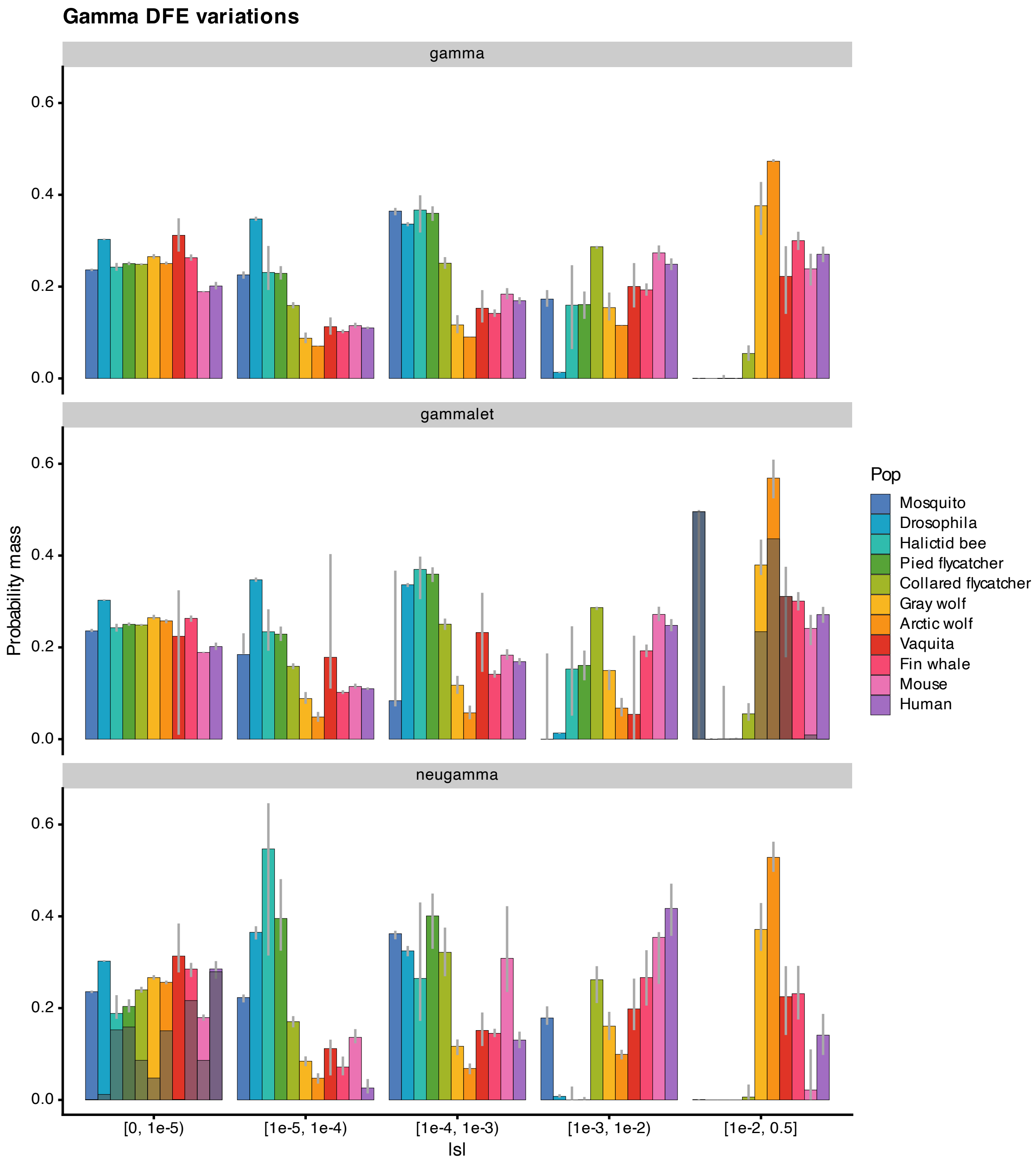


Figure S7: Proportions of mutations in various categories of $|s|$. From left to right, mutations range from (nearly) neutral ($-10^{-5} < s \leq 0$) to very strongly deleterious ($s \leq -0.01$). Different panels represent different assumptions for the functional forms of the DFE: gamma, neugamma, or gammalet. In the neugamma panel, the darker proportion at the left-most column ($-10^{-5} < s \leq 0$) represents the inferred point mass at neutrality. In the gammalet panel, the darker proportion at the right-most column ($s \leq -0.01$) represents the inferred point mass at lethality. Gray lines represent FIM-derived confidence interval.

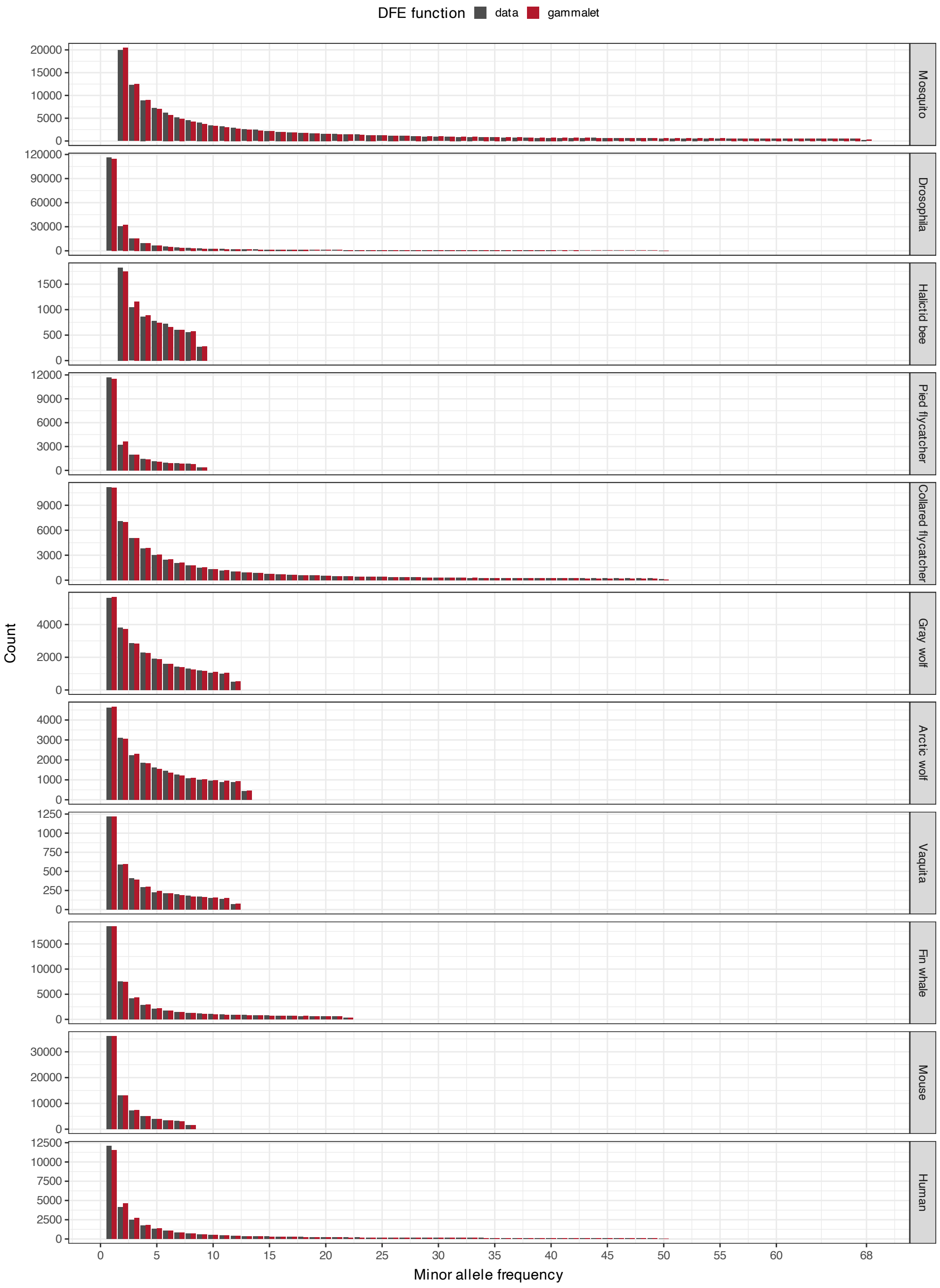


Figure S8: Comparison of the observed nonsynonymous SFS (MIS-SFS) to the predicted SFS assuming the DFE is a mixture of a gamma distribution and a lethal point mass (gammalet DFE). The MIS-SFS data (gray) and expected SFS at the maximum-likelihood parameter estimates (red) are shown for each dataset.

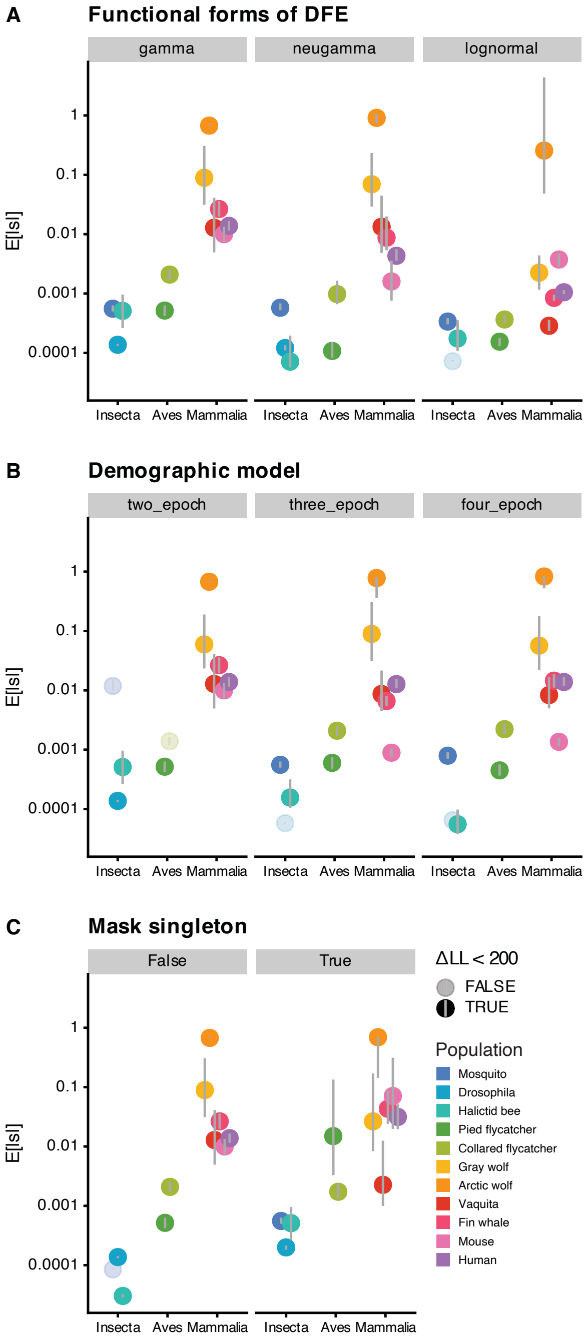


Figure S9: The DFE inference is mostly robust to potential confounding factors. The expected value of the selection coefficient ($E[|s|]$) for each species under different (A) functional forms of the DFE (B) demographic model assumptions and (C) singleton-masking treatments. Points with different colors represent different (sub)species. In DFE inferences that did not fit the data, defined as the log-likelihood difference between the data and model exceeds 200 ($\Delta LL < 200$ is FALSE), points are marked as translucent.

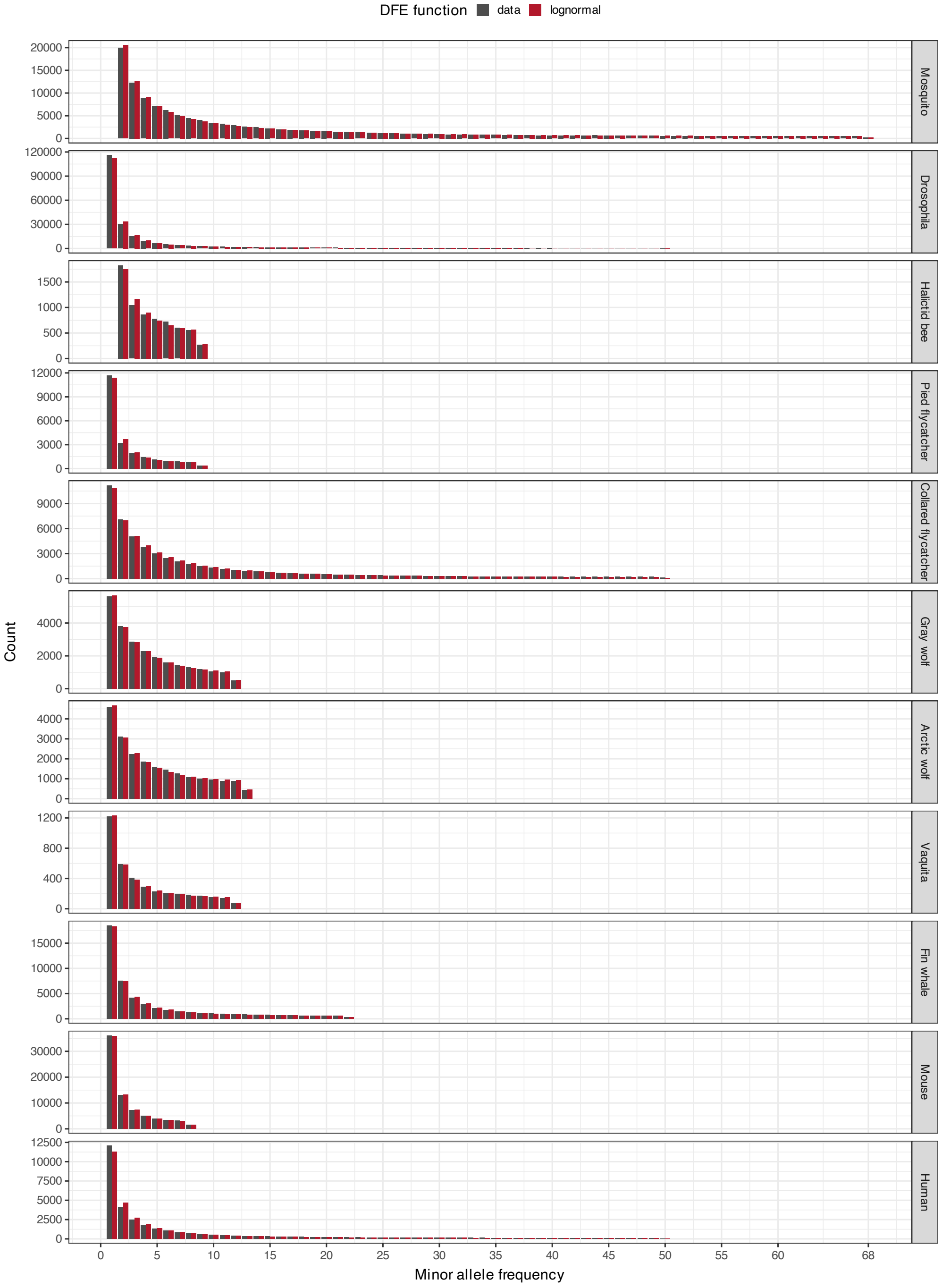


Figure S10: Comparison of the observed nonsynonymous SFS (MIS-SFS) to the predicted SFS assuming the DFE follows a lognormal distribution. The MIS-SFS data (gray) and expected SFS at the maximum-likelihood parameter estimates (red) are shown for each dataset.

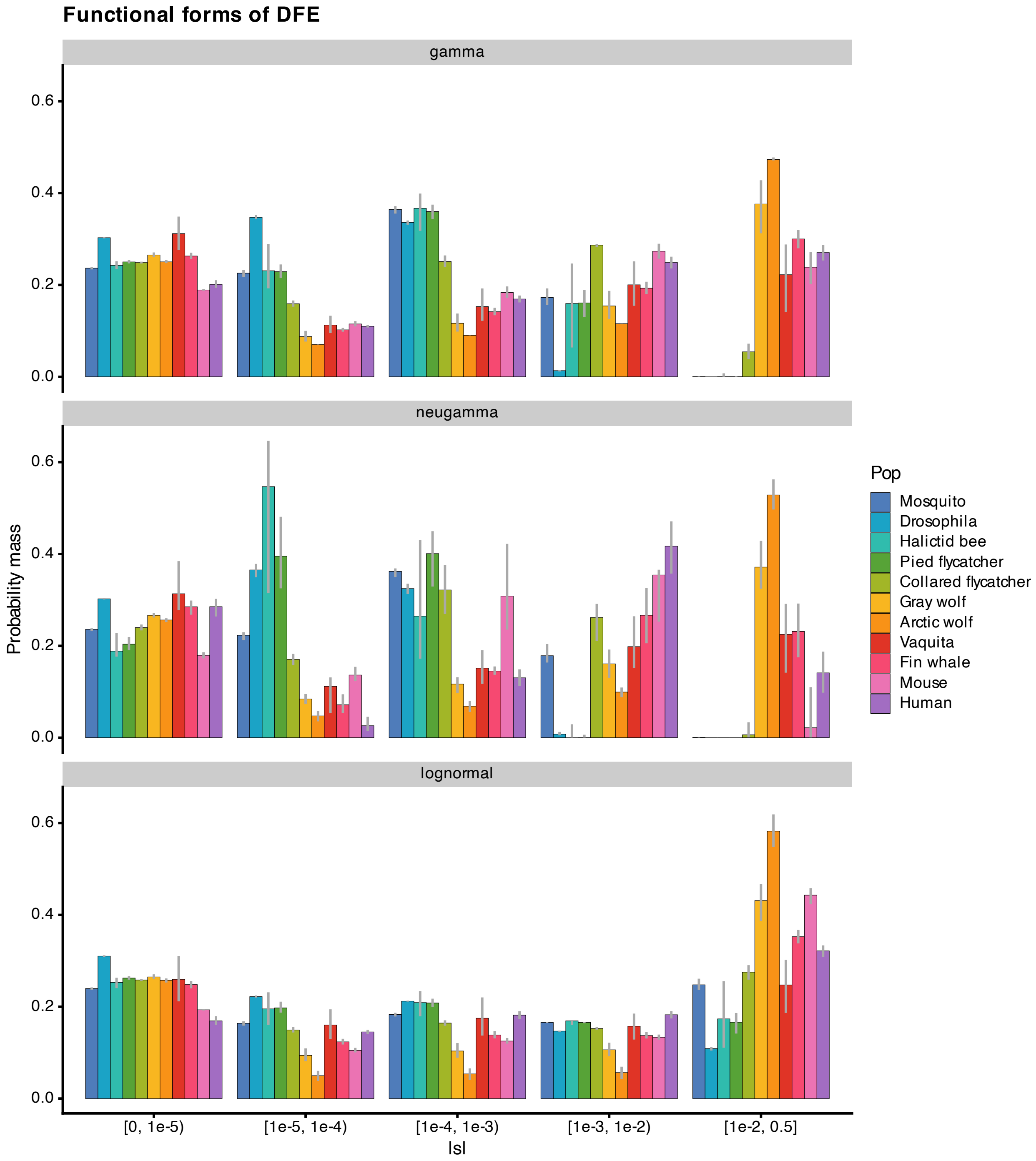


Figure S11: Proportions of mutations in various categories of $|s|$. From left to right, mutations range from (nearly) neutral ($-10^{-5} < s \leq 0$) to very strongly deleterious ($s \leq -0.01$). Different panels represent different assumptions for the functional forms of the DFE: gamma, neugamma, or lognormal. Gray lines represent FIM-derived confidence interval.

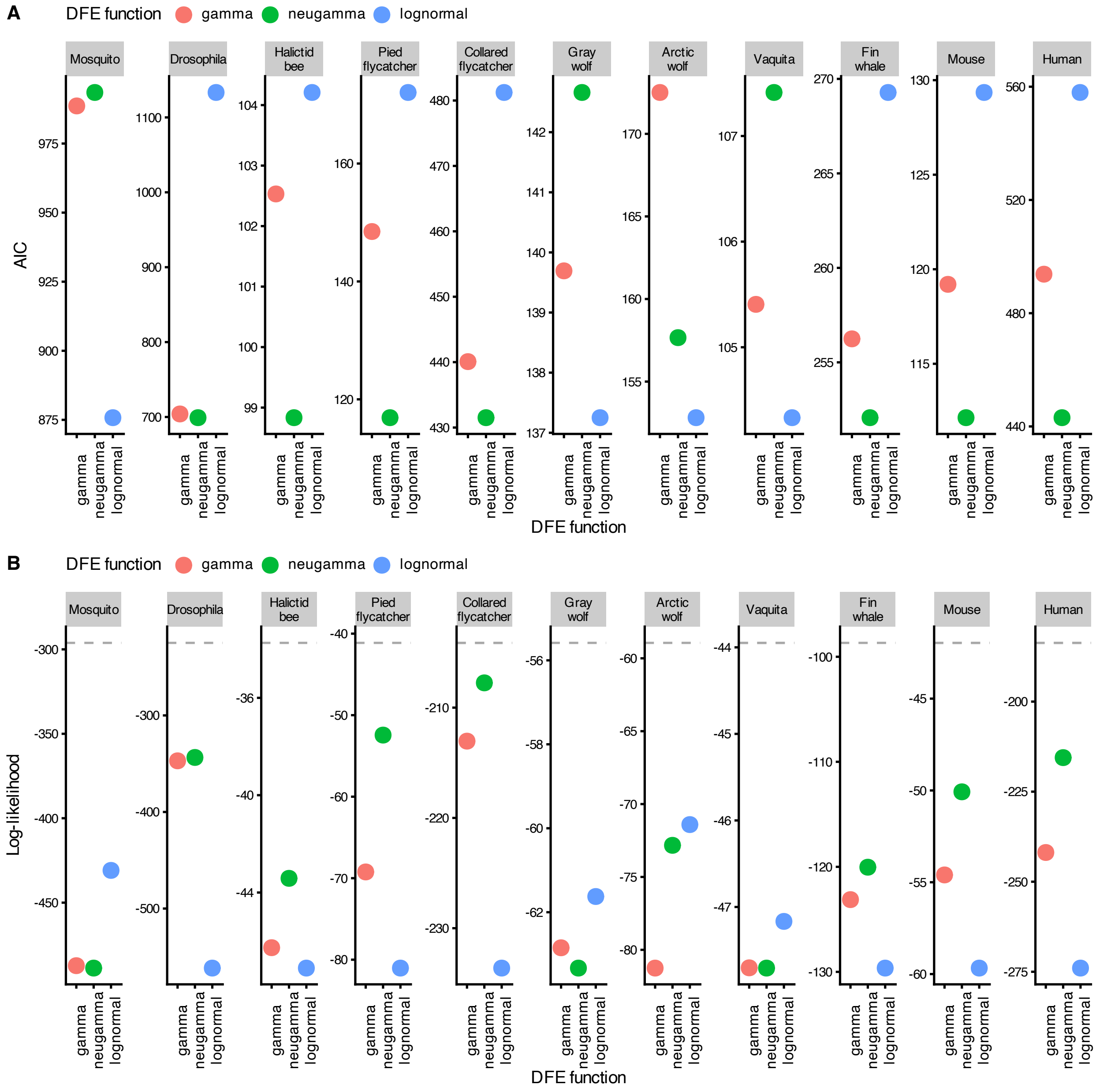

Figure S12: Comparisons on the (A) AIC, and (B) log-likelihoods of the DFE inference assuming that the DFE follows a gamma, neugamma, or lognormal distribution. The gray dashed line in (B) represents the maximum log-likelihood derived from the true data.

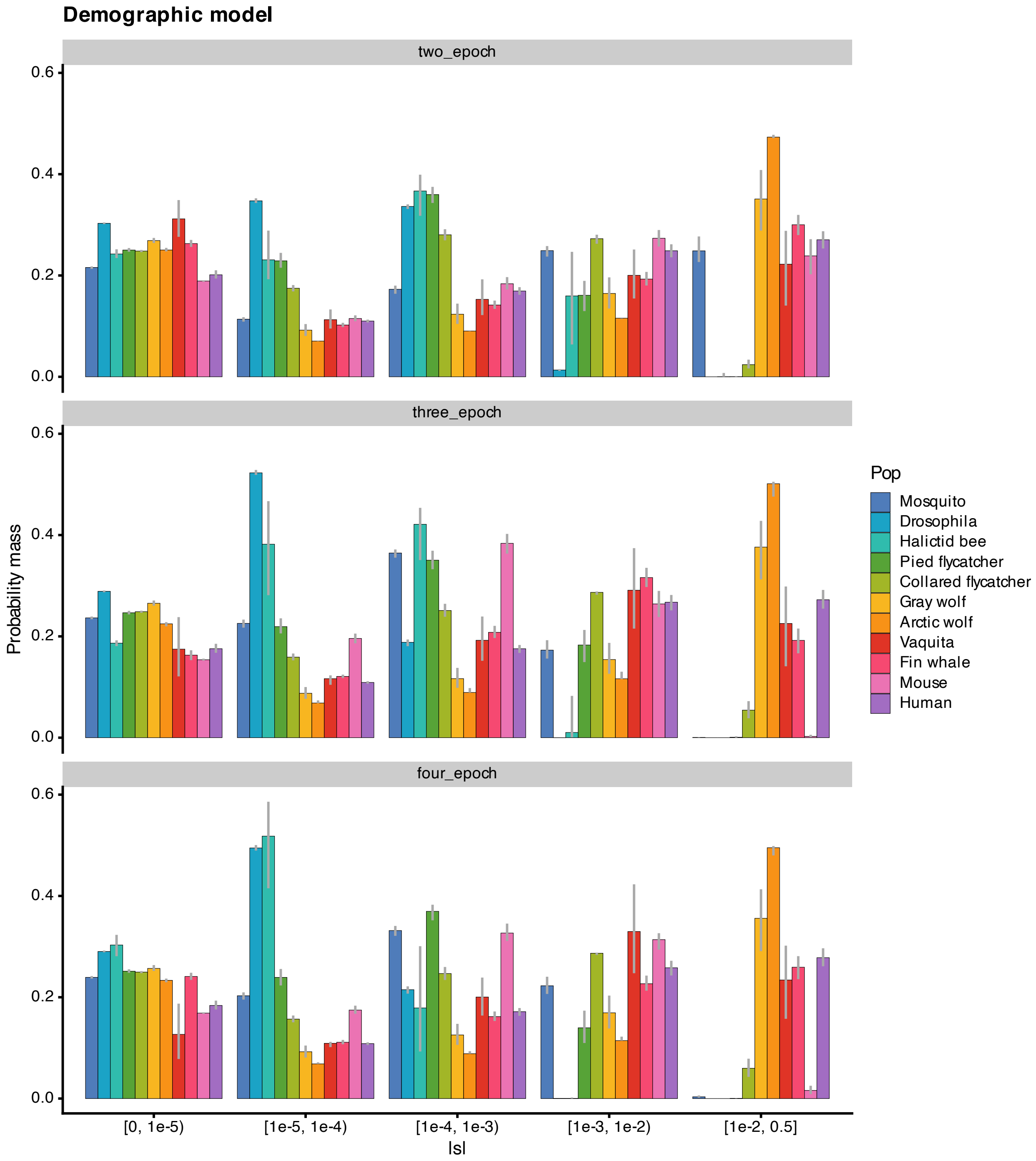


Figure S13: Proportions of mutations in various categories of $|s|$. From left to right, mutations range from (nearly) neutral ($-10^{-5} < s \leq 0$) to very strongly deleterious ($s \leq -0.01$). Assuming a gamma-distributed DFE, different panels represent different assumptions for the underlying demographic models of the population: two-, three-, and four-epoch. Gray lines represent FIM-derived confidence interval.

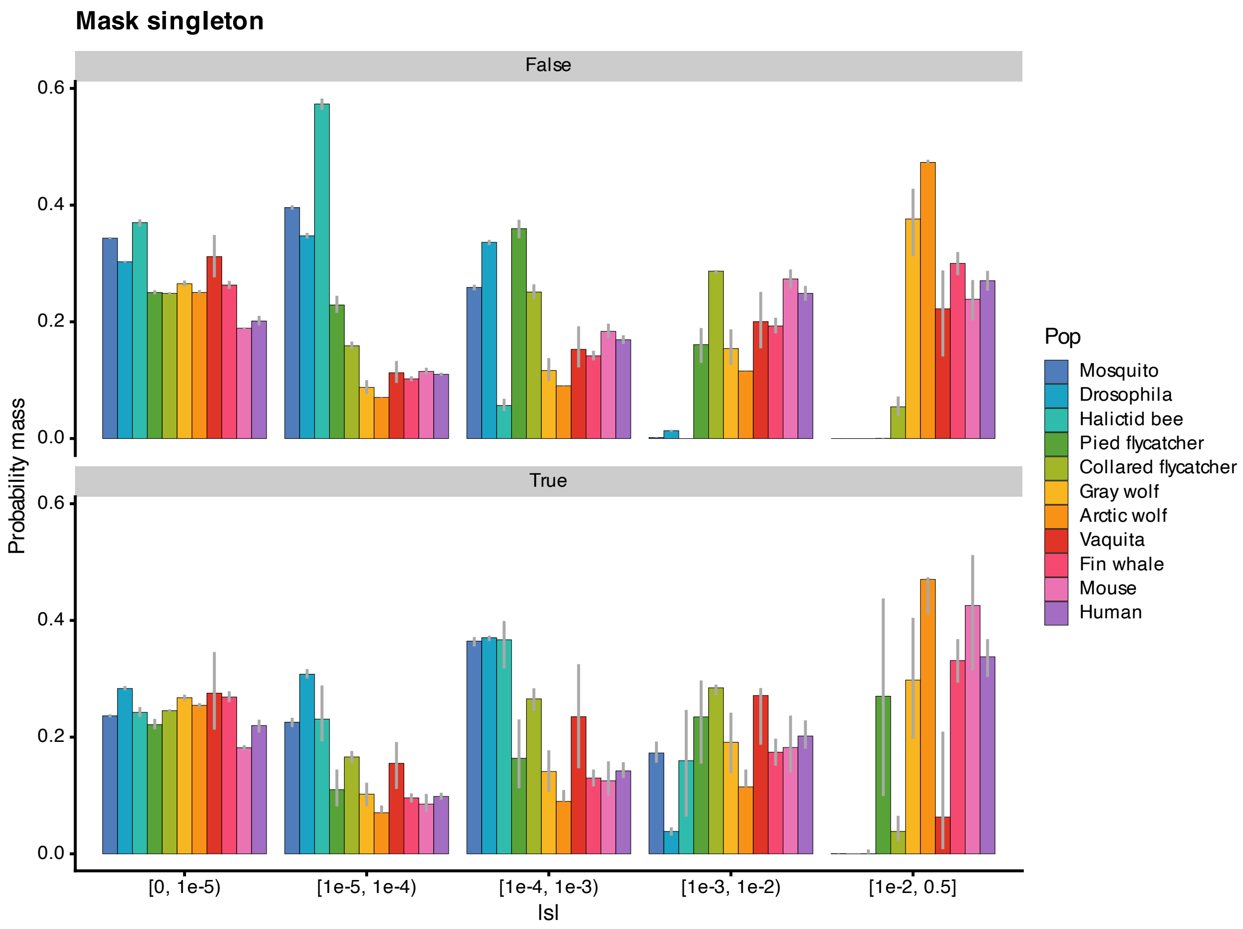


Figure S14: Proportions of mutations in various categories of $|s|$. From left to right, mutations range from (nearly) neutral ($-10^{-5} < s \leq 0$) to very strongly deleterious ($s \leq -0.01$). Assuming a gamma-distributed DFE, the top panel's inferred proportions are based on the full SFS and the bottom panel represents the inferred proportions of $|s|$ after masking singletons in the SFS. Gray lines represent FIM-derived confidence interval.

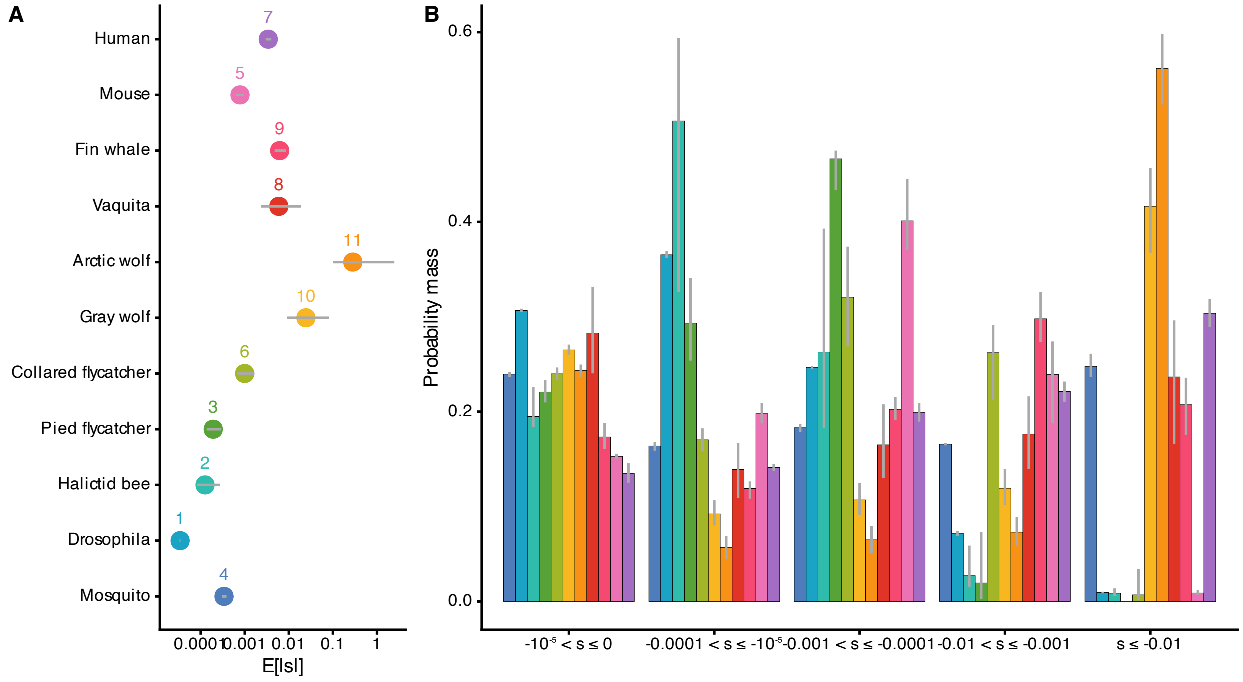


Figure S15: Model-averaged DFE estimates using the best singleton-masking treatments treatment per dataset. For mosquitos and halictid bees species, inference from the singleton-masked SFS are shown. For nine other species, results are from the full SFS. The estimates are weighted-averages of six DFE and demographic inference runs per species, combining outputs from gamma-, neugamma-, lognormal-distributed DFE given two-epoch or three-epoch demographic models by their AIC. (A) The model-averaged expected deleterious mutation effects ($E[|s|]$) for eleven (sub)species. Numbers show the ranks of $E[|s|]$. (B) The model-averaged proportions of mutations in various categories of $|s|$. From left to right, mutations range from (nearly) neutral ($-10^{-5} < s \leq 0$) to very strongly deleterious ($s < -0.01$). (A-B) Gray lines represent 95% confidence intervals.

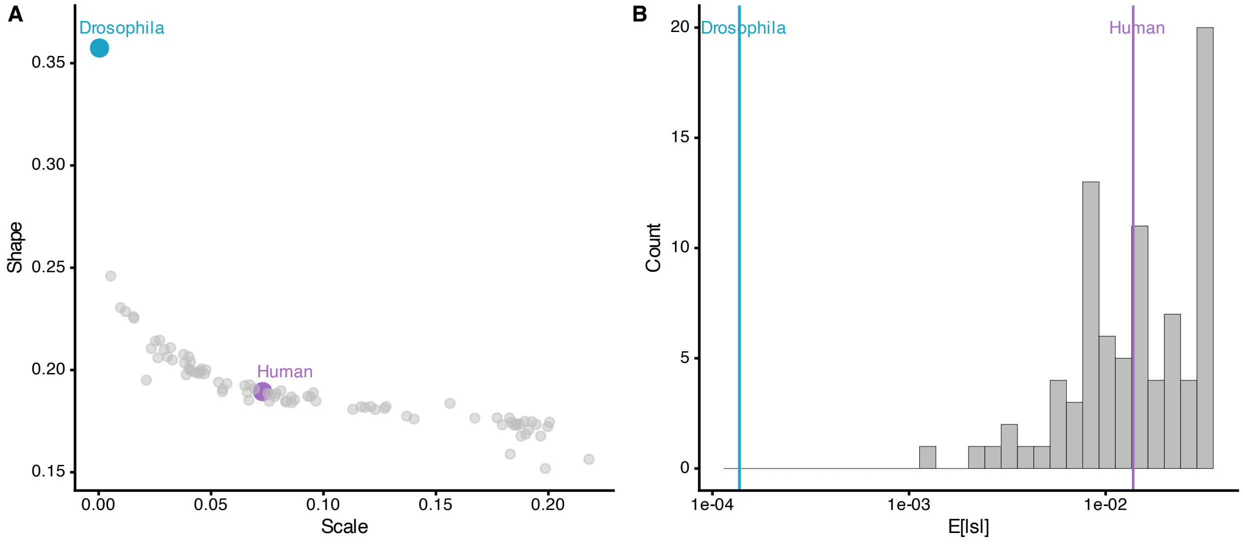


Figure S16: DFE inference from SLiM simulations using *Drosophila* demography and human DFE parameters. (A) The inferred gamma DFE shape and scale parameters in the 84 replicates (gray) are plotted against the human shape and scale parameters (purple points) and *Drosophila* shape and scale parameters (blue). The simulation was set with human DFE, and the inferred DFE is more similar to human than *Drosophila*, suggesting our DFE inference methods can detect strongly deleterious mutations even in large populations. (B) The histogram of the inferred $E[|s|]$ in the 84 simulation replicates (gray) is shown along with the human $E[|s|]$ and *Drosophila* $E[|s|]\dot{T}$he inferred $E[|s|]$ is more similar to human than *Drosophila*.

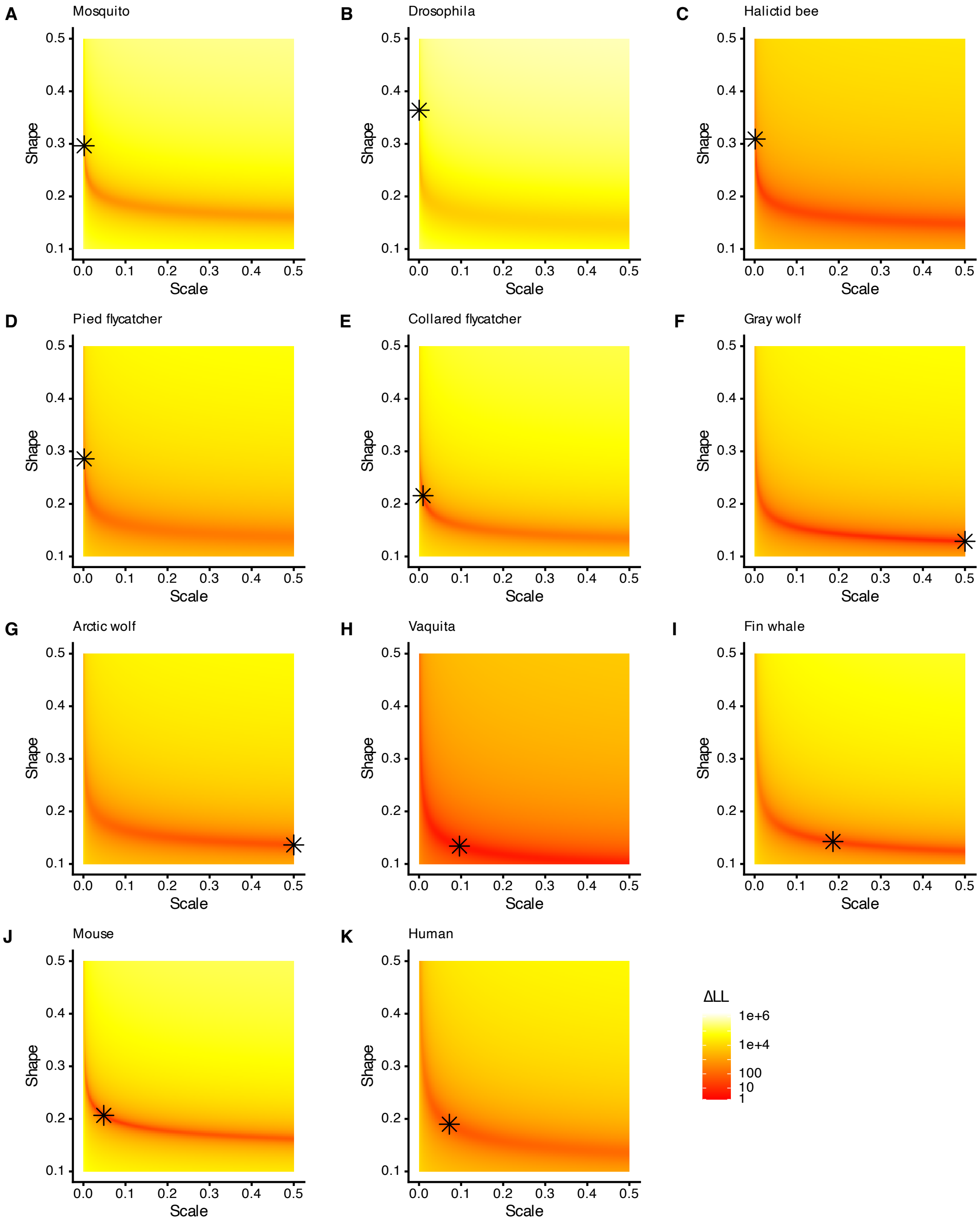


Figure S17: Full grid search results complementary to Fig. 3. The log-likelihood surface for the shape ($\alpha$) and scale ($\beta'$) parameters are shown for each species. Background colors from yellow to red indicate the differences in log-likelihood for given parameters to data. On each log-likelihood surface, the maximum likelihood estimate (MLE) derived for the respective species is overlaid as the asterisk.

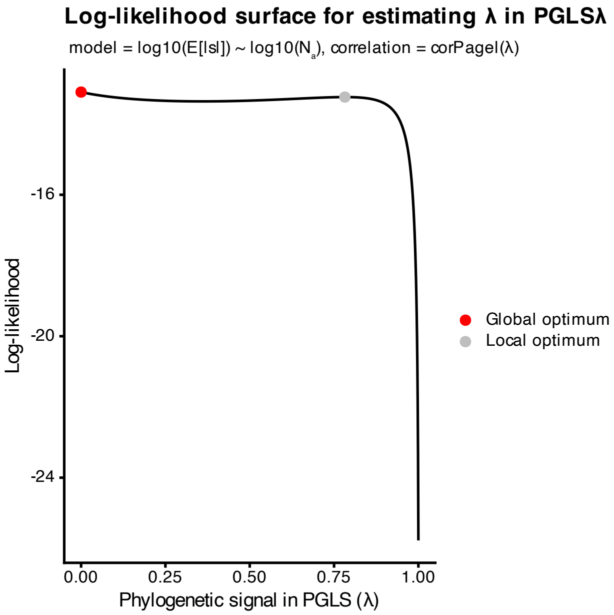


Figure S18: The log-likelihood surface for $\lambda$ estimation in the PGLS$_\lambda$ model of expected mutation effects ($E[|s|]$) and long-term population size ($N_a$). Both $E[|s|]$ and $N_a$ are log10-scaled to normalize the data. The log-likelihood surface exhibits a relatively flat profile between the global optimum at $\lambda = 0$ (red point, $LL_{\lambda=0}$ = -13.10) to the local optimum at $\lambda = 0.78$ (gray point, $LL_{\lambda=0.78}$ = -13.24). This flatness indicates comparable model performance across various $\lambda$ values, suggesting uncertainty regarding the independence of the correlation between $E[|s|]$ and $N_a$ from phylogenetic signal in the residual error.

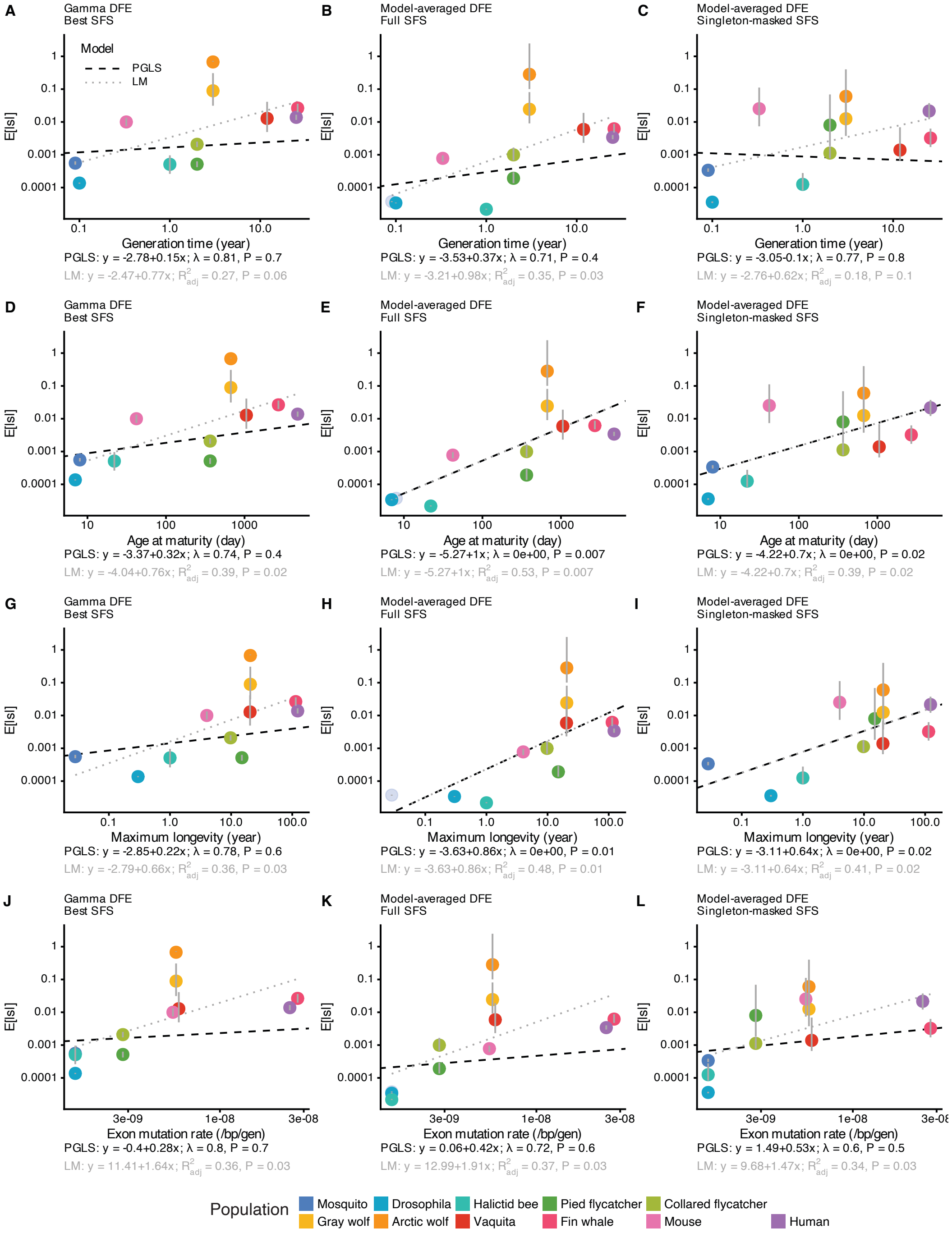


Figure S19: The phylogenetic generalized least square model with simultaneously inferred phylogenetic signal ($PGLS_\lambda$) was fitted with $E[|s|]$ derived from (left) gamma DFE, (middle) model-averaged DFE using full SFS, or (right) model-averaged DFE using singleton-masked SFS as the response variable. The explanatory variables are: (A-C) generation time, (D-F) age at maturity, (G-I) maximum longevity, and (J-L) exon mutation rates. The $PGLS_\lambda$ results are overlaid as a dashed black line, with equations, inferred phylogenetic signal ($\lambda$), and likelihood-ratio derived p-value annotated at the bottom. The linear regression results are overlaid as dotted gray line, with equations and p-values annotated at the bottom. All axes are in log10 scale.

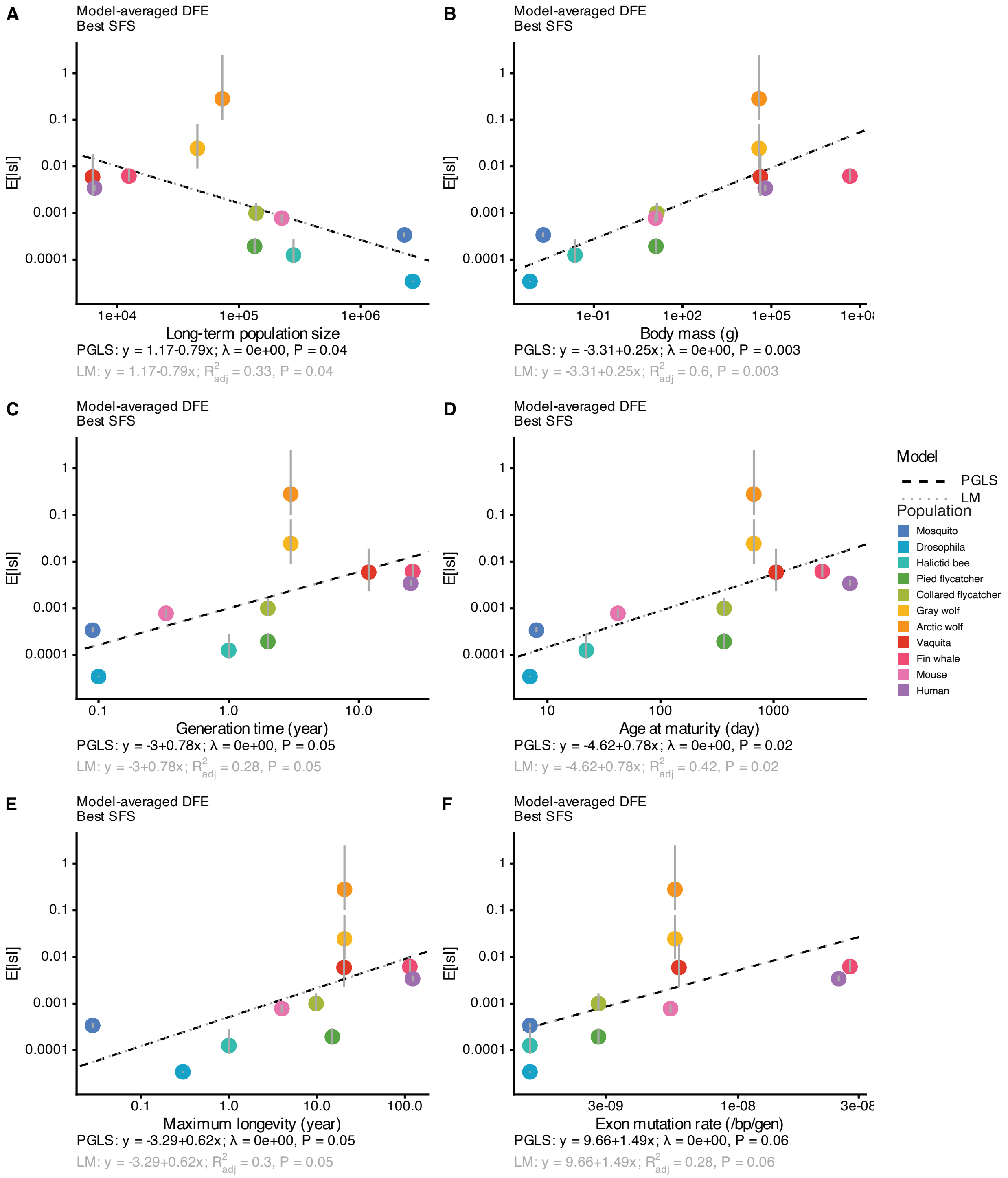


Figure S20: The phylogenetic generalized least square model with simultaneously inferred phylogenetic signal ($PGLS_\lambda$) was fitted with $E[|s|]$ derived from model-averaged DFE using the best singleton-masking treatments as the response variable. The $PGLS_\lambda$ results are overlaid as a dashed black line, with equations, inferred phylogenetic signal ($\lambda$), and likelihood-ratio derived p-value annotated at the bottom. The linear regression results are overlaid as dotted gray line, with equations and p-values annotated at the bottom. All axes are in log10 scale.

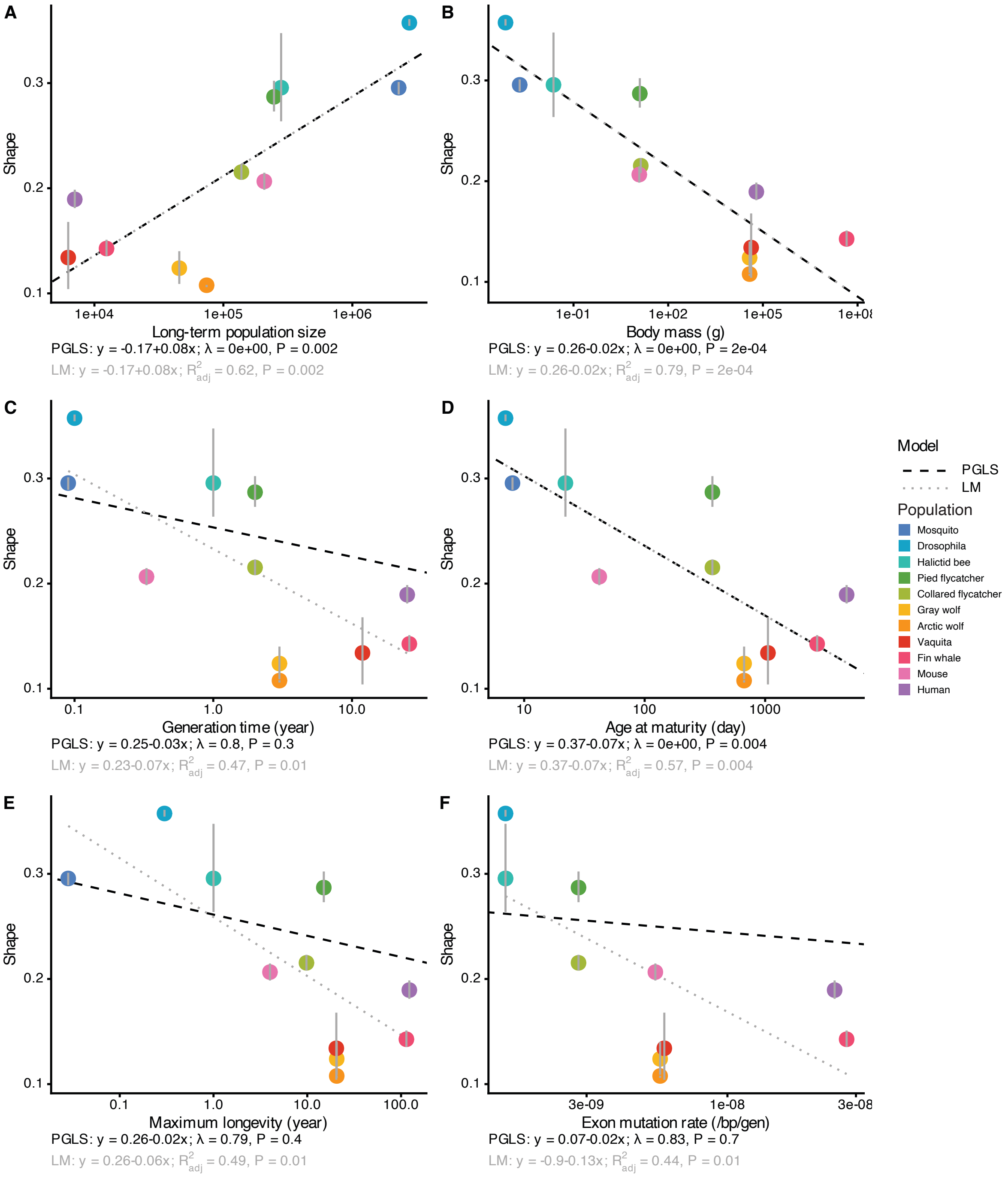


Figure S21: The phylogenetic generalized least square model with simultaneously inferred phylogenetic signal ($PGLS_\lambda$) was fitted for the inferred gamma DFE's shape ($\alpha$) parameter and (A) long-term population size, (B) body mass, (C) generation time, (D) age at maturity, (E) maximum longevity, and (F) exon mutation rates. The $PGLS_\lambda$ results are overlaid as a dashed black line, with equations, inferred phylogenetic signal ($\lambda$), and likelihood-ratio derived p-value annotated at the bottom. The linear regression results are overlaid as a dotted gray line, with equations and p-values annotated at the bottom. The x-axes are in log10 scale.

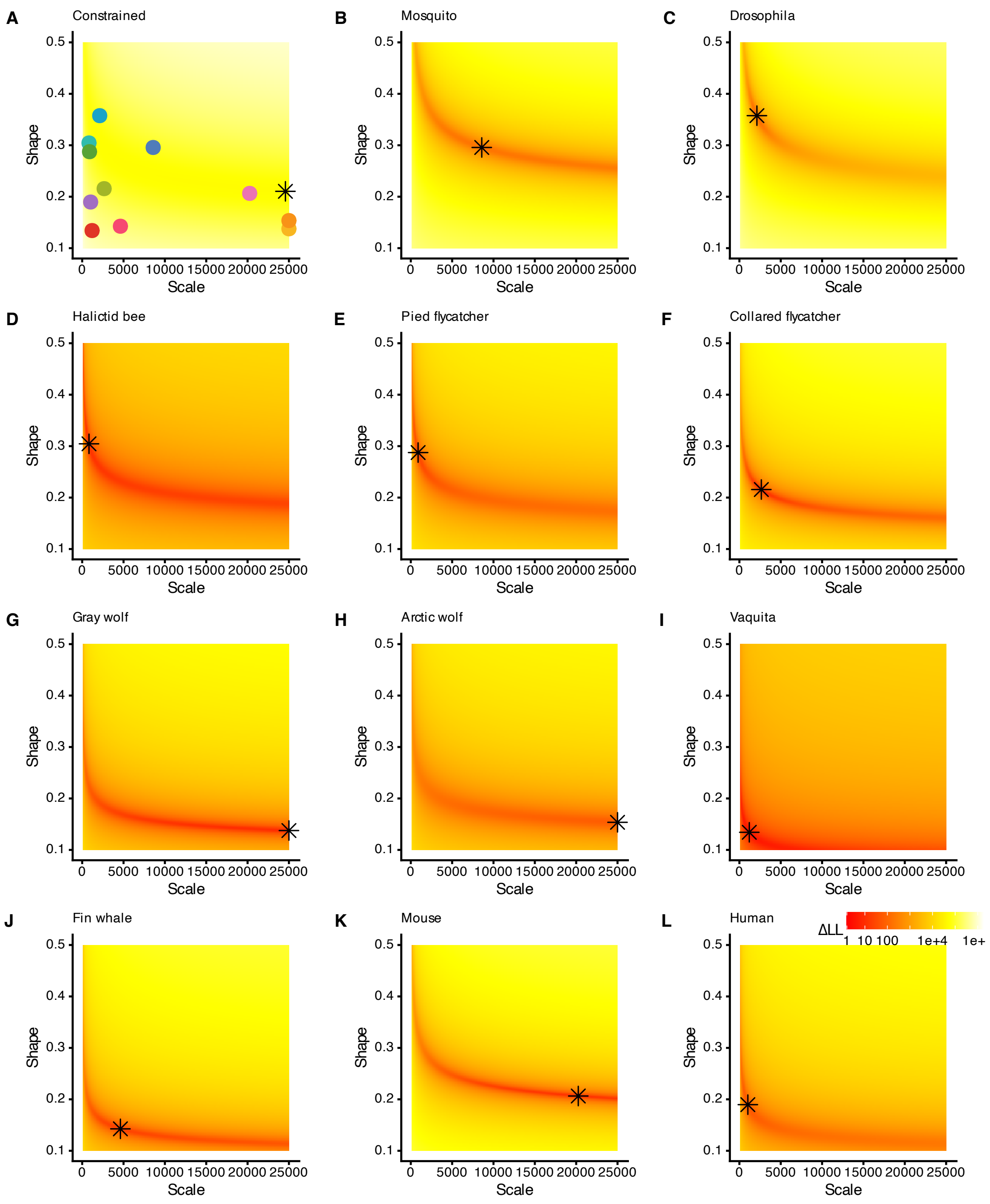


Figure S22: Full grid search results for the population-scaled DFE. The log-likelihood surface for the shape ($\alpha$) and population-scaled scale ($\beta$) parameters are shown (A) under the constrained model where all species have the same parameters, or (B-L) under alternative models allowing each species' parameters to vary across datasets. Background colors from yellow to red indicate the differences in log-likelihood for given parameters to data. The color schemes are consistent with Fig. 3. On each log-likelihood surface, the maximum likelihood estimate (MLE) derived for (A) the constrained model or (B-L) respective species is overlaid as the asterisk. In (A), the colored points show MLEs derived from each species' grid search.

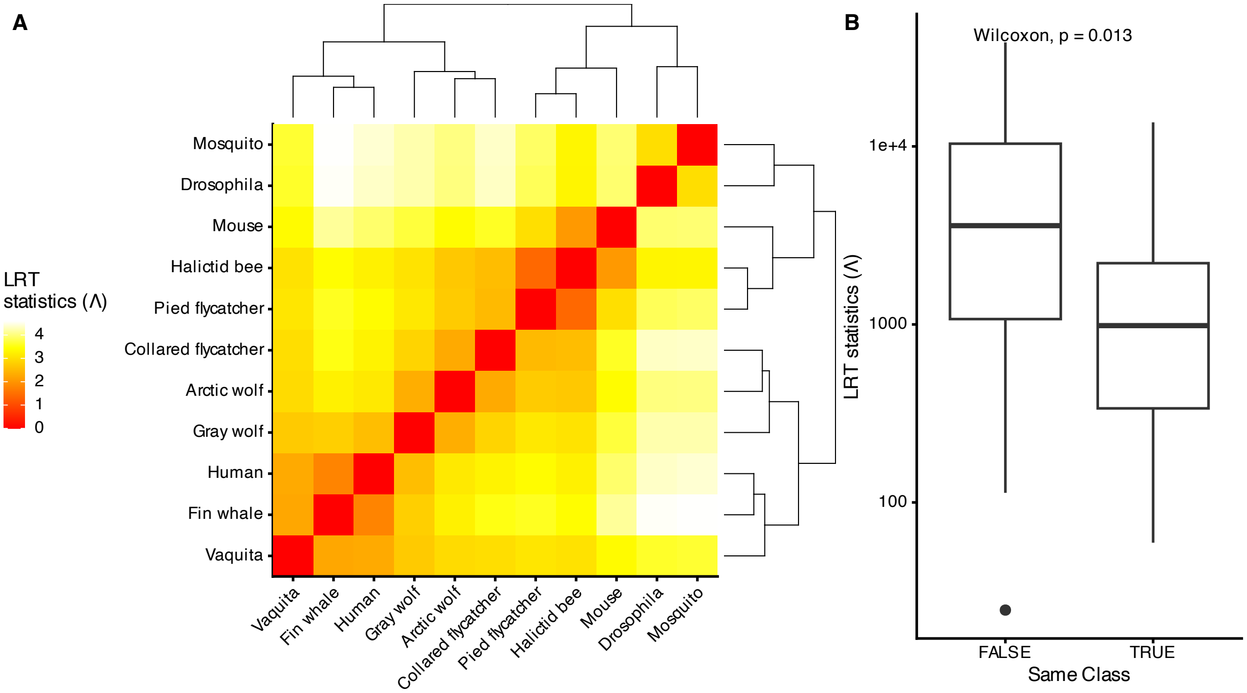


Figure S23: Pairwise likelihood ratio test (LRT) statistics for the population-scaled DFE. (A) Pairwise LRT statistics for each species pair are colored on a log10 scale ($\log_{10}(\Lambda)$) and hierarchically clustered. Darker cells represent more similar population-scaled DFE estimates in the species compared, such as fin whales and humans, whereas lighter cells represent more distinct DFEs, such as fin whales and *Drosophila*. The dendrogram derived from hierarchical clustering is annotated. (E) The boxplot of $\Lambda$ values, categorized by whether species pair belong to the same taxonomic Class. The y-axis is in log10 scale.

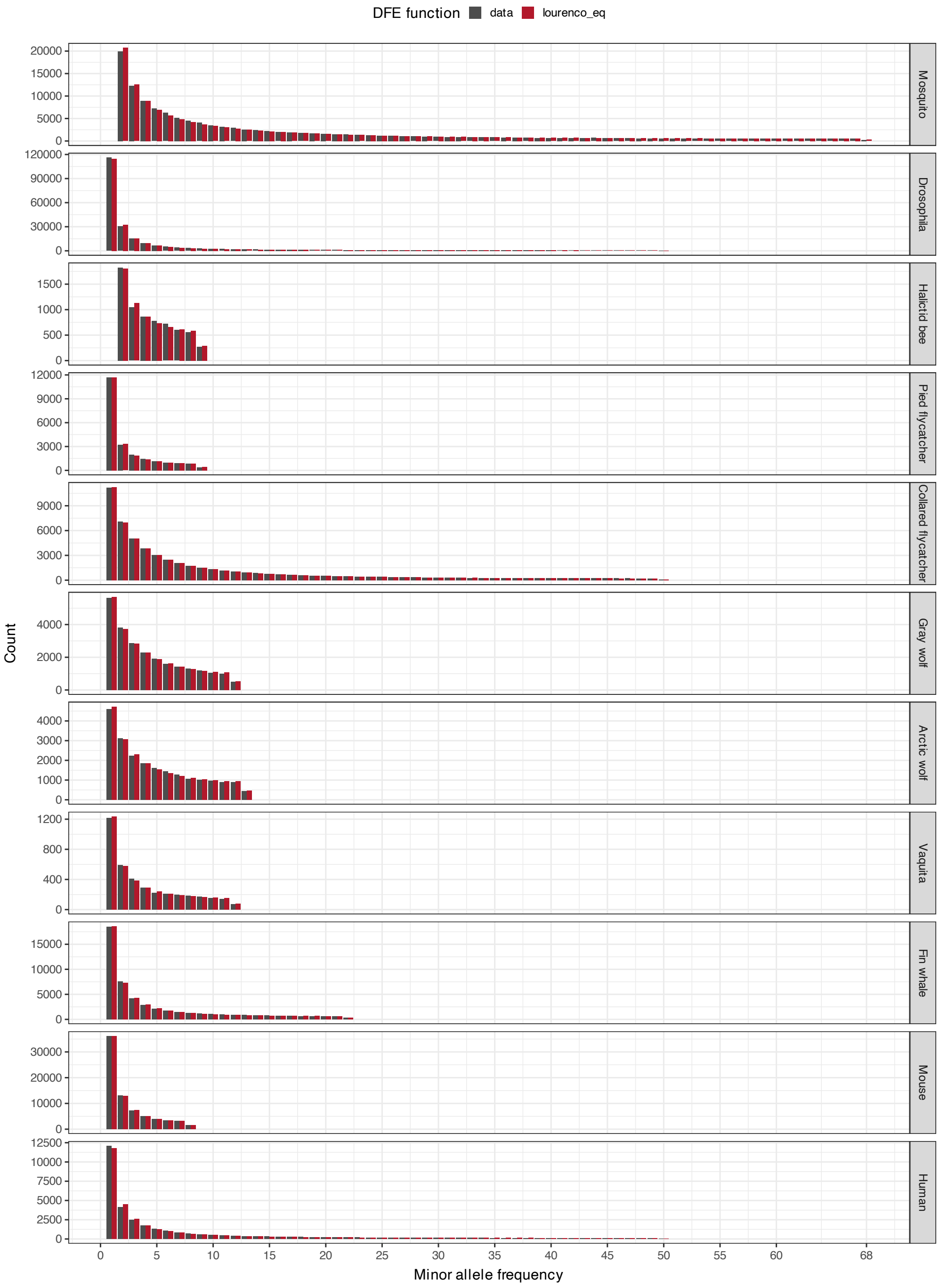


Figure S24: Comparison of the observed nonsynonymous SFS (MIS-SFS) to the predicted SFS assuming the DFE follows the Fisher's geometric model derived distribution [43]. The MIS-SFS data (gray) and expected SFS at the maximum-likelihood parameter estimates (red) are shown for each dataset.

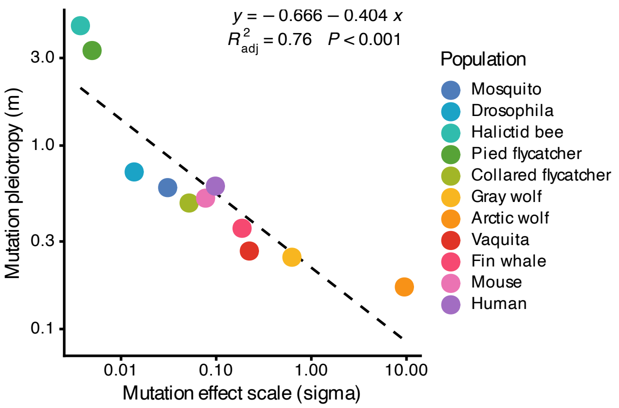


Figure S25: The inferred mutation pleiotropy ($m$) and mutation effect scale ($\sigma$) parameters in the FGM-derived DFE are negatively correlated. Both axes are in log10 scale. Points are colored by datasets.

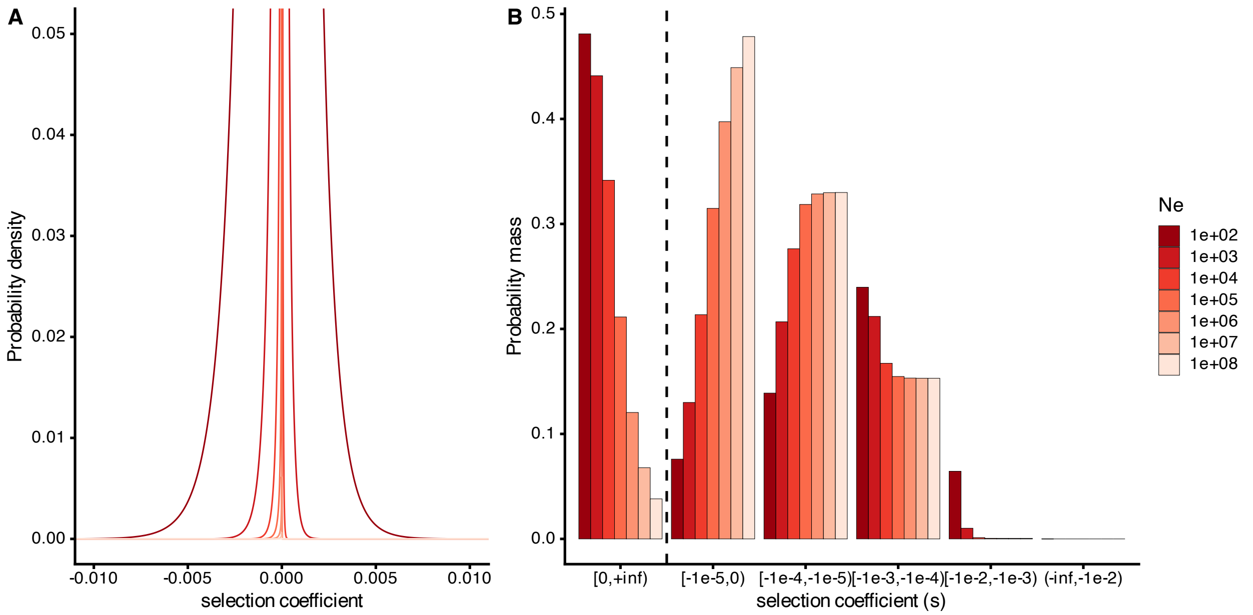


Figure S26: The FGM-derived DFE at mutation-selection-drift equilibrium becomes more dispersed as the long-term population size ($N_a$) declines. (A) The probability density function near $-0.01 < s < 0.01$ for $N_a$ ranging from 100 to $10^8$ (red to light-red). The mutation pleiotropy ($m$) is set at 0.5 and the mutation effect scale ($\sigma$) is set at 0.01. (B) The estimated proportion of each class of mutations with $N_a$ ranging from 100 to $10^8$. From left to right, mutations range from beneficial ($s > 0$), (nearly) neutral ($-10^{-5} < s \leq 0$) to very strongly deleterious ($s \leq -0.01$). Dashed line marks $s = 0$. As $N_a$ increases from 100 to $10^8$, the proportion of beneficial mutation decreases while the proportion of (nearly) neutral mutations increases.

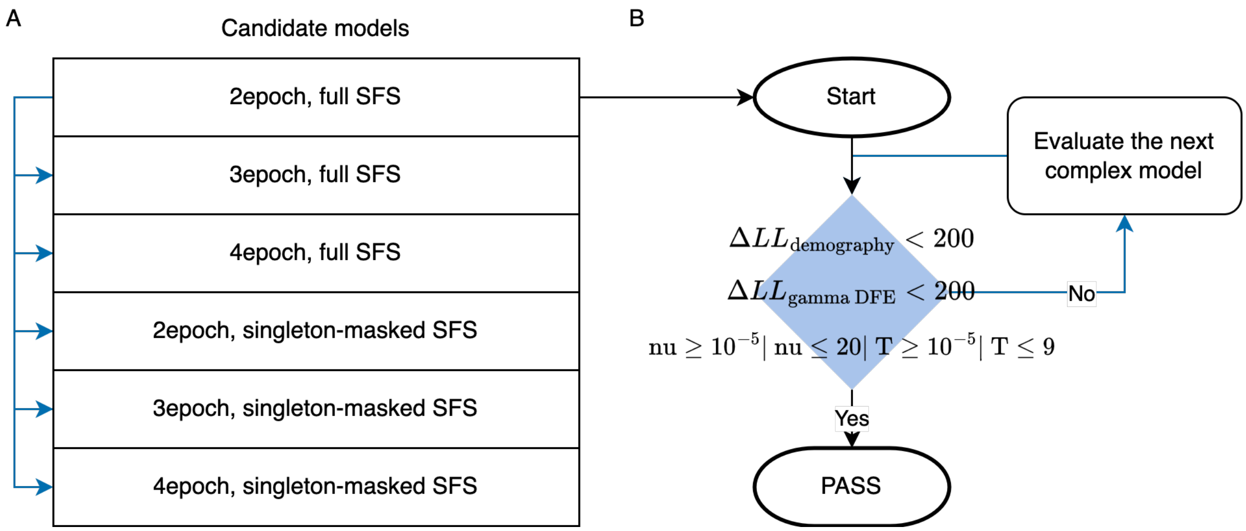


Figure S27: The step-wise demographic model selection procedure. (A) To maximize the data available and use the most parsimonious model whenever possible, at the start of model evaluation, the two-epoch demographic model based on unmasked SYN-SFS run was chosen for all datasets (top row). (B) This "2epoch, full SFS" run was chosen for a dataset (black workflow lines), if it met all three criteria listed in the blue diamond. 1. The log-likelihood differences between the data and model maximum likelihood $\Delta LL = LL_{\text{data}} - LL_{\text{model}}$ for demographic inference is less than 200. 2. The $\Delta LL$ for gamma DFE inference based on the current demographic model and singleton-masking treatment is less than 200. 3. The parameter estimates in the current demographic model are realistic (unrealistic parameters defined as nu $< 10^{-5}$ or nu $> 20$ or T $< 10^{-5}$ or T $> 9$). (A-B)Otherwise, we proceeded to evaluate the more complex run "3epoch, full SFS" with the same criteria (Blue workflow lines). If the "3epoch, full SFS" failed the criteria again, we proceeded to choose the next model listed in (A) iteratively. If all models based on the full (unmasked) SFS failed to provide a good fit (rows 1-3 in A), we proceeded to mask the singletons in the SFS (rows 4-6 in A).

### 6 Legends for Datasets S1 to S15

1. Polymorphism dataset and life history traits summary.
2. Synonymous and missense site-frequency spectra for each subspecies.
3. Demographic inference summary.
4. Demographic inference selection results.
5. Demographic inference comparisons with previous studies.
6. DFE inference summary assuming that the DFE follows a gamma distribution.
7. DFE inference summary assuming that the DFE follows a neugamma distribution.
8. DFE inference summary assuming that the DFE follows a gammalet distribution.
9. DFE inference summary assuming that the DFE follows a lognormal distribution.
10. DFE inference summary assuming that the DFE follows a gamma distribution under different demographic model assumptions.
11. DFE inference summary assuming that the DFE follows a gamma distribution under different singleton-masking treatments.
12. Model-averaged DFE inference summary.
13. Pagel's $\lambda$ analysis summary.
14. Phylogenetic generalized least square model summary.
15. DFE inference summary assuming that the DFE follows a Fisher's geometric model derived distribution (lourenco DFE).
16. DFE inference comparisons with previous studies.
